# Stem girth changes in response to soil water potential in lowland dipterocarp forest in Borneo: a phenomenological and individualistic time-series analysis

**DOI:** 10.1101/2020.07.28.222547

**Authors:** D. M. Newbery, M. Lingenfelder

## Abstract

Time series data offer a way of investigating the causes driving ecological processes as phenomena. To test for possible differences in water relations between species of different forest structural guilds at Danum (Sabah, NE Borneo), daily stem girth increments (*gthi*), of 18 trees across six species were regressed individually on soil moisture potential (SMP) and temperature (TEMP), accounting for temporal autocorrelation (in GLS-arima models), and compared between a wet and a dry period. The best-fitting significant variables were SMP the day before and TEMP the same day. The first resulted in a mix of positive and negative coefficients, the second largely positive ones. An adjustment for dry-period showers was applied. Interactions were stronger in dry than wet period. Negative relationships for overstorey trees can be interpreted in a reversed causal sense: fast transporting stems depleted soil water and lowered SMP. Positive relationships for understorey trees meant they took up most water at high SMP. The unexpected negative relationships for these small trees may have been due to their roots accessing deeper water supplies (if SMP was inversely related to that of the surface layer), and this was influenced by competition with larger neighbour trees. A tree-soil flux dynamics manifold may have been operating. Patterns of mean diurnal girth variation were more consistent among species, and time-series coefficients were negatively related to their maxima. Expected differences in response to SMP in the wet and dry periods did not clearly support a previous hypothesis differentiating drought and non-drought tolerant understorey guilds. Trees within species showed highly individual responses when tree size was standardized. Data on individual root systems and SMP at several depths are needed to get closer to the mechanisms that underlie the tree-soil water phenomena in these tropical forests. Neighborhood stochasticity importantly creates varying local environments experienced by individual trees.

## INTRODUCTION

The influence of climatic perturbation by drought on rainforest tree dynamics has received increasing attention during the last decades, especially with the last major El Niño Southern Oscillation (ENSO) events in 1982/83, 1997/98 and 2015/16. Field studies of the phenomenon were conducted in South-East Asia (Nakagawa et al. 2000, Harrison 2001, Aiba and Kitayama 2002, Potts 2003, Van Nieuwstadt and Sheil 2005, Vincent et al. 2009, Itoh et al. 2012). These forests experience a normally equable climate which is interrupted by infrequent strong dry spells. In Borneo ENSOs happen on average approximately once per 15 years (Walsh 1996, Walsh and Newbery 1999). Extensive studies have also been conducted in the Amazonia on the effects of ENSO related droughts (Phillips et al. 2009, Lewis et al. 2011), but the events are of a rather different nature as they come in addition to a regular intense dry season in that tropical region. Such climate perturbations in the long run form a process called environmental stochasticity.

Long-term research plots at Danum Valley in a primary lowland dipterocarp forest of Sabah, NE Borneo, have been maintained since 1985 in order to follow how the forest changed but also the effects of tropical climatic variability. They have been enumerated five times, in 1986, 1996, 2001, 2007 and 2015, and the forest dynamics recorded and analysed (Newbery et al. 1992, Newbery et al. 1996, Newbery et al. 1999, Newbery and Lingenfelder 2004, Lingenfelder and Newbery 2009, Newbery and Lingenfelder 2009, Newbery et al. 2011). In the first half of 1998 the forest at Danum was affected by unusually dry conditions associated with the 1997/98 ENSO, one of the strongest events on record for that part of the island. The usually aseasonal and continuously very wet climate (monthly rainfall 236 mm, (Newbery et al. 2011) was broken by a drought across almost 2 months between March and April 1998 with a 30-d running total of < 100 mm. Around the event the Oceanic Nino Index (ONI) was > 2 for 5 months (max. 2.4) (CPC 2020). An additional census on a subset of parts of the plots in 1999 revealed that many trees were strongly affected by the lack of rainfall with especially decreases in stem growth rate (Newbery and Lingenfelder 2004, Newbery et al. 2011).

The tree species in the ecosystem at Danum showed some evidence of adaptation to the moderate disturbance regime of occasionally severe droughts (Newbery et al. 1999, Walsh and Newbery 1999, Gibbons and Newbery 2003, Newbery and Lingenfelder 2004). The understorey is thought to play a key role in maintaining long-term structural stability via a feedback process in which species of the more numerous understorey species protect the smaller trees of overstorey ones, by keeping the forest microclimate favourable to them under the drought conditions (Newbery et al. 1999, Newbery et al. 2011, Newbery and Lingenfelder 2017). This model involves a mix of drought- and shade-tolerance among the understorey tree species leading to their different spatial distributions on the local topographic gradient (Newbery et al. 1996). The model does not necessarily imply any trade-offs in traits; it does not consider the growth and survival of seedlings or saplings. Trees of the understorey species, those of ≥ 10 cm stem girth, whose growth was advanced in dry periods are hypothesized to have different ecophysiological responses to limiting water availability, and to increased light levels, from trees of understorey species that were not advanced and from those small trees of overstorey species.

Moving closer to an understanding of the mechanisms that cause differences in effects of water limitation on tree growth across the species is not afforded by standard plot census monitoring with measurement intervals of 5-8 years. The clear 1996-1998 differences had narrowed and faded by 2001: the ‘memory effect’ was judged at 3-4 years (Newbery et al. 2011, Newbery and Lingenfelder 2017). A finer temporal resolution of stem size increment was required for a new period of dryness, preferably with a prior wet one for comparison. Unfortunately, against reasonable statistical expectation, another drought as severe as the 1997-1998 one did not occur by 2011. One of similar magnitude to 1997-98 did eventually occur though within a 6-month period of 2015-2016 (ONI > 2, maximum 2.6). In 2007 the decision had to be taken to start daily recording of tree girth in the hope of catching one or more dry periods which would demonstrate effects of limitation of tree water availability. A moderate event in 2009-2010 did, however, appear, and this allowed the analysis presented in this paper. Such considerations illustrate the precarious nature of field work involving environmental stochasticity operating at a scale of 1-2 decades. To have repeated events, i.e. a statistical sample over time, a century of recording might be just sufficient. The best prior indicators of a coming strong ENSO are at best 6 months ahead (Chen et al. 2004), and even that would not allow setting up a wet-period comparison beforehand.

Tree water relations can be explained to a large extent on the basis of physical laws (Monteith and Unsworth 1990, Jones 1992). How plants cope with limiting water supplies is today generally well understood, and this can be achieved with different mechanisms expressed collectively in the different species, in terms of their morphology and physiology, as traits (Zimmermann 1983, Tyree and Ewers 1991). Tree species with different ecologies are expected to show traits which are likely adaptive (Lambers et al. 1998). These traits, often quite complex though, may not be fully reducible to the detailed mechanisms but in their functioning they should show differences which are expressed as responses at an intermediate or phenomenological level. One example is how stem girth changes in time as water (sap) fluxes through its active tissues and, at the same time, the stem grows incrementally. How this girth increment changes with respect to driving environmental factors may give an insight as to how different tree species, or guilds of species when their ecologies and traits are similar, are adapted to variation in the demands and availability of water supply.

Several studies have demonstrated that tree girth fluctuation is closely correlated with water flux in the active xylem (e.g., (Kozlowski and Winget 1964, Herzog et al. 1995, Irvine and Grace 1997, Peramaki et al. 2001, Zweifel et al. 2006, Sevanto et al. 2008). Growth is positively dependent on water supply – drought conditions tend to limit it. So in itself growth may not be easily separated from changes induced by water flux, though being only positive it will often be much smaller than the positive-negative flux on a daily basis. Growth may be expected to be non-linearly related, and temporarily lagged, to tree water status because it depends on cell metabolism.

Previous research on growth patterns of individual trees in tropical forests has shown the use of mechanical band dendrometers to be straightforward and generally reliable over time periods of months to years when they are used carefully (Pelissier and Pascal 2000, Sheil 2003, Stahl et al. 2010). More precise measurements of tree stem expansion and contraction though need electronic band dendrometers in order to record at a finer within- and between-day resolution (Drew and Downes 2009). Such instruments, with their electronic loggers, require intensive maintenance in the field under demanding humid tropical forest conditions. The alternative electronic point dendrometers (Zweifel et al. 2006) require either mountings bolted into the wood, possibly causing damage to the stem or unnatural growth around the bolts, and outer bark removal: or an invar steel frame with screws that can be difficult to operate on wet bark surfaces (Sevanto et al. 2008). Furthermore, point dendrometers only measure radial change at one point on the circumference of a stem. There appears to be no evidence that point dendrometers do work effectively on trees in tropical everwet forests.

Band dendrometers have the potential disadvantage that temporary lowering of air humidity in wet periods or occasional short rain showers in dry periods may lead to the bark under the bands of some trees to respectively slightly shrink or expand, and therefore influence readings which are intended to reflect stem-wood girth changes (Stahl et al. 2010, Herrmann et al. 2016, Mencuccini et al. 2017, Oberhuber et al. 2020, Ilek et al. 2021). This can happen due to the hydroscopic nature of bark. This problem is dealt with in some detail in the present paper.

The present study followed daily stem size variation over time for six tree species at Danum (three under- and three overstorey), in a wet and a dry period, and related relative diurnal (= diel) changes, and daily increment time-series, to simultaneous recordings of soil moisture potential and ambient temperature. It is not a study of stem growth as such, in fact, growth trends are removed. The question was whether species in the proposed drought-tolerant and -intolerant guilds showed different responses, in the directions predicted, and whether these differences were strong enough to support the suggested over-understorey dynamics model. The following hypotheses and expectations were tested: (i) The rate of increase in daily girth increment with increasing soil moisture potential will be less steep for trees of understorey species per se than small trees of overstorey species. (ii) As soil moisture potential decreases, drought-tolerant understorey species will have smaller reductions in daily girth increment than drought-intolerant ones. (iii) Drought-tolerant understorey species should be less affected in terms of daily girth increment in dry compared to wet periods than drought-intolerant understorey ones and small trees of overstorey species. It is further possible that there is both a gradient in drought tolerance across understorey species, or within species an appreciable variation between individuals with respect to water stress that masks species differences.

## METHODS AND MATERIALS

### Study site

This research was conducted in permanent forest dynamics plots within the Danum Valley Conservation Area, in Sabah, Malaysia. The site is situated in lowland dipterocarp forest, c. 70 km east of the town of Lahad Datu on north-eastern Borneo. The climate is ever-wet and aseasonal, with on average 2881 mm/yr rainfall (1985 – 2011; www.searrp.org).

During the study period 6 March 2007 to 4 December 2010, mean (± SE) daily rainfall was 8.110 ± 0.37 mm (*n* = 1370), mean dry bulb temperature at 14:00 h was 30.250 ± 0.068 °C (*n* = 991), and mean relative humidity at 14:00 h was 79.65 ± 0.28% (*n* = 983), at the Danum Valley Field Centre (DVFC), 0.8 km west of the site.

Two main 4-ha plots had been fully enumerated in 1986, 1996, 2001 and 2007, for all trees with stem girth (*gbh*) ≥ 10 cm (or 3.1 cm dbh, diameter at breast height). The plot SW corners lay at 208 and 220 m a.s.l., and each had a topographic gradient, lower-slope-to-ridge of ca. 35 m change in elevation. More details on the site and the plots can be found in (Newbery et al. 1992, Newbery et al. 1996, Lingenfelder and Newbery 2009).

To find trees suitable for the dendrometers, data from the 2001 enumeration were used. Six common species that showed responses to the 1998 drought (three understorey, and three overstorey, Table 1), were sought from both ridge and lower slope locations (Newbery et al. 1996, Gibbons and Newbery 2003) in the main plots. Trees ≥ 10 cm diameter (*dbh*), or 31.42 cm girth (*gbh*) with intact stems and crowns were selected although some did have slightly smaller *dbh*-values. For ease of direct reference to previous papers for the site, *gbh* rather than *dbh* will be used for stem size.

**Table 1.** Species and numbers of trees selected for the electronic dendroband study at Danum (eDDS), and their distribution with topography.

| Species | Family<sup>1</sup> | Storey<sup>2</sup> | number |  |  |
| --- | --- | --- | --- | --- | --- |
|  |  |  | lower slope | ridge | all |
| *Dimorphocalyx muricatus* (Hook. f.) Airy Shaw | Euph | U | 1 | 3 | 4 |
| *Lophopetalum beccarianum* Pierre | Cela | U | 1 | 1 | 2 |
| *Mallotus wrayi* King ex Hook. f. | Euph | U | 3 | 1 | 4 |
| *Parashorea malaanonan* Merr. | Dipt | O | 1 | 2 | 3 |
| *Shorea fallax* Meijer | Dipt | O | 3 | 1 | 4 |
| *Shorea parvifolia* Dyer | Dipt | O | 1 | 2 | 3 |
| Totals |  |  | 10 | 10 | 20 |

<sup>1</sup> Dipt: Dipterocarpaceae; Euph: Euphorbiaceae; Cela: Celastraceae
<sup>2</sup>: O: overstorey species; U: understorey species (OUI)

### Selection of tree species

Within a list of 34 most abundant species (≥ 100 trees; (Newbery and Lingenfelder 2009) four were canopy-emergent species in the Dipterocarpaceae, the dominant, defining family in terms of basal area and the second most abundant in terms of stem density at Danum (Newbery et al. 1992, Newbery et al. 1996). Three of these dipterocarps showed strong responses to the 1998-drought: *Shorea parvifolia* decreased in relative growth rate (*rgr*) by *c*. 14% after the drought, but still had the highest *rgr* of all of the most abundant species; *Shorea fallax* was severely and persistently affected by the drought, dropping in *rgr* by 50% below the pre-drought value; *Parashorea malaanonan* was the only dipterocarp of the most abundant species to increase its *rgr* from the period 1986-1996 into the period 1996-2001; *Shorea johorensis*, however, showed little response (Newbery and Lingenfelder 2009). The former three species were therefore selected as the representatives of the overstorey species.

Abundant understorey species that showed strong responses to drought conditions include the ridge specialist *Cleistanthus contractus* (Euphorbiaceae; the most abundant family, 2^nd^-ranked in terms of basal area), but this species had too few trees ≥10 cm *dbh* available for sampling (these being located exclusively on the ridge of one plot). The other species that seemed to be adapted to ridges, and to be able to cope with drought-stress, is *Lophopetalum beccarianum* (Celastraceae), which was relatively unaffected during, and yet showed resilient growth after, the 1998 drought. The two most numerous species of the list of 34 were also included due to their importance in forming the understorey at Danum: *Dimorphocalyx muricatus* (Euphorbiaceae), another ridge specialist, which recovered strongly in growth, displaying a four-fold increase in *rgr* after the drought, and *Mallotus wrayi* (Euphorbiaceae), ubiquitous in both plots, which also recovered its growth (Newbery and Lingenfelder 2004, 2009). The latter three species were accordingly selected to represent the understorey species. Tag reference numbers identify the trees among the complete long-term census data. As explained later, bands on two of the 20 trees (g13, g21) were dropped.

Plot locations, species’ distributions and characteristics of the sampled trees are shown in Fig. 1, Tables 1 and 2, and Appendix 1: Table S1. Crown status was visually assessed in March 2009 and February 2010 for the variables layer, space and light each on a scale of 1 to 5. Layer was scored (in increasing order) as treelet, understorey, mid-level, top canopy and emergent tree; space (with regard to neighbours) as completely suppressed, mixed and entangled crowns, significantly competing, crowns just touching, crowns free; and light as < 30, ≥ 30 - < 50, ≥ 50 - < 70, ≥ 70 - < 90 and ≥ 90 % of full light received by crown. These scores were averaged over the two dates to give a composite variable called CanStat. Thirteen trees had unchanged CanStat values between dates, for three they decreased, and for only two they slightly increased.

**Fig. 1.**
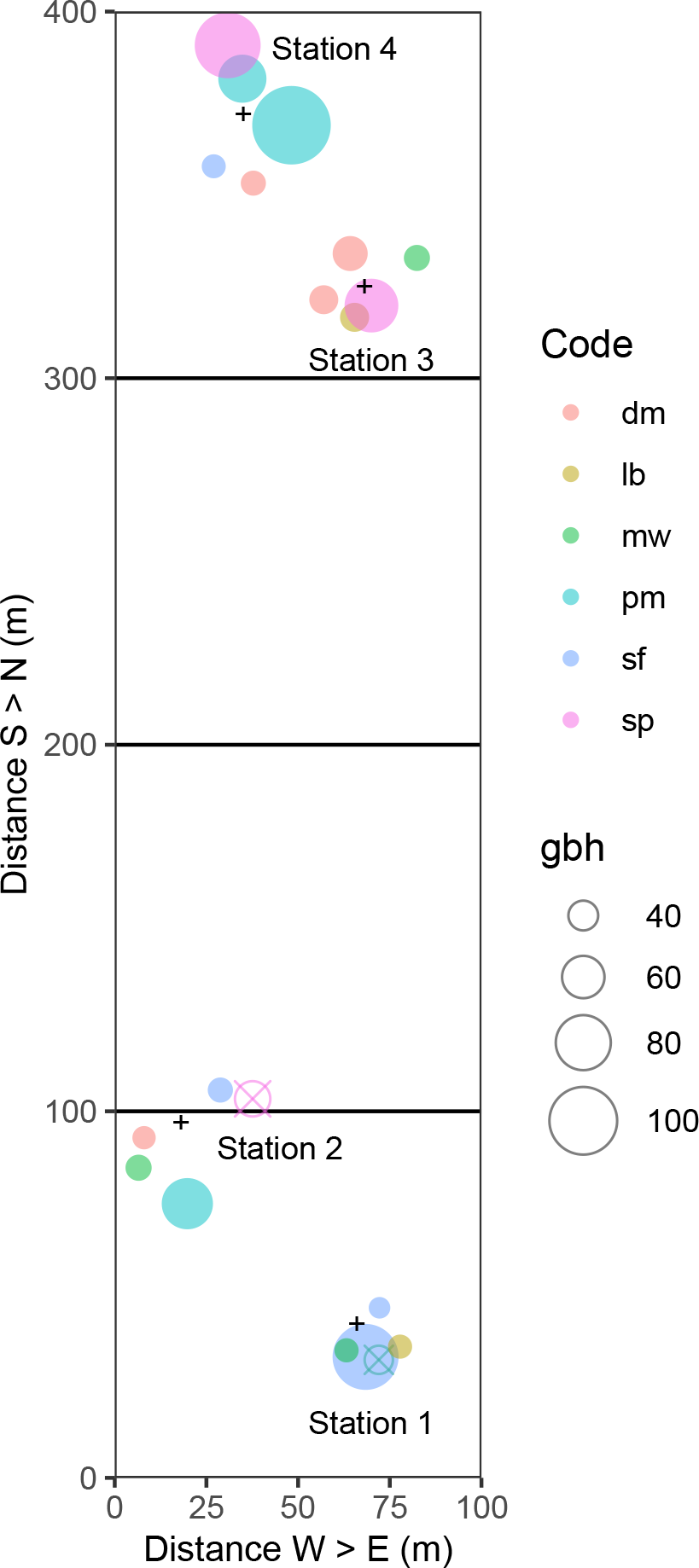
Positions of the eDDs trees in main plot 2 at Danum at the four stations. The small plus signs indicate the positions of the loggers. Closed colored symbols, scaled to tree stem girth (gbh, in cm), show the species of tree used (Table 1): codes for them are listed in Table 2. The open crossed symbols indicate the two tree (g13, g21) that were omitted from the analyses.

**Table 2.**
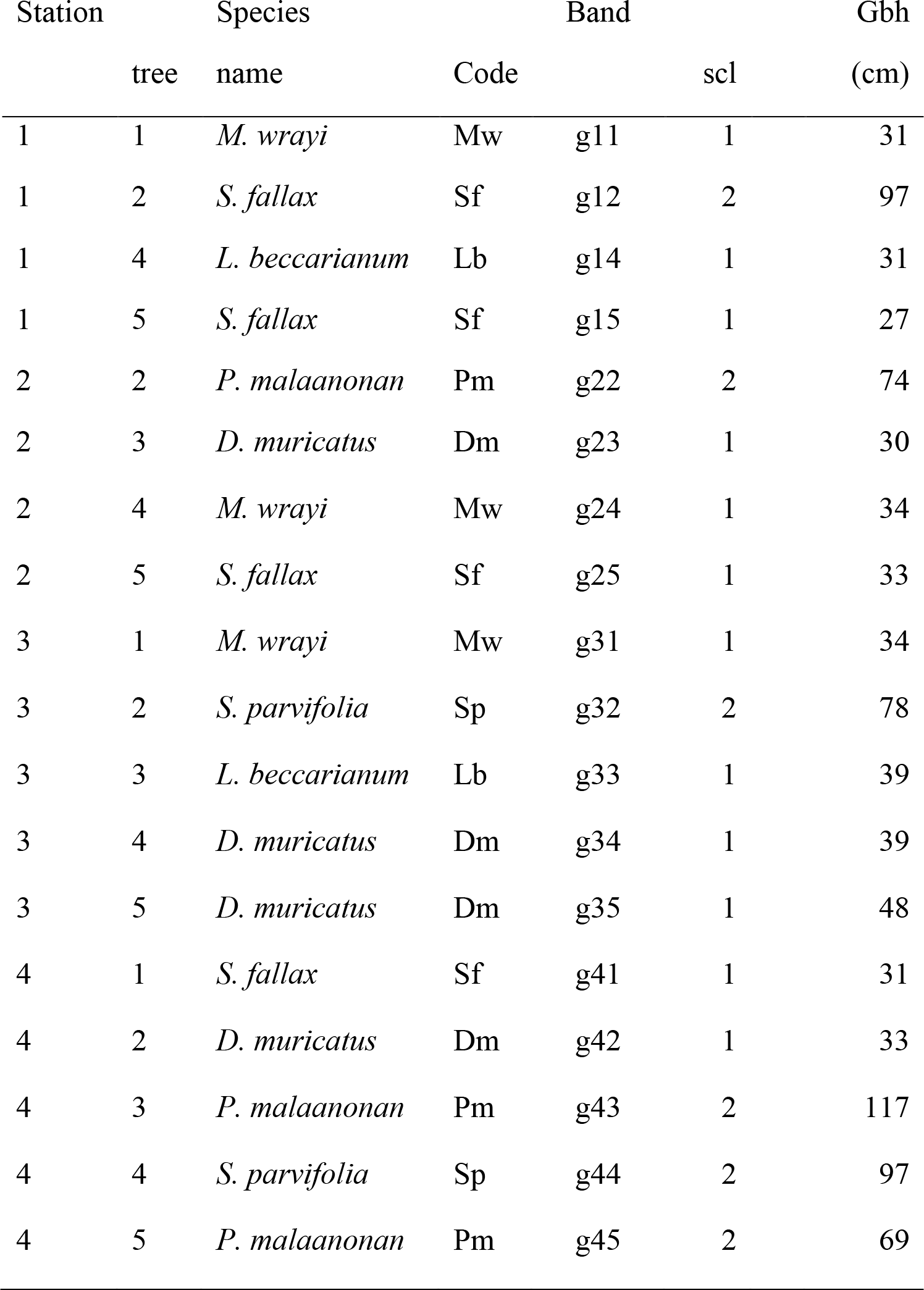
The 18 selected trees and their electronic bands (eDDS), with species names and codes, sizes (gbh, cm) at start of recording (March 2007) and corresponding size classes (scl). Two of the original 20 trees could not be used (g13 and g21: see main text). Further details and covariables are given in Appendix 1: Table S1.

### Set-up of bands

Continuous dendrometer-bands (type ‘D6’; UMS, Munich, Germany) were installed 05-08.03.2007. The device consists of an Invar-steel measuring cable with a strain-gauge sensor clip, placed around a tree and resting on a Teflon-net to minimize friction (resolution of 5 µm gbh; dependence on temperature < 4 µm/K). In plot 2, bands were fitted to 20 trees of the six species, these being selected from a larger stratified random sample of 234 candidates considered for an *in situ* parallel study with conventional mechanical dendrobands to be reported elsewhere (Table 1). Plot 2 alone was used because the required numbers of trees within reasonable cable-length distances (maximum 20 m) to loggers were more suitable than in plot 1. The trees were therefore not fully at random in the plot as they were necessarily clustered around the four loggers.

Bands were distributed across four stations (two lower slope, two ridge; Fig. 1). Each was equipped with a data-logger (CR10X; Campbell Scientific Inc., Logan, USA). On tree number 3, band g13 broke in the wet period, and there was long interval of no data whilst it was under repair; so no ‘wet’ means and limits were found. For tree number 6 (band g21) the tree died early on, and hence there were neither wet- nor dry-period data. Each station also had three to four soil equitensiometers (EQ2; Delta-T Devices Ltd, Cambridge, UK) installed, at a depth of *c*. 20 cm below ground surface, within a 10-m radius of loggers, to enable fluctuations in stem size to be related to soil water potential. Logger temperature was simultaneously recorded. The SMP correction for logger temperature, T (°C), was − (T−20)/4 in kPa, i.e. ∼ 2.5 kPa or 2%.

Measurements made by tree and soil sensors were recorded automatically at 10-min intervals, and the half-hour averages logged. Stations were downloaded every *c*. 4 wk. During the first 3 yrs, only minor difficulties were experienced (one slight disturbance by elephants, some cable damage by termites, broken fuses due to occasional lightning strikes). In 2010 problems became more frequent and serious, and accordingly there became larger gaps in the time-series. In October 2010 two stations were attacked by elephants, which resulted in severe damage to loggers and sensors so that the study had to be terminated by December 2011.

## DATA ANALYSIS

### Analysis of eDDS and SMP data

Dendroband records on trees and SMP values each spanned slightly different times for the four stations, and within stations there were differing periods of missing days. All data were first matched to a common time frame of 1614 days (48 half-hour points within each day) between 01.03.2007 and 31.07.2011 (Fig. 2). If an incidental or consecutive pair of values was missing, they were substituted by the means of values immediately before and after them. Daily means, of girth (cm) for each of the *n* = 20 trees, and of SMP (kPa) for the three or four soil sensors at each of the four stations, were found; and this allowed an identification of the complete, or nearly complete, periods of measurement. One long ‘wet’ period of 18 months (550 days), 17.03.2007 to 16.9.2008, had very few missing SMP-values (one of 15 days at station 3 and two of six and seven days at station 2), and the spectrum of SMP-values was fairly constant over time and similar across stations. SMP rarely fell below – 50 kPa and on only five days < −100 kPa. There were 3-4 short periods of about 20 days later in 2008-09 which had complete ‘wet’ records but these were amongst numerous repeatedly failing sensor runs, and showing a curious lack of a spectrum typical of the 18-month period indicated they were likely to have been unreliable throughout. However, a 3-month period, from 17.2.2010 to 16.5 2010, included three distinct yet short dry spells. These were clearly reproduced at each of the four stations, although stations 2 and 3 had a few days missing, and station 4 had a strange recording anomaly for 2 weeks.

**Fig. 2.**
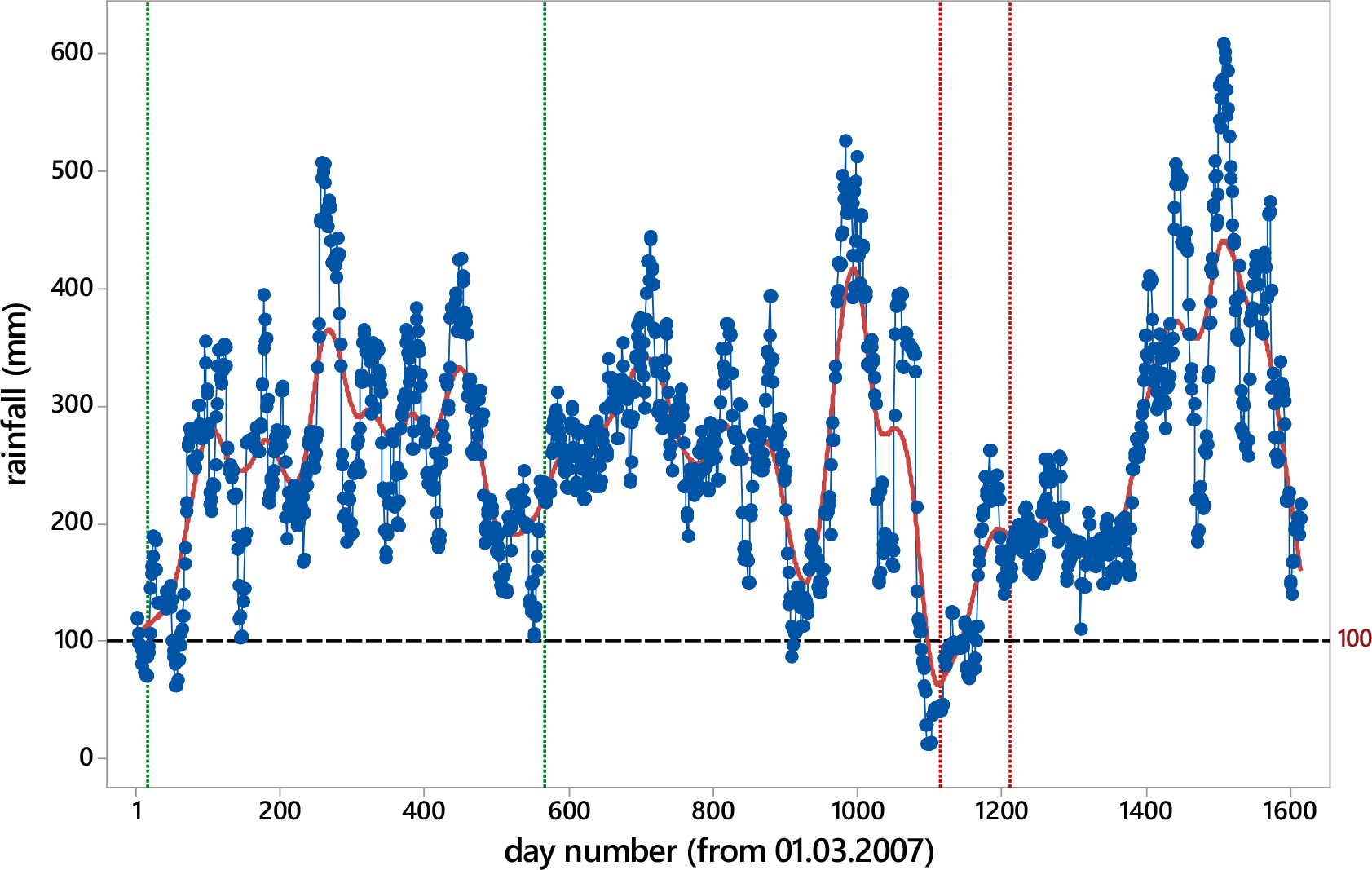
Running 30-day rainfall totals at Danum over the dendroband study period, showing the wet and dry periods used, between the green and red vertical dashed lines respectively.

The SMP values were handled in two ways to improve their robustness in defining wet and dry periods. Because of the existence of many small positive values of SMP after the conversion from logged date/time equipotentials to kPa, values were adjusted by subtracting the (upper) 0.95-quantile (per sensor) and the remaining positive values set to zero. These were called ‘offset’ SMP-values. Further, occasionally very low potentials (down to c. −2 MP), introduced by the fourth sensors at each station, appeared unrealistic. SMP-values were therefore truncated at the extremes: those positive were set to zero, and those very low values rounded up to −1000 kPa. These were called ‘strict’ SMP-values.

Stations 1 and 2 were ‘LS’ (lower slope) and stations 3 and 4 ‘R’ (ridge) in the topographic classification (Fig. 1), and averaging these pairs’ SMP-values fortunately allowed almost complete runs of data over the 3 months, and afforded a useful comparison with Gibbons and Newbery (2003). That study recorded SMP at ‘LS’ and ‘R’ locations within the same plots (slightly different sample layout) and built models to predict it from daily rainfall. SMP had then been measured with calibrated glass psychrometers. These last instruments are more reliable than the equitensiometers used in the present study, especially for lower potentials. The authors found a weighted rainfall index per day defined as w_40_ = _n=1_Σ^40^ (r_n_/n), i.e. the sum of the preceding 40 days’ rain and each weighted by its day number (e.g. *n* = 1 for the day before). This led to two equations (w_40_ in cm; SMP in MPa): one for ‘LS’, SMP = −0.0096 − 1.4070b^w40^; the other for ‘R’, SMP = −0.0525 − 2.0280b^w40^, with *b* = 0.7554. The equations were applied to the complete rainfall series at Danum, and for this study between 01.03.2007 and 30.07.2011 to give ‘predicted’ SMP at the ‘LS’ and ‘R’ stations, and thereby match the (non-missing) observed equitensiometer values.

Comparisons between the two approaches to adjusting SMP-values and their comparison with predicted values led to two clear results. (1) For ‘LS’ and ‘R’ (averaging SMP-values across the two stations each), ‘offset’ and ‘strict’ SMP-values were closely and similarly related. SMP-values were first transformed as SMPtr = √│SMP│ (e.g. −150 kPa becomes 12.25), and allowing the lower (more negative) original values (now relatively more positive transformed ones) to weight accordingly against the many low and near-zero values, the linear regressions were SMPtr_LS_ = 3.46 + 0.889 SMP_LS_ (R^2^ = 91.5%, *n* = 1350) for ‘LS’, and SMPtr_R_ = 3.27 + 0.890 SMP_R_ (R^2^ = 90.4%, *n* = 1248) for ‘R’. Substituting an ‘offset’ threshold SMP = −150 kPa in these equations gives after back-transformation ‘strict’ thresholds of −97.9 and −101.8 kPa respectively. Hence, a threshold of −150 kPa with ‘offset’ translates to −100 kPa with a ‘strict’ setting. (2) Regressing ‘offset’ and ‘strict’ SMP-values against w_40_-predicted SMP (days where the latter was ≤ − 50 kPa for ‘L’, ≤ − 100 kPa for ‘R’; *n* = 89 and 94 respectively) showed significant, though not particularly strong, quadratic relationships (R^2^-values of 57.3 to 67.0%). ‘Offset’ SMP-values gave much closer to w_40_-predicted values than ‘strict’ ones at ‘LS’ (the latter underestimating by c. 20%), whilst both ‘offset’ and ‘strict’ (more so) underestimated w_40_-predicted values at ‘R’, by c. 30 and 45% respectively. It seems then that SMP-values on ridges were likely, in reality, to have been lower (taking the psychrometer values too as being the more reliable) than those recorded by the equitensiometers.

Dry spells were therefore defined as runs of (> 3 days) with ‘LS’-and ‘R’-averaged ‘offset’ SMP-values ≤ − 150 kPa, or equivalently as ‘strict’ values ≤ − 100 kPa (Appendix 1: Fig. S1). The three ‘dry’ spells were thus: 18.2.-7.3.2010 (18 days), 12.3 – 28.3.2010 (17), and 11.4 – 28.4.2010 (18). If ‘R’ values had been corrected according to the w_40_-predicted regression equations, this threshold would have been lower still after averaging stations; it would have led to slightly different periods being defined as ‘dry’ on ‘R’ compared with ‘LS’.

But differences in SMP between ‘dry’ and ‘wet’ are so large, and events developed so rapidly down to values of − 250 kPa and lower, that this was considered unnecessary.

Daily temperature (TEMP) was taken as the mean of logger internal readings at 14:00 h, across the four stations, over the period 6.03.2007 to 4.12.2010. Mean TEMP (with SE) was 26.91 ± 0.04 _o_C (*n* = 1370). Over the study period, recordings at the nearby DVFC climate station were incomplete for many variables other than rainfall, but those days with matching values showed dry-bulb temperature at 14:00 h (Tdry14), maximum daily temperature (Tmax), and number of sunshine hours all to be significantly positively (*r* = 0.734, 0.804 and 0.652; *n*’ = 991, 1333 and 786; respectively), and relative humidity at 14:00 h (RH14) significantly negatively (*r* = −0.658, *ń* = 983), correlated with TEMP. Correlations using only stations 2-4 were very similar. Plotting graphs showed the relationships all to be linear. TEMP acts as a surrogate for evapotranspiration pull by the trees: it is used as one of the two main predictors of *gthi*, along with SMP.

SMP correlated fairly strongly and positively with 10-, 20-, 30-, 40- and 50-day running rainfall totals (both variables sqrt-transformed to achieve linearity), in both wet and dry periods, most strongly with the 10-d value in the wet, and the 20-d one in the dry, period (Table 3). The relationships between SMP and 20-day rainfall totals were adequately described by linear-exponential curves (Appendix 1: Fig. S2). Fits were statistically similar in terms of *F*-ratios for both periods, but the wet with ∼ 5.5 times the number of observations of the dry, showed considerably more scatter in SMP (∼ 0 to – 120 kPa over the range 100 - 200 mm rainfall totals) than the dry and correspondingly half the *R^2^*-value. SMP was significantly negatively correlated (*P* ≤ 0.001) with temperature (TEMP; Tdry14 and Tmax at DVFC), particularly with the first (Table 3). Temperature variables were also strongly positively correlated among one another in both periods (*P* ≤ 0.001).

**Table 3.** Correlations (Pearson’s *r*) between mean soil water potential, SMP [sqrt-transformed] and logger temperature at 14:00 h, TEMP recorded at the stations), for the wet and dry periods, and climatic variables recorded at the Danum Valley Field Centre – or derived from them (10-, 20-, 30- and 40-d running rainfall totals, rain10rft to rain50rft, from daily rainfall [sqrt-transformed]: dry-bulb temperature at 14:00, Tdry14, maximum daily temperature, Tmax. All correlations are significant at *P* ≤ 0.001. SMP and TEMP are not lagged.

| variables: |  | wet |  |  | dry |  |  |
| --- | --- | --- | --- | --- | --- | --- | --- |
| | | $r$ | $t$ | $n$ | $r$ | $t$ | $n$ |
| SMP | rain10rft | 0.681 | 21.74 | 550 | 0.776 | 12.05 | 98 |
|  | rain20rft | 0.553 | 15.53 |  | 0.782 | 12.30 |  |
|  | rain30rft | 0.501 | 13.56 |  | 0.761 | 11.49 |  |
|  | rain40rft | 0.485 | 12.97 |  | 0.756 | 11.33 |  |
|  | rain50rft | 0.557 | 15.70 |  | 0.741 | 10.81 |  |
| SMP | TEMP | -0.389 | -9.89 | 550 | -0.449 | -4.92 | 98 |
|  | Tdry14 | -0.225 | -4.36 | 357 | -0.311 | -2.64 | 67 |
|  | Tmax | -0.254 | -5.97 | 520 | -0.361 | -3.77 | 97 |
| TEMP | Tdry14 | 0.739 | 20.64 | 357 | 0.713 | 8.21 | 67 |
|  | Tmax | 0.747 | 25.58 | 520 | 0.818 | 13.85 | 97 |
| Tdry14 | Tmax | 0.664 | 16.30 | 340 | 0.716 | 8.200 | 66 |

Diurnal tree girth change was calculated as *per mille* mm with reference to girth at midnight (t = 1), i.e. at 00:00 of the new day, using the formula: gthch_t_ = ((girth_t_ – girth_1_)/girth_1_) ·10^4^, where t = 1, 2, … 48 for the half-hour times through the day. Whenever a diurnal time series was incomplete, the series of changes was set to ‘missing’ and that day did not count in the finding of means per time of day in a period. The multiplying constant brings the values of relative change into a range of 0 to c. 20. In consideration that showers on certain days in the dry period may have affected band readings due to the rewetting of the generally relatively dry bark, data for four days within the three spells with ≥ 10 mm per day (viz. days with 19.2, 25.5, 39.3 and 43.7 mm) were removed from the calculations of *gthch*. The maxima of rainfall remaining in the three spells were 5.7, 6.5 and 7.0 mm. Most other days had zero rainfall. In the following ‘*Time-series analyses*’ section, regressions of girth increment per day on rainfall suggested that showers of < 10 mm would very likely not have affected band readings to any appreciable extent.

To test whether daily patterns in girth changes differed significantly in any of the three (17-)18-day dry periods from the longer wet one, the latter was divided into 30 consecutive and similarly long 18-day periods (ignoring the last 10 of the 550 days, and assuming periods to be independent). Mean changes were found per 18-day wet or dry period, and for the wet one the mean of those 30 means and their 99% confidence limits were computed. Should one or another of the dry-period individual means lie outside these limits, the null hypothesis — that changes in wet and dry are the same— can be rejected at *P* < 0.01. This assumes that no other factors confounded comparisons over time. Variation in mean diurnal SMP for the 30 wet sub-periods and three dry ones (together) was extremely small, only dry-3 showing some downward trend after 10:00 h (Appendix 1: Fig. S3a). By contrast, TEMP fluctuated in a clear manner over the day, peaking at *c*. 15:00 h, higher in dry than wet period (Appendix 1: Fig. S3b).

Means of SMP in the three dry sub-periods were −260, −411 and −290 (together −320) kPa, compared with −36.3 (± 3.7) kPa for the 30 wet sub-periods. These dry sub-periods did not differ significantly in SMP from one another (repeated measures analysis of variance, *F* = 8.24, df = 2, 4; *P* = 0.097), nor in topographic location with the plots (lower slope, −354; ridge −287, kPa, P ∼ 0.4). The nine-fold difference in SMP between wet and dry periods underwrote the comparison of following tree girth change analyses.

In the following analysis, SMP_0_, SMP_–1_, and SMP_–2_, and TEMP_0_, TEMP_–1_, and TEMP_–2_, are respectively SMP, and TEMP, values at no, one- and two-day lags. Calculations were carried out using Fortran f77 code (FB 6.0, Numerical Algorithms Group, Oxford) to program basic data handling, followed by exploratory analysis and statistical summaries in Minitab (vers. 18, Minitab LLC, Pennsylvania), non-linear regression models in GenStat version 18 (Payne et al. 2011), the time series regressions in R (R_Core_Team 2017), and most of the graphics in `ggplot2` in R (Whickam 2016). Two valuable guiding texts were Venables and Ripley (2002) and Maindonald and Braun (2010).

### Time-series analyses

All trees, minus g13 and g21 were taken with dates (*n* = 550 days) and complete SMP (of four stations) data for the wet period. Numbers of values to be imputed: g14 (12 d), g25 (1 d), g23 (6 d), g22-25 (8 d, 9 d), g32 (3 d), g31-35 (17 d), g41-45 (1 d). These imputed values were set equal to the average of the last and next values. Usually they differed from neither, or from just one by 0.01 cm. In this way a complete set of 550 dates x 18 trees (g11 to g45, minus g13 and g21) was available for autocorrelation analysis. The finding of missing values and their imputation was achieved in R with the package ‘zoo’ version 1.8-6 (Zeileis and Grothendieck 2005). Simple wet- and dry-period time series for each girth band are given in data archive, together with their SMP and TEMP series.

For the dry period to be continuously complete as possible several aspects had to be considered. The 6-mo span 01.01 to 30.06.2010 would have been ideal; here station 1 would have been be possible, stations 2 and 3 for 24.02-23.06 only though, and station 4 not at all (because of too many large gaps). By excluding station 4, it was feasible to have 24.02-23.06 as a set ‘A’, and 18.03-23.06 as a set ‘B’. Imputation was as follows: g22-24 (5 d), g31-35 (12 + 2 d), g23 (three scattered pairs of days), so that for set ‘A’ this meant for stations 2 and 3, 8 and 14 d imputed respectively, and for set ‘B’ 8 and 2 d. This gave finally a max set ‘A’ set of 120 dates x 13 trees, and or for set ‘B’ 103 dates x 13 trees (i.e. as for wet, minus the station 4 trees. To allow sets A and B to be as closely matching as possible, yet make maximal use of the information available, set A was taken from 18.03.2010 to 23.06.2010 (98 d), so-called set A(B) in the archive, and set B 18.03.2010 to 18.06.2010 (93 d).

To remove the effects of showers occurring on some of the days in the dry period, ones which might have caused temporary bark swelling so that changes in *gthi* were not accurately reflecting changes in *gth* related to tree water status per se, quadratic regressions were fitted to the relationship between *gthi* and daily rainfall (ln-transformed) for each band, for those days with ≥ 1 mm rain. For days with 0 mm rain, mean *gthi* was found. From the fitted regression curve, and the mean-at-0 mm, *gthi* residuals were then used in place of original *gthi.* This catered for band expansion and contraction related to immediate wetting in functional manner, that is any change in this respect would be expected to be a physical non-linear function applying in common to each day of an individual tree’s recording. Using all days of the dry period (i.e. those with rain ≥ 0 mm) in one regression model would have violated the assumption of an even approximately homoscedastic distribution of values along the independent variable’s axis; because many days had no rainfall at all. Quadratic regressions were first run using rainfall on the current day, and then rainfall on the previous day.

### Arima and GLS regression

Mean girth, *gthm* (cm), was found for each day that had complete recordings for the forty-eight 30-min intervals, for each tree in the dry and wet time series. For each day *i*, increments were then found for an interval of 1 day ahead, i.e. *gthi* = (*gthm_i+1_* − *gthm_i_*) · 10^3^, i.e. finding the mean *gth* per complete day first, then the increment across days and not the mean of *gthi*-values per day. Corresponding mean SMP-values were transformed as – 1· (|SMP|)^1/2^, to reduce the strong left skew in the data, and matched to each day’s increment.

The dependence of girth increment per tree on SMP in each time series (presented in the data archive associated with this paper) was modelled in two ways: (1) by ARIMA, with SMP as an independent variable, and the parameters p and q fitted either automatically, each allowed to vary up to values of 2, or fixed at either 1 or 2 (Brockwell and Davis 2002, Chatfield 2004), with analysis performed in R using the package ‘forecast’ (function *auto.arima | Xreg*; (Hyndman and Khandakar 2008); and (2) GLS, generalized least squares, regression, where temporal autocorrelation was accounted for with p and q set at either 1 or 2 each (Pinheiro and Bates 2000, Fox 2008), performed with the R package ‘nlme’ (function *gls | corAMRA*; (Fox and Weisberg 2011, Pinheiro et al. 2013). ARIMA and GLS analyses were repeated with SMP lagged by 1 and 2 days.

Parameter p is the degree of lag in the dependent variable included in the autoregressive model, and parameter q is the number of current plus proceeding error terms that are also included (as moving averages) (Brockwell and Davis 2002). The values of p and q used in the final GLS models were thus guided by the best fitting ones from ARIMA. Decomposition of the time-series was not possible. Besides there being no clear seasons at Danum, the series were shorter than 1 year (550 and 98 days respectively for the wet and dry periods). Determination of p and q for the dry period used the original *gthi*-values.

The first ‘automatic’ ARIMA with the three SMP lags selected p = 1 or 2 (occasionally 3), q = 1 or 2, and d = 1, most often for the best fitting models. By placing restrictions of p ≤ 2 and q ≤ 2, very similar results were obtained, and leading to generally lower AIC-values. Differencing was mostly not required (6 or 7 of 18 wet, and 0 or 1, rarely 2, dry bands, had d = 1 selected). Plots of the partial autocorrelation function (ACF) showed that for p ≤ 2 the functions were often significant (*P* < 0.05), although its ± value declined relatively slowly over higher p-values, indicative of some heteroscedasticity. Differencing in the form of *gthi*, prior to time-series analysis had the added benefit of making the series more stationary; it also removed net accumulated growth (*gbh* increase) over the time span.

Evaluation of p and q for the dry period used the original *gthi*-values.

Since the aim of ARIMA here was to find an omnibus for removing autocorrelated variation in the regressions, and not for forecasting, and to a large degree differencing had *de facto* been achieved in the calculation of growth increment, tree-specific settings of d = 1 were not used. Such ‘differences in differences (i.e. increments)’ would also make physiological interpretations obscure. Furthermore, when d = 0 a constant usually (as in R) is estimated by ARIMA. On fitting SMP as the independent variable, this time-series constant would be integrated within the regression intercept term. The value of that final constant is not of prime interest, the focus being response in incremental growth to change in SMP. The p = 2 and q = 2 parameter settings were therefore taken for the *corARMA* function in GLS. Thus with p = 2, the current, day-before and two-days-before growth increments are terms in the model, and with q = 2, so also are the current and the day-before errors as terms. It led to just four fitting failures, these due to the coefficients matrix being non-invertibile (Brockwell and Davis 2002): they were remedied by setting p = 3 or q = 1. Overall, using the same parameterization meant a common model for all trees, across both wet and dry time-series.

For one tree (g22) a sharp hiatus in recordings (likely caused by falling branch damage to the dendroband) on 27-28.05.2007 meant that only the first part of the wet series could be used. SMP coefficients from GLS with q < 2 were highly correlated with those with q = 2 across species within series indicating strong robustness in the estimations. (For q = 1, wet: 0.993, 0.973 and 0.991; dry: 0.985, 0.971 and 0.983. For q = 0, wet: 0.993, 0.971 and 0.951; dry: 0.987, 0.970 and 0.993, lags 0, 1 and 2 respectively; *n*_wet_ = 18, *n*_dry_ = 13, *P* for all << 0.001). The final GLS regressions with p = 2 and q = 2 were therefore taken as the most robust and realistic set of tests of the influence of SMP on *gthi* adjusted for temporal autocorrelation.

In preliminary runs of time series analysis (not reported here) using growth increments over 2- to 5-day intervals, interpretation was evidently compromised and made very difficult when q (the moving average term) was ≥ 1. For example, a 3-day increment using q = 0 is a sum, whose average daily equivalent equals the 1-day increment value for the middle day of the three when setting q = 2; this middle day being 1 day advanced with respect to the SMP value (on the first day). Working with growth increments of ≥ 2 days in ARIMA and GLS would necessitate finding mean SMP per increment, which would then lose the diurnal resolution. On these grounds analysis was limited to 1-day growth increment series. In addition, using longer increments led to increasingly higher proportions of model fits giving warnings of statistical conditions being not met in ARIMA, or failing.

GLS regressions were also fitted for *gthi* versus TEMP in the same way as for SMP, and then for both terms SMP*TEMP to assess the significance of the interaction. In the latter case only lags of 0 and 1 days were tested. Since SMP_−1_ was correlated with rainfall totals, *rft* (Table 3), to judge how far the *gthi* regressions on SMP_−1_ might be accounted for by these totals, GLS regressions (p = 2 and q = 2) were run with 10-d-*rft* in the wet period and 20-d-*rft* in the dry (given their improved correlations over 30-d-*rft*). Regressions fitting SMP_−1_ to *gthi* were then repeated with corresponding *rft* terms as the second predictor.

For comparisons across trees and species, and in relation to other variables, SMP_−1_ and TEMP_0_ coefficients from the time-series GLS fitting were standardized to 50 cm *gbh* by multiplying by 50.0/*gbh* (in cm).

### Non-stationarity

After fitting auto-arima models (Xreg = SMP) with lags in SMP of 0, 1 or 2 days, the ACFs of the squared residuals were examined for evidence of non-stationarity of variances, and any departures fitted using GARCH models (Enders 2015). These calculations were made with R package ‘tseries’ (function *garch*). In testing at *P* < 0.05, emphasis was placed on ACF for p = 1 to 3, and whether ACFs (ε^2^) were *P* << 0.05 significant. Occasional small divergences at p = 1 to 3 were counted as weakly non-significant. For SMP lag = 0 (and 1 and 2), three of the 18 ‘wet’ bands (viz. g14, g23 and g41) showed obvious and strong heteroscedasticity, removable with GARCH model fits (p = 1, q = 1) for bands g14 and g41 but not band g23 (with any p = 1 to 3). Band g22 had an isolated very high variance episode early in the time-series and this was clearly highlighted by the ACFs (ε^2^) and GARCH.

Using SMP lags 0, 1 and 2, for a second test of non-stationarity, the wet series of 18 bands were each divided into three parts. For the four bands where ACFs (ε^2^) indicated non-stationarity (heteroscedacity), were divided unequally so as to have a ‘normal’ subseries, then a volatile part, and finally a return to ‘normality’. For the other 14 bands the series was divided into equal parts. `Auto.arima` (max.p = 2, max.q = 2) and `gls` (p = 2, q = 2) were run for each band, and estimated SMP coefficients for subseries plotted against one another. (In three of 54 cases it was necessary to reset q to 1 to obtain a solution.) The coefficients from arima and GLS generally matched very well for each subseries, but not always well *between* them (within series). Of the four ‘suspected’ series, g14 and g22 certainly behaved differently in the middle periods (i.e. not only was variance higher but slope different), whilst for g23 and g41 there was scarcely a difference (i.e. despite the raised variance, slopes remained similar). For g14 and g22, coefficients were respectively much more positive and negative than the low variance subseries before and after.

To summarize, the tree/species *gth* vs SMP relationships with minimum effect of volatility, GLS coefficients for g14, g23 and g41 bands were averaged for their first and third subseries, and combined SEs found as the root mean squares (geom. means) of the two subseries’ values. Likewise, the two *t*-values were averaged and a new *P*(*t*) found (with (550/3) – 2 = 181.3 df). For band g22, the first 100 days of the wet series were omitted because, beside the extreme volatility early on, there was the above-mentioned unusual down- shift in *gth* in May 2007; and thus its GLS was based on a third subseries of 450 days. The four bands’ new estimates replaced the full series ones, giving more robust and conservative time-series regression model fits for them.

### Assessing effects of relative humidity on dendroband measurements

To test whether a fall in RH in the understorey on dry days (especially in the mid- afternoons) could have affected dendroband readings because of possible bark shrinkage, external climate data from the DVFC station were used. Although RH was not recorded in the forest understorey, it has to be assumed that a drop in RH outside in a clearing would be paralleled by a drop inside the forest, even if the latter was less steep and the absolute change was not as great. Dry-bulb temperature and RH at 14:00 h (Tdry14, RH14) were significantly inversely correlated (*r* = –0.896, df = 352, *P* < 0.001 in the wet, and *r* = –0.932, df = 65, *P* < 0.001 in the dry, periods). These temperatures are likely to be very similar to those above the forest canopy. Temperature, together with rainfall, are the principal causes of RH. The changes in *gthi* resulting from water flux are being driven by radiation and thus temperature.

In the wet period the SMP is continuously high, therefore any effects of RH on bark properties and hence *gthi* readings, might be shown once the influence of temperature on RH has been removed (see Warner 2013 for a discussion of this approach). Accordingly, RH14 was regressed on Tdry14, and the residuals then used as a new independent variable to modeling *gthi* with GLS-arima regression. This will not use absolute RH but the deviation at a given temperature. Alternatively, fitting *gthi* to Tdry14 + Tdry14·RH14, RH14 is nested within Tdry14 ‘levels’ so that the coefficient of the interaction term expresses the rate of change of the dependence on RH14 as Tdry14 increases, i.e. RH14 is conditional on Tdry14 (e.g. Aiken and West 1991). The main effect of Tdry14 is partialed out, the collinearity between Tdry14 and RH14 is avoided, and RH remains on its original scale.

A more direct, but one involving fewer data, way to test for effects of RH aside from temperature was to select very narrow slices in temperature and then relate *gthi* to RH within each of them, assuming that the differential effect of temperature was then minimal. The effect of temperature on *gthi* could be minimalized by selecting days on which the soils were fully wet and therefore presumably creating minimal resistance to tree water flux. Of the 550 days of the wet period, 153 had current- and previous-day soil moisture potentials (SMP_0_, SMP_1_) both > –20 MPa. Of these last, however, only 90 days had records of Tdry14 and RH14 from the DVFC station. The largest amount of vertical spread in RH-values was between 29 and 31°C. Points were divided into three 0.7-°C-wide slices: (1) 29.3 – 29.9, (2) 30.0 – 30.6, and (3) 30.7 – 31.3 °C (*n* = 16, 22 and 14 respectively). On these days, within each of the temperature slices, ET was expected to have been very similar: what remained to affect *gthi* via any putative bark changes would have been principally RH.

### Granger causality

Granger causality testing provides an omnibus method for detecting whether for two time-series variables, Y_t_ depends on X_t_ when both are included in an auto-regressive model, each with increasing lags in *t* up to *p* (Granger 1969). A significant *F*-ratio indicates whether at least one X_t_ term is determining at least one Y_t_ term. The test was implemented with `grangertest` in R package `lmtest,` for *gthi* as Y and either SMP_0_ or TEMP_0_ as X, for the wet and dry periods separately. By setting *p* = 2, SMP_−1_, SMP_−2,_ TEMP_−1_ and TEMP_−2_ were included in the model.

## RESULTS

### Relative humidity and potential effects on dendroband recordings

Dry-bulb temperature and relative humidity (Tdry14, RH14) were very similarly — and close to Gaussian — distributed in the wet and dry periods (Appendix 2: Table S1). Mean Tdry14 was just 1.4 °C higher in dry than wet, and RH14 3.5% lower. Ranges in these variables were also very similar, with RH14 minima achieving 60% in both periods, and maxima 98-100%. Thus the potential for low RH affecting bark properties and thus biasing dendroband readings was similar in both periods, despite the dry one being ‘drier’ than the wet one in terms of 30-day rainfall totals and SMP.

Evidence for relative humidity affecting *gthi*, either by applying the residuals of fits of RH14 to Tdry14, or using the nested interaction term, was found for just a single band (g14), the same one in both periods, and then with only marginal statistical significance (Appendix 2: Table S2). The approach which analyzed the effects of RH14 within very narrow ‘controlled’ slices of Tdry14 provided even less evidence. In none of the 54 cases (three temperature slices x 18 dendrobands) was significance at *P* < 0.05 attained. In slice Tcl2, there was a marginal correlation of *gthi* and RH14 for g12 (*r* = –0.431, df = 20, *P* = 0.051) and, in that same class, a weaker correlation for g42 (*r* = 0.404, df = 20, *P* = 0.069) of opposite sign. All other cases were with *P* > 0.10, the majority with *P* = 0.3 to 0.9). Appendix 2: Fig. S1 reveals no clear patterns with sometimes weak positive, sometime weak negative, trends; and no patterns even across temperature slices for the same tree band. Together these tests indicate that the effect of mid-afternoon drops in RH on some days had scarcely any consistent influence on *gthi*, and can be dismissed as a potentially biasing factor. Important also to note in that any small drop in RH in part of the day was happening in both wet and dry periods with a similar frequency.

### Temperature and band mechanism

Any errors introduced by expansion or contraction of the bands due to day-to-day temperature changes (Pesonen et al. 2004, Wang and Sammis 2008) on temperature-dependent accuracy of band dendrometers were found also to be very small under the Danum forest conditions and could therefore be neglected (see Appendix 3 for an analysis). They were certainly very minor compared with diurnal ranges and day-to-day differences.

Accordingly, the analyses presented for SMP and temperature effects on *gthi* are presented without adjustment. No correction or allowance was made for possible thermal expansion of stem wood though. This is likely to have been of the same order of magnitude as the bands (Sevanto et al. 2005) but estimation of the necessary parameters for the six Danum tree species was not feasible during the present study. Presumably if temperature increase were to lower RH and shrink the outer bark, it would simultaneously expand the inner bark and sapwood, creating a tension. The thin bark, especially on understorey species, would have needed to evolve an ability to stretch in order to avoid splitting.

### Effect of rainfall in the dry period on dendroband recordings

In the dry period 38 of 98 days had no rainfall, most of the remainder received very light rainfall, and a few had heavier showers (Appendix 4: Fig. S1). For nine of the 13 bands the quadratic relationship of *gthi* on rainfall (ln-transformed) was significant at *P* < 0.01 (Appendix 4: Table S1). However, of the nine it was not always positive: for g15 and g35 there was significant decline at *P* < 0.01, and likewise for g32 at *P* < 0.05. The relationship of *gthi* to rainfall on the previous day was largely non-significant across the bands, but there were two exceptions: for g12 and g33, both with positive slopes (Appendix 4: Fig. S2).

Interestingly, for those two bands the relationship was insignificant with current day rainfall. Mean of *gthi* on the days with zero rainfall matched up well with the line fitted across 1-3 mm rainfall (Appendix 4: Figs S1 and S2), justifying, in a non-formal way, the combining of residuals about the 0-mean with those from the regression curve. Using the *gthi* residuals of the fitted rain function satisfactorily catered for parts of *gthi* that putatively could have resulted from bark swelling, even when the effect was weak or opposite to what would have been expected. The important adjustments were for those few days with > 15 mm, although for not all of them because the scatter in *gthi* on those wetter days was appreciable (Appendix 4: Figs S1 and S2). Across all rainfall values the variation in *gthi* was indeed wide. Short rains would probably not have penetrated far into the top soil layer, evidenced by the remarkable constancy of the SMP readings within the dry period.

Comparing estimates of the SMP and TEMP coefficients (at three lags) from the single-term GLS-arima model fits, using the *gthi* rain-residuals and those using the original *gthi*-values showed rather little difference overall to the general pattern of effects though (Appendix 4: Fig. S3). The 39 (13 x 3) pairs of coefficients – not fully independent because of correlations between lags – were linearly related (SMP, *r* = 0.864, TEMP, *r* = 0.914; df = 37, *P* < 0.001 both). Corresponding reduced major axis regressions estimated slopes of 1.264 and 1.019 for the SMP and TEMP coefficients respectively. In both cases, lines passed close to the origins, which means for SMP in particular that by using rain residuals, both positive and negative coefficients *increased* in size by ∼25%. For TEMP there was, on average, no change but individual coefficients, particularly the positive ones, altered plus or minus more than the SMP ones. The final GLS-arima results for the dry period are nevertheless those which now used *gthi* rain residuals in place of *gthi* per se.

### Diurnal variation

All 18 trees allowed a comparison in daily *gthch* between wet and dry periods, expressed on a per mille (°/oo) basis and referenced to zero-midnight. The 30 sub-periods showed a repeatable and consistent pattern, an increase (swelling) in *gthch* in the mid-morning and a decrease (shrinking) in mid-afternoon (Fig. 3). The diurnal variation was barely detectable for large trees over 60 cm *gbh* in this wet period, particularly the dipterocarps; it was more pronounced for the understorey species *D. muricatus* (`Dm`), *L. beccarianum* (`Lb)` and *M. wrayi* (`Mw`), and for the smaller trees of the dipterocarps (Fig. 3). In the dry period compared with the wet one, again the large dipterocarps showed only small increases (outside of the wet period confidence limits), suggesting that they were little affected by the drier conditions, but the small dipterocarps were much more affected especially in the mid-afternoon with strong decreases. The understorey species showed an interesting mix of response curves with generally an increase in *gthch* in the late morning, and even into the afternoon (especially for *M. wrayi*), and then a decrease in mid- to late afternoon.

**Fig. 3.**
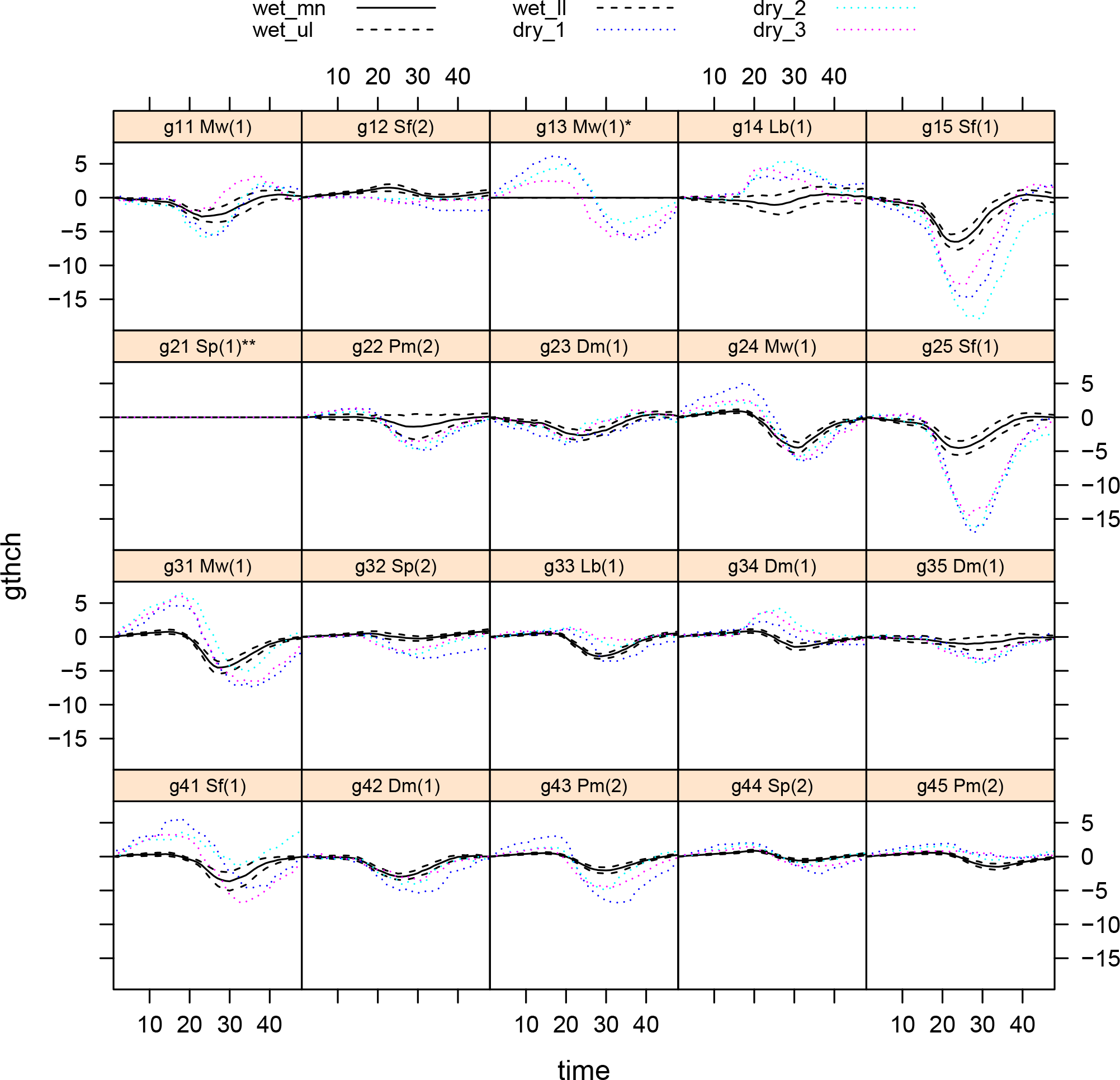
Mean diurnal change in tree girth (*gthch*), integrated for 48 half-hour time intervals referenced daily to 00:00, expressed per mille, for each of 20 originally selected trees. Two trees’ series (see Fig. 1) were insufficient and were later omitted. Panel headers: following the band codes, are species codes and, in parenthesis, tree size classes (*scl*), as listed in Table 2. The solid black curves are the means of the 30 18-day sub-periods of the wet period and the black dashed lines their 95% confidence limits. The dotted colored curves are the three sub- periods of the dry period. Positive values of *gthch* indicate stem swelling, and negative ones, stem shrinkage. Each successive row of panels is for stations 1 to 4, each row with its four- five trees (see Table 2).

By contrast to *M. wrayi* and *L. beccarianum, D. muricatus* showed little response (Fig. 3). Interesting also is that among the large-treed species, *P. malaanonan* (`Pm)` tended in its behaviour towards the understorey ones. Largest responses diurnally were again for the small trees of the overstorey dipterocarps, very clearly for two of these three *S. fallax* (`Sf)` trees, one with a late-morning increase like *M. wrayi*. The two trees of *S. parvifolia* (`Sp)` behaved quite differently in the dry period but not markedly so. Small dipterocarps appeared therefore to be strongly affected by the dry conditions, the understorey species little in comparison.

Among all 18 trees (*scl* = 1 and 2, Table 2), the six species did not differ significantly from one another in their mean minimum or maximum diurnal *gthch*-values in either period (*P* = 0.20 – 0.80; unbalanced analysis of variance, with stations as randomized ‘block’ term). For the 12 small trees (*scl* = 1), the four species showed barely more significance between them across these two *gthch* variables and periods (*P* = 0.059 – 0.65): there was some indication of `Sf` being marginally significantly lower than `Mw` in *gthch_min_* in the dry period (see Fig. 3).

### Changes over longer time in relation to soil moisture and temperature

The growth increment, *gthi*, i.e. between the current day (*gthm*_i_) and the next (*gthm*_i+1_) was regressed against the current day’s mean soil moisture potential (SMP_0_), that of the day before (SMP_−1_) and of two days before (SMP_−2_). To recall, four bands’ series in the wet period were adjusted for non-stationarity. Bonferroni correction to the *P*-values for the regression *t*-values was applied by dividing the critical α by 18 and 13 for the wet and dry periods’ samples of trees respectively, quite conservatively guarding against type I errors due to multiple testing. In the results reported in the next three paragraphs ‘significance’ in this context refers to the adjusted *P*-values.

Of the 18 wet period trees, *gthi* was significantly related to SMP_0_ for eight of them (six negative, two positive), but with many more for SMP_−1_, with 14 trees (eight negative, six positive), and then with 11 trees for SMP_−2_ (six negative, five positive) (Fig. 4; Appendix 5: Table S1 and its footnote). For most trees the highest significances were attached to the slopes was for SMP_−1_, though sometimes they were similarly strong for SMP_0_. In all but two cases the direction was the same across SMP_0_ to SMP_−2_, and in those opposite cases the change was with insignificant slopes for SMP_0_. Of the 13 trees in the dry period, five were significant (two negative, three positive) for SMP_0_, eight for SMP_−1_ (three negative, five positive), and then five again for SMP_−2_ (two negative, three positive). Two conclusions follow. The dependence of *gthi* on SMP was strongest (i.e. most significant) largely with the soil potential of the previous day (SMP_−1_), in both wet and dry periods. Significance will depend in part on sample sizes: the wet had more time points than the dry period (Appendix 1: Table S3).

**Fig. 4.**
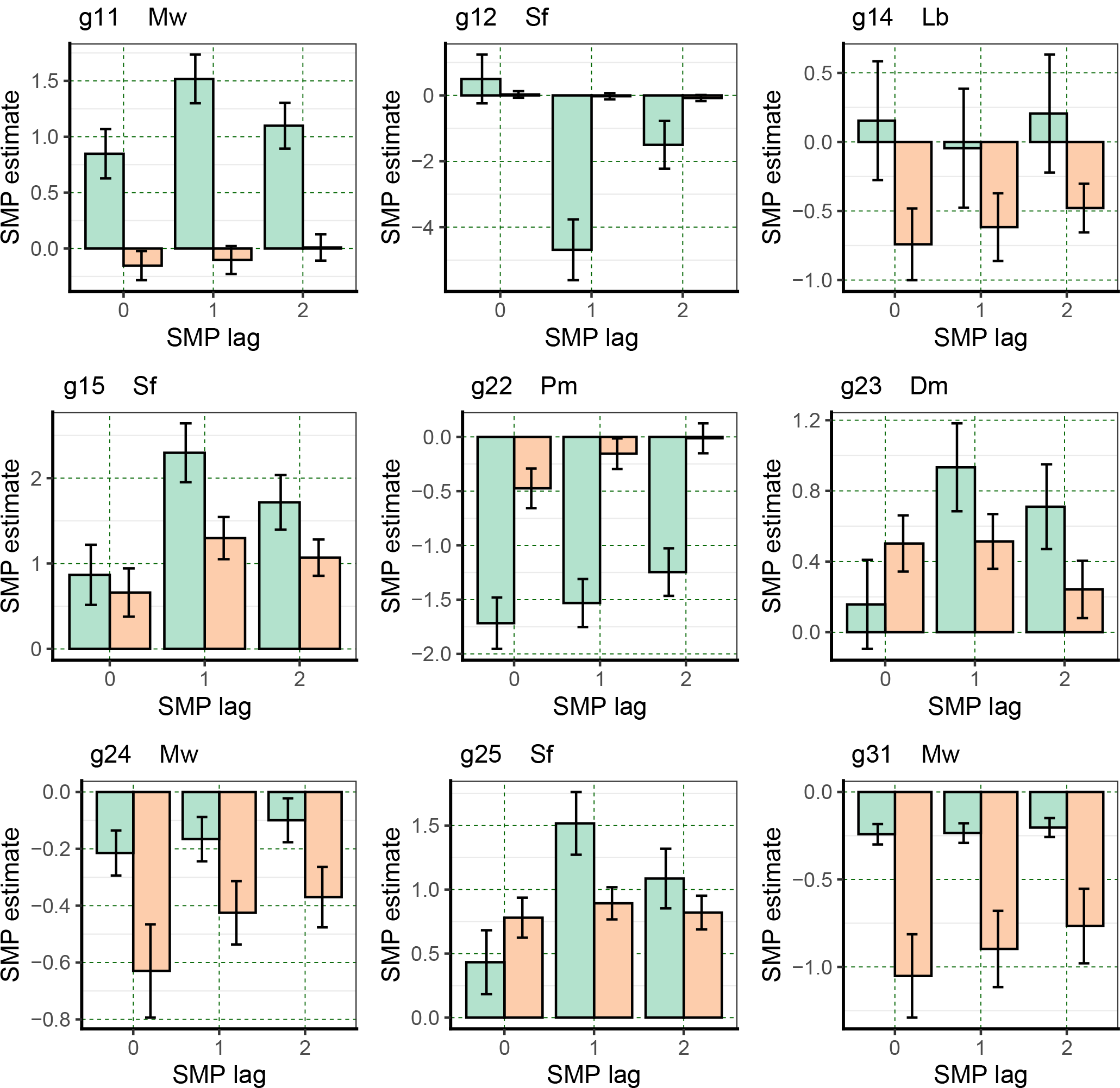
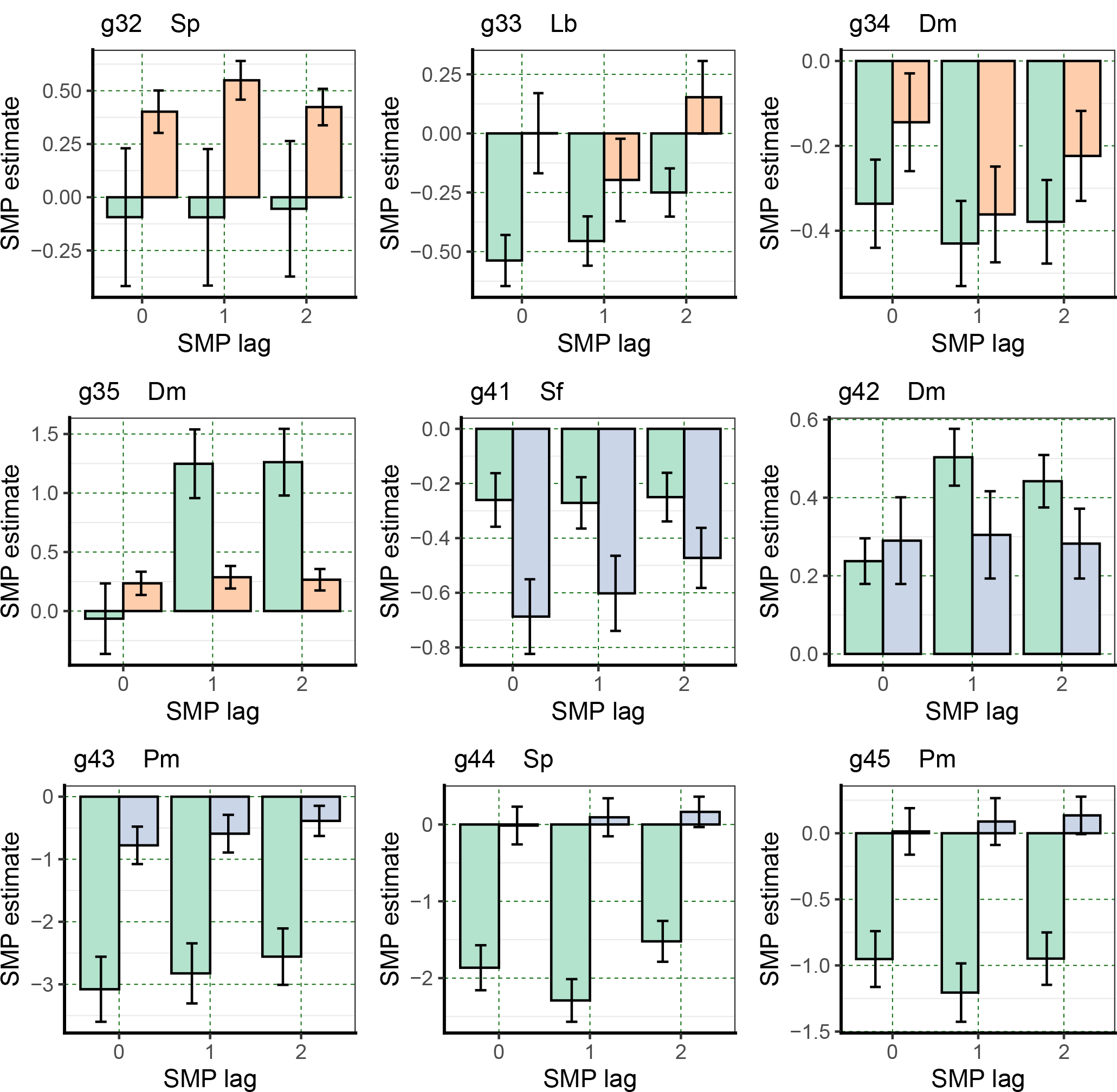
Mean ± SE coefficients (slopes) of *gthi* regressed on soil moisture potential, SMP, at 0-, 1- and 2-day lags, in the wet (green bars) and dry (orange bars) periods for the 18 trees taken for the time-series analyses. There were no dry-period time-series estimates for station 4, but these are predicted here (blue bars) for lag 1 from the species’ relationship to maximal diurnal girth change. Bands are listed in Table 4 with individual tree codes.

**Table 4.** Mean adjusted coefficients for Dimorphocalyx muricatus (Dm) and Mallotus wrayi (Mw) from the time series GLS regressions of 1-day girth increment (gthi) on soil moisture potential lagged by 1 day (SMP_−1_) and unlagged ambient temperature (TEMP_0_), in the wet and dry periods. Individual replicate tree values are plotted in Fig. 7 which displays the species’ ranges.

| variable | species | n | wet | n | dry |
| --- | --- | --- | --- | --- | --- |
|  |  |  | coeff. |  | coeff. |
| SMP<sub>-1</sub> | Dm | 4 | 0.77 ± 0.47 | 3 | 0.23 ± 0.38 |
|  | Lb | 2 | -0.33 ± 0.26 | 2 | -0.62 ± 0.37 |
|  | Mw | 3 | 0.62 ± 0.92 | 3 | -0.70 ± 0.34 |
|  | Sf | 3 | 2.04 ± 1.36 | 2 | 1.88 ± 0.53 |
| TEMP<sub>0</sub> | Dm | 4 | 1.77 ± 0.59 | 3 | -0.40 ± 0.70 |
|  | Lb | 2 | -0.04 ± 1.15 | 2 | 1.58 ± 3.07 |
|  | Mw | 3 | 1.08 ± 0.22 | 3 | 2.76 ± 1.41 |
|  | Sf | 3 | 2.69 ± 0.89 | 2 | 1.94 ± 0.27 |

Girth increment dependence on temperature was similarly analyzed using current logger temperature (TEMP_0_), and the lagged values TEMP_−1_ and TEMP_−2_. It was clear that highest significances were reached with TEMP_0_, with 13 trees (12 positive, one negative), yet TEMP_−1_ had only four cases (all negative), and TEMP_−2_, just two weakly (one each positive and negative) (Fig. 5; Appendix 5: Table S2). Interestingly, for at least those significant, the relationship was largely with opposite slope using TEMP_−1_ than it was using TEMP_0_. For the dry period, five trees (one negative, four positive) were significantly related to TEMP_0_, and five also with TEMP_−1_ (all but one negative) yet with lower significances, and then six (two negative, four positive) with a return to higher significances, for TEMP_−2_. Switching of slope direction was less consistent in the dry than it was in the wet period. Hence, the most significant effects were concentrated on TEMP_0_, but with some trees having 2-day lag effects in the dry period (Appendix 1: Table S3).

**Fig. 5.**
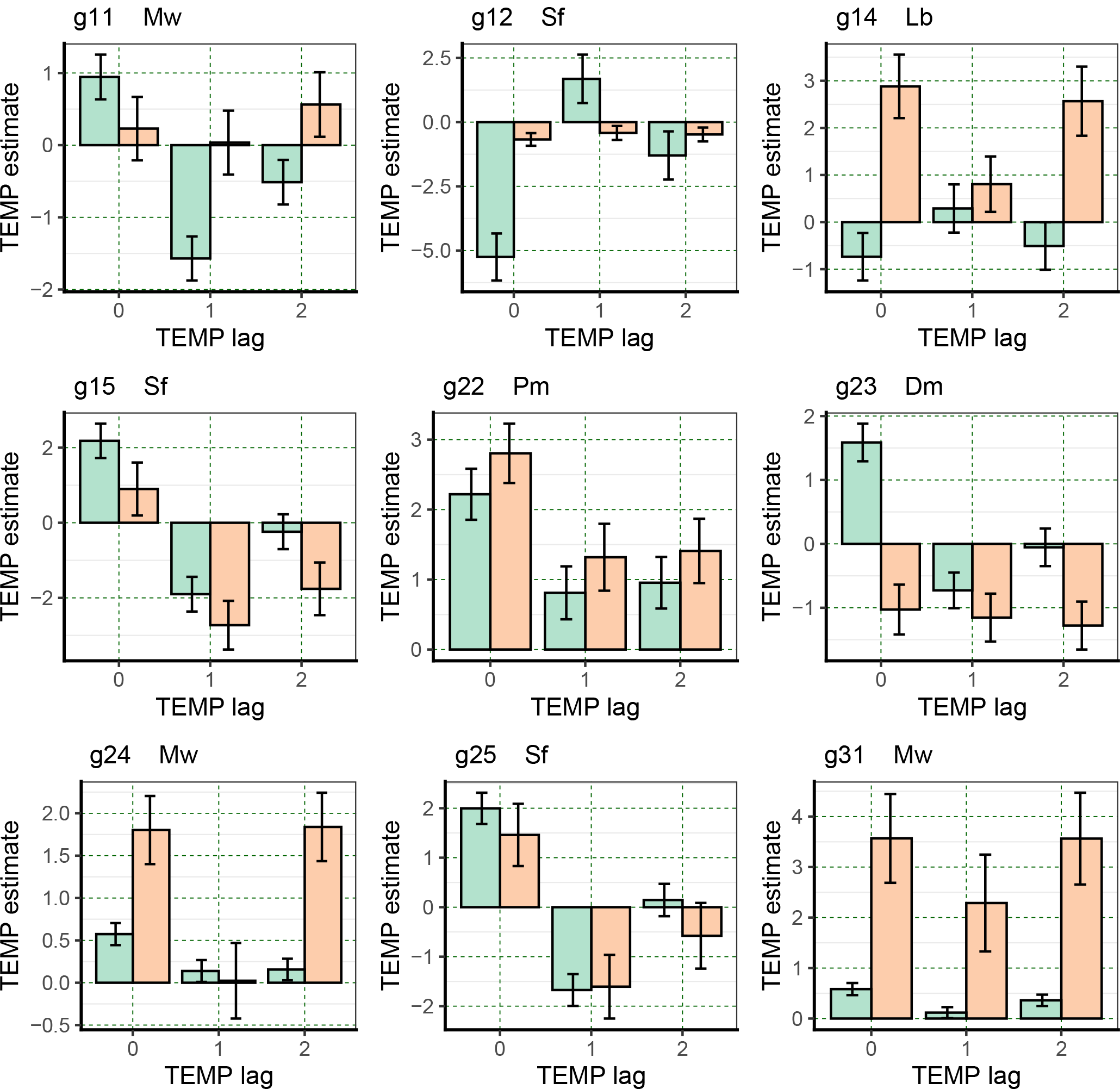
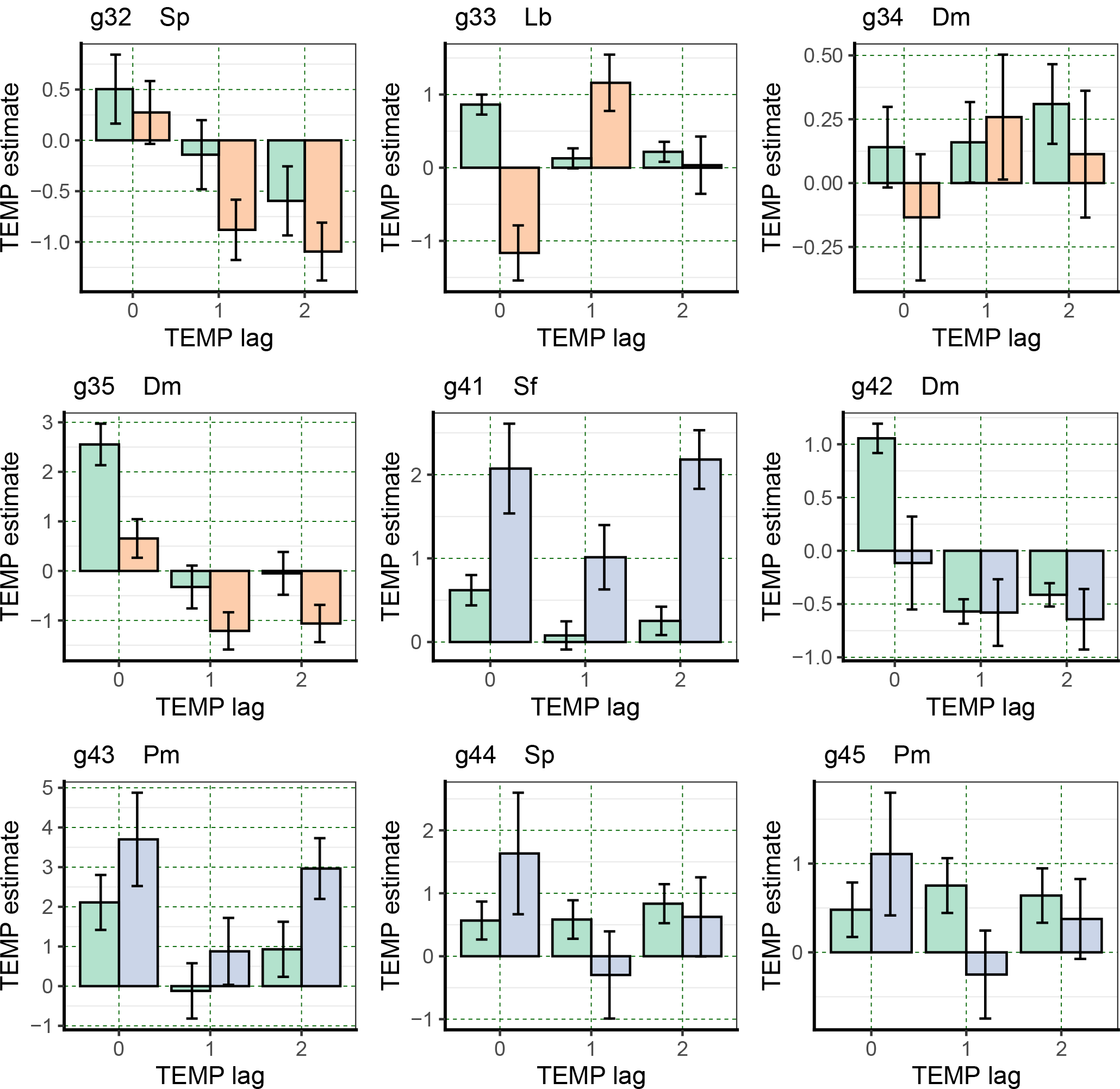
Mean ± SE coefficients (slopes) of *gthi* regressed on logger temperature, TEMP, at 0-, 1- and 2-day lags, in the wet (green bars) and dry (orange bars) periods, for the 18 trees taken for the time-series analyses. There were no dry-period time-series estimates for station 4, but these are predicted here (blue bars) for lag 0 from the species’ relationship to maximal diurnal girth change. Bands are listed in Table 4 with individual tree codes.

The two-term regressions support the one-term ones with the most significant slopes being for SMP_−1_ and TEMP_0_, with SMP_0_ and TEMP_−1_ being important in wet only. In the wet and dry periods, *t*-values for the SMP coefficients were strongly correlated across columns, i.e. across the four combinations of SMP and TEMP lags (Appendix 5: Table S3), especially when SMP_−1_ was involved (Appendix 6: Fig. S1): six correlations per period with *r* = 0.81 to 0.99, *P* < 0.001. For temperature, however, in the wet period, when the TEMP-lag was the same (0 or 1), correlations were positive and strong (*r* = 0.926 to 0.994, *P* < 0.001), but not otherwise when the correlations became strongly negative. This testifies to an alternating effect from day to day. In the dry there was the same positive significance for same-lag cross-correlations, but the negative different-lag ones were fully lost.

Dry period significances were often not high, although more often so for SMP_−1_ and TEMP_0_. The slopes for temperature in the two-term regressions (Appendix 5: Table S4) showed more significant outcomes when potential was SMP_0_ or SMP_−1_, but the temperature was TEMP_0_ (Appendix 6: Fig. S2). Agreement across the columns was very high, and for the dry period a similar number of slopes were significant, as in the one-term regressions for SMP_0_/TEMP_0_ and SMP_−1_/TEMP_0_. Tests of significance for SMP*TEMP interactions were almost completely non-significant in the wet period (Appendix 5: Table S5), but in marked contrast several interactions were significant in the dry period (up to six, of mixed sign), in the three combinations which were *not* SMP_−1_ and TEMP_0_ (Appendix 5: Table 5B, columns 1-4). Hence, the two-term regressions affirmed the one-term ones, with SMP_−1_ and TEMP_0_ being most important, and their having negligible interactions. It implies that SMP_−1_ and TEMP_0_ were acting quite independently.

The sizes of the regression coefficients in the wet and dry periods do not necessarily match the relative sizes of the *t*-values since the dry had larger SEs than the wet period.

Guided by the most significant outcomes from the regressions, i.e. focusing on those that used SMP_−1_ and TEMP_0_, wet and dry period slopes may be compared for each tree in Figure 4 in turn. Starting with SMP_−1_ (centre bars of each panel), the majority showed similarly positive or respective negative estimates across the two periods. The two exceptions in Figure 4 are g12 and g33 (marginally also g35). Although the effects were strongest for a lag of 1 day they were not absent for lags 0 and 2 days. The ± 1 SE bars give an approximate indication of strengths of differences between periods but, since the same tree is considered in both periods wet and dry period, the data are not fully statistically independent, even though within period temporal autocorrelation was accounted for by arima modelling. Eleven of 13 trees had wet and dry period estimated slopes in the same direction for SMP_−1_ (i.e. both positive or both negative) (Fig. 4), even though for the opposite two cases (g11, g32) the minor estimates were very small and with large SEs. For nine trees the wet slope estimate was larger than the corresponding dry one, and in four cases the converse. In summary, there was no overall common differential response to period. For g11, g12, g22, g25, g31, g32 and g35 (i.e. ∼ half of the trees) the difference was towards significance. The slopes for SMP_0_ and SMP_−2_ generally showed less strong differences between the periods than did SMP_−1_. These relative SMP_−1_ slope values will be returned to later. The five trees at station 4, which lacked dry period estimates, showed much more consistent estimates across SMP_0_ to SMP_−2_.

Taking the values of TEMP_0_ slopes (left bars in the panels of Fig. 5) for reference, lags and wet-dry differences in mean *gthi* can be compared too for the girth response to temperature. Compared with SMP, there are many more switches in slope directions (sign of estimates) between TEMP_0_ and TEMP_−1_ and TEMP_−2_. Values that were often positive for both wet and dry periods became negative; occasionally, this was in the opposite way (for g12 and g33). Differences between wet and dry period slopes were also very variable. For g15, g22, g25, g32 and g34 they were similar (i.e. likely not significantly different), but the other bands showed a mixture of ‘wet > dry’ and ‘dry > wet’ estimates (Fig. 5). Switching from positive to negative sign between periods was obvious for g23 and g33, and negative to positive for g14, g24 and g31. The SEs on TEMP coefficients were similar to those for SMP. Compared to the SMP, the five trees at station 4 had a lower consistency across lagged TEMP estimates.

Graphing the mean coefficients for *gthi* depending on SMP, and on TEMP, from the two-term regressions in a factorial SMP_0_-SMP_−1_ x TEMP_0_-TEMP_−1_ arrangement for each tree in Figs S1 and S2 of Appendix 6 illustrated a high degree of correspondence across the four combinations of the variables, except where significant interactions occurred. With the focus still on SMP_−1_/TEMP_0_, the key bars are the second and third sets (paired wet and dry) in Appendix 6: Figs S1 and S2 respectively. Notable deviations occurred for g25, g33, g35 and g42 (wet only) for SMP coefficients, and marginally just g35 and g44 (wet only) for the TEMP ones. Nevertheless, analysis of variance for all trees (*scl* = 1 and 2) (analyses run in the same way as for *gthch*) showed no significant differences between species in their *gbh*-scaled coefficients (*P* = 0.33 - 0.93) for the two variables in either period. Considering the small trees (*scl* = 1), again significances were a little better, yet only marginal for SMP_−1_ in dry period due to `Sf` having the largest positive value (*P* = 0.066 to 0.67; Fig. 7b or 10a for instance). These comparisons give support to concentrating on comparing tree sizes and species identities between wet and dry periods for essentially SMP_−1_/TEMP_0_.

Under the Granger causality test, SMP was highly significant for many more trees than was TEMP in the wet period (*F*-values for p = 2 in Appendix 1: Table S4): 15 for the former, and six for the latter variable, out of 18 at *P* ≤ 0.01. In the dry period significant effects were likewise frequent for SMP and rarer for TEMP: nine and four out of 13 respectively. On this basis, SMP would appear the commoner and stronger predictor of *gthi* than TEMP.

### Variation in girth increment with tree size

Girth increment, *gthi*, varied widely across the banded trees in both wet and dry periods in terms of means and standard deviations (Appendix 1: Table S2). The quantiles indicated that positive and negative values were roughly equally distributed in the wet period, but in the dry one they were more positively skewed. Although *gbh* was a major factor in determining *gthi*, there was still considerable variation in mean and SD of *gthi* among the small trees (*scl* = 1; Fig. 6). Mean and SD of *gthi* were positively correlated in the wet period (*r* = 0.482, df = 16, *P* ≤ 0.05) but not significantly correlated in the dry one (*r* = −0.272, df = 11, *P* = 0.37). Large trees (in the overstorey) are well separated from the small ones (in the understorey) in Fig. 6. When just the small ones are considered, these form a cluster with mean *gthi* close to zero in the wet period (Fig. 6a) but with much variation in the SD of *gthi* (Fig. 6b). Contrastingly, in the dry period the spread in mean *gthi* was quite obvious (Fig. 6c) and the corresponding SD values similarly so (Fig. 6d). Differences and variation in responses among trees were more marked in the dry than wet period. As expected, simple-difference absolute growth rates (*agr*, mm/year) of the banded trees over wet and dry periods were almost perfectly linearly related to mean *gthi* (slight variation due to shrinking/swelling on the first and last date of period): *agr_W_* = -0.09 + 3.64 *gthi*_mn_ for the wet, and *agr*_D_ = 0.25 + 3.78 *gthi*_mn_. The difference in intercept from zero in the dry period reflects the negative skew in the *gthi*-values.

**Fig. 6.**
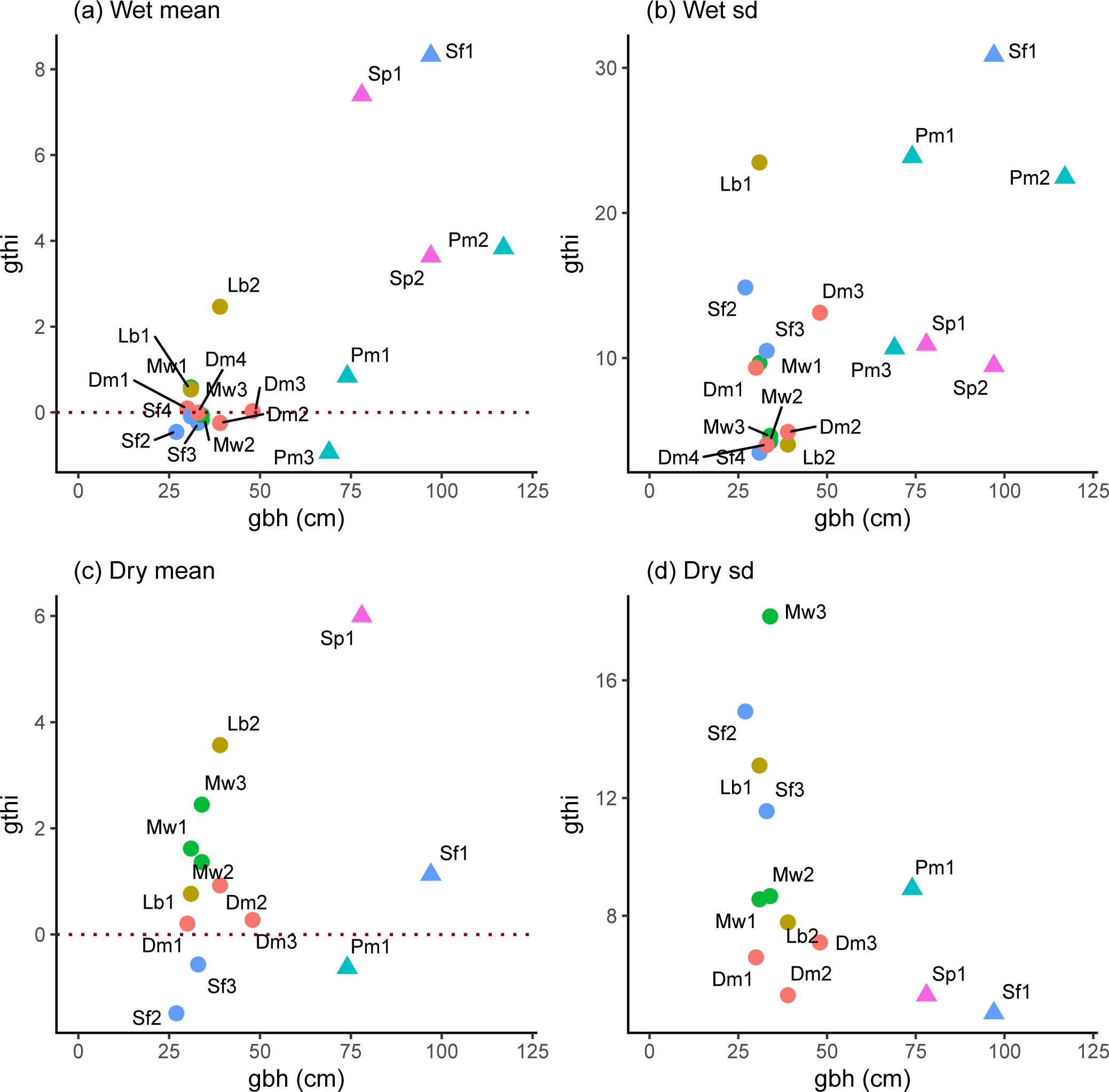
Mean and standard deviation (sd) of daily girth increments (*gthi*), plotted against stem girth (*gbh*) in the wet (a, c; *n* = 18) and dry (b, d; *n* = 13) periods. Species symbols are color- coded, their abbreviations as labels with the individuals distinguished by numbers, are listed in Table 2. Size classes: *scl* 1, circles; 2, triangles.

Absolute values of the coefficient of variation (SD/mean) *gthi* were generally higher in the wet (1.5 – 375) than dry (1 – 25) period, for the 13 trees recorded in both (Appendix 1: Table S2). This implies that the contribution of growth to *gthi* compared with water-flux related fluctuations not only varied very considerably (relatively much when the ratio was low, conversely little when it was high), but more so in the dry than wet even though the dry created larger fluctuations. Across these 13 trees in the dry period, however, mean mean-*gthi* (and SE) were 1.47 ± 0.82 and 1.20 ± 0.54 cm·10^−3^ in the wet and dry periods respectively.

Based on this limited sample, growth rates overall barely differed between periods. The results highlight again the considerable variability between individuals in their growth rates and responses.

### SMP and rainfall

Since *n*-d rainfall totals were positively correlated with SMP_−1_ (Table 3, variables sqrt-transformed), using both as predictors in a regression may have involved some collinearity.

Using the original dry-period *gthi*-values, i.e. not those already adjusted with the daily rainfall data, cross-correlations of estimates of the SMP_−1_ coefficients from regressions of *gthi* on SMP_−1_ alone (the original model) and those from regressions now of *gthi* on SMP_−1_ and *rft* (using the 10-d and 20-d values for wet and dry periods respectively), were *r* = 0.406 (df = 16, *P* = 0.095) in the wet, and *r* = 0.661 (df = 11, *P* = 0.014) in the dry, period. Their corresponding *t*-values were better correlated (*r* = 0.771 and 0.671, *P* < 0.001 and 0.012). Using the residuals from a first linear regression of SMP_−1_ on *rft* led to almost identical coefficients and *t*-values.

In the wet period slopes of *gthi* on 10-d-*rft* were significant for just one band, g31 (*P* < 0.01, Bonferroni corrected α’ = α/18), and in the dry period the same on 20-d-*rft* for two bands g32 and g34 (*P* < 0.05, α’ = α/13 resp.). Comparing SMP_−1_ estimates from GLS-arima model fits with and without the corresponding wet- or dry-period *rft*-term, now omitting the four bands which showed volatility and needed GARCH adjustments, the relationship became approximately linear especially when the estimates were both negative (Appendix 1: Fig. S4). However, where the single-term model estimate of SMP_−1_ was positive, that from the two-term model deviated considerably from the line, more so in the wet than dry period (Appendix 1: Fig. S4a, b). Band g12 would qualify as an outlier in Fig. S4a; and it was noted before as having unusual lag patterns in the wet period (Fig. 4). Bands g12 and g33 had likewise unusual patterns in the dry period, and in earlier regressions *gthi* was correlated atypically with ln(rainfall) on the day before and not the current day. Leaving aside these unusual bands, concave curves fit the points quite well (Appendix 1: Fig. S4c, d). When *rft* was included in the GLS models, slopes of regression of *gthi* on SMP_−1_ that were negative in the single-term GLS fit became more negative, and those that were positive, more positive. The gradient where the estimate of SMP_−1_ in the single-term model was equal to 0 was 4.29 in the wet, and 3.35 in the dry, period.

### Relationships between coefficients and descriptor variables

Standardized SMP_−1_ and TEMP_0_ coefficients were positively and strongly significantly correlated in the wet period (*r* = 0.811, df = 16, *P* ≤ 0.001) and it is clear from Fig. 7a that small trees (*scl* = 1) had mostly positive, and large trees (*scl* = 2) all had negative SMP_−1_ values. In the dry period (*gthi* as rain-regression residuals) the relationship was negative but weaker and non-significant (*r* = –0.348, df = 11, *P* = 0.24); the three large trees occurring near the centre in among the small-tree SMP_−1_-and TEMP_0_-values (Fig. 7b).

**Fig. 7.**
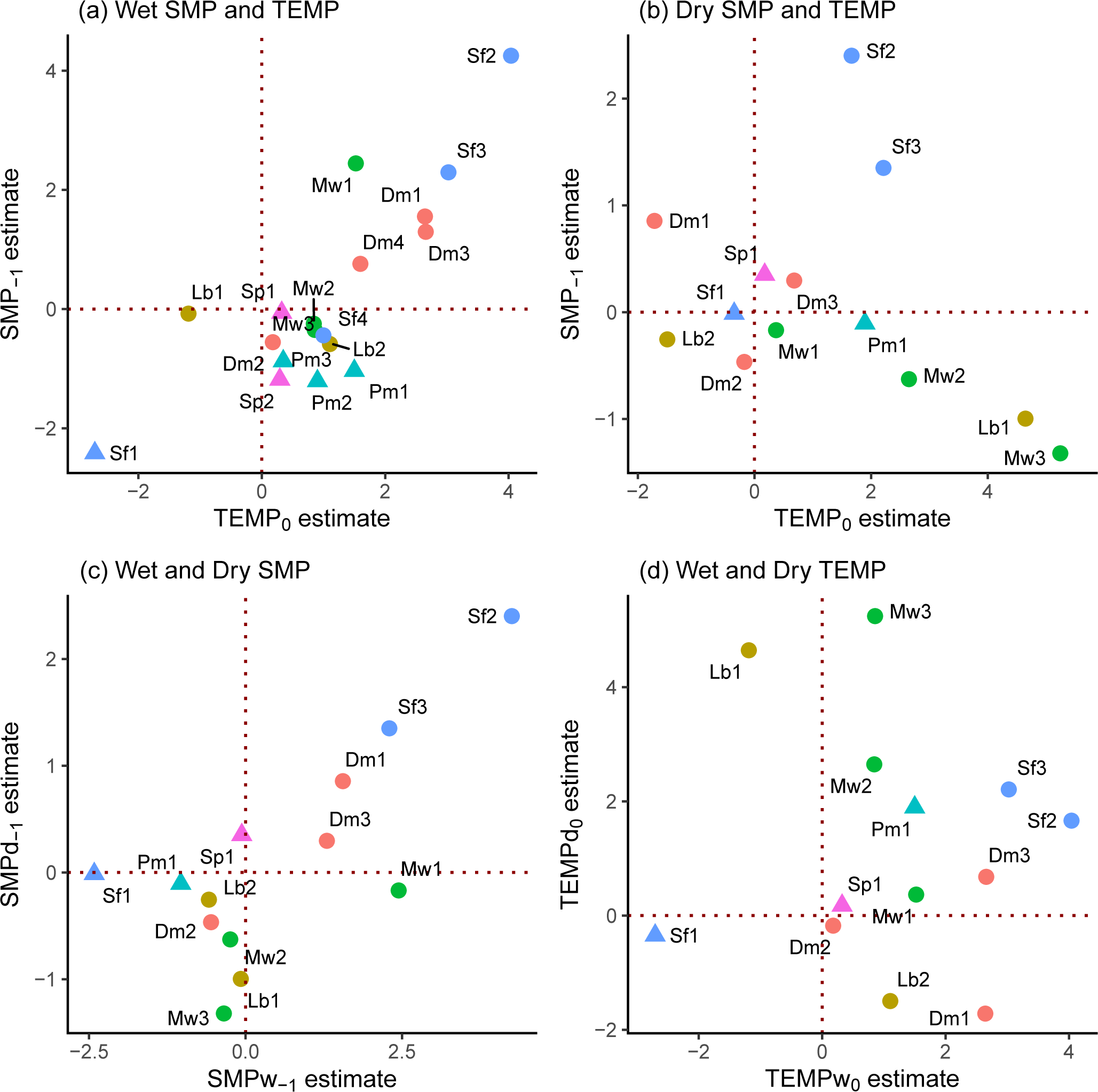
Standardized SMP_−1_ versus TEMP_0_ coefficients graphed together for (a) wet [w] and (b) dry [d] periods for the 18 respective 13 trees analyzed for their *gthi* time-series, and their corresponding dry- and wet-period (c) SMP_−1_ and (d) TEMP_0_ values against one another. Symbols as in Fig. 6.

Considering just the small trees (*scl* = 1), in the wet period the correlation between SMP_−1_ and TEMP_0_ was very similar and significant to using all trees (*r* = 0.809, df = 10, *P* = 0.001), yet no better in the dry period (*r* = –0.356, df = 8, *P* = 0.31). SMP_−1_-dry and SMP_−1_-wet were positively correlated (*r* = 0.716, df = 11, *P* ≤ 0.01; Fig. 7c) but this was not the case for TEMP_0_-dry and TEMP_0_-wet (*r* = –0.074, df = 11, *P* = 0.81; Fig. 7d). Again, taking just the small trees these last two relationships became respectively more significant and less insignificant (*r* = 0.869, df = 8, *P* ≤ 0.001; *r* = –0.347, df = 8, *P* = 0.33). The correlations between SMP_−1_ and TEMP_0_ involved species that were each represented by two to four trees, so there may have been some phylogenetic effect in trait terms, points for the same species being not fully independent of one another. However, the replicate trees within species were so well-dispersed that any effect would have been very small.

Canopy status, CanStat, was strongly and significantly positively correlated with *gbh* (*r* = 0.917, *P* < 0.001, df = 16; CanStat = 1.426 + 0.02761 *gbh*), and on the basis of trees divided into the two *scl* classes (under- and overstorey), the ∼ 60 cm *gbh* threshold was equivalent to a CanStat value of 3.08. Canopy status is used further here, rather than *gbh* per se, because as a composite measure this first variable probably reflects much better the light levels directly above crowns and, therefore, indirectly the evapotranspiration as radiation and thus temperature changed. CanStat was strongly positively correlated with absolute stem growth rate in the third main-plot census period P3 (2001-2007), *agr*_P3_ (*r* = 0.668, df = 16, *P* = 0.002) but less well with the corresponding relative stem growth rate, *rgr*_P3_ (*r* = 0.400, df = 16, *P* = 0.10).

SMP_−1_ was strongly negatively correlated with CanStat in the wet period (*r* = −0.688, df = 16, *P* < 0.002), but not in the dry one (*r* = −0.139, df = 11, *P* = 0.65) (Fig. 8 a, b). For TEMP_0_ in the wet and dry periods correlations were similar (*r* = −0.628, *P* ≤ 0.01; and *r* = −0.194, *P* = 0.53 respectively; Fig. 8 c, d). For small trees only, SMP_−1_ was again correlated negatively with CanStat in the wet period (*r* = −0.530, df = 10, *P* ≤ 0.10), and also in the dry one (*r* = −0.451, *P* ≤ 0.05, df = 8); and it was so for TEMP_0_ in the wet (*r* = −0.610, df = 10, *P* ≤ 0.05) but not in the dry (*r* = 0.076, df = 8, *P* = 0.83). It is interesting that the strong negative correlation of SMP_−1_ and TEMP_0_ with CanStat, occurring in the wet but *not* dry period for all trees, was shown again especially for SMP_−1_ in the dry period for small trees.

**Fig. 8.**
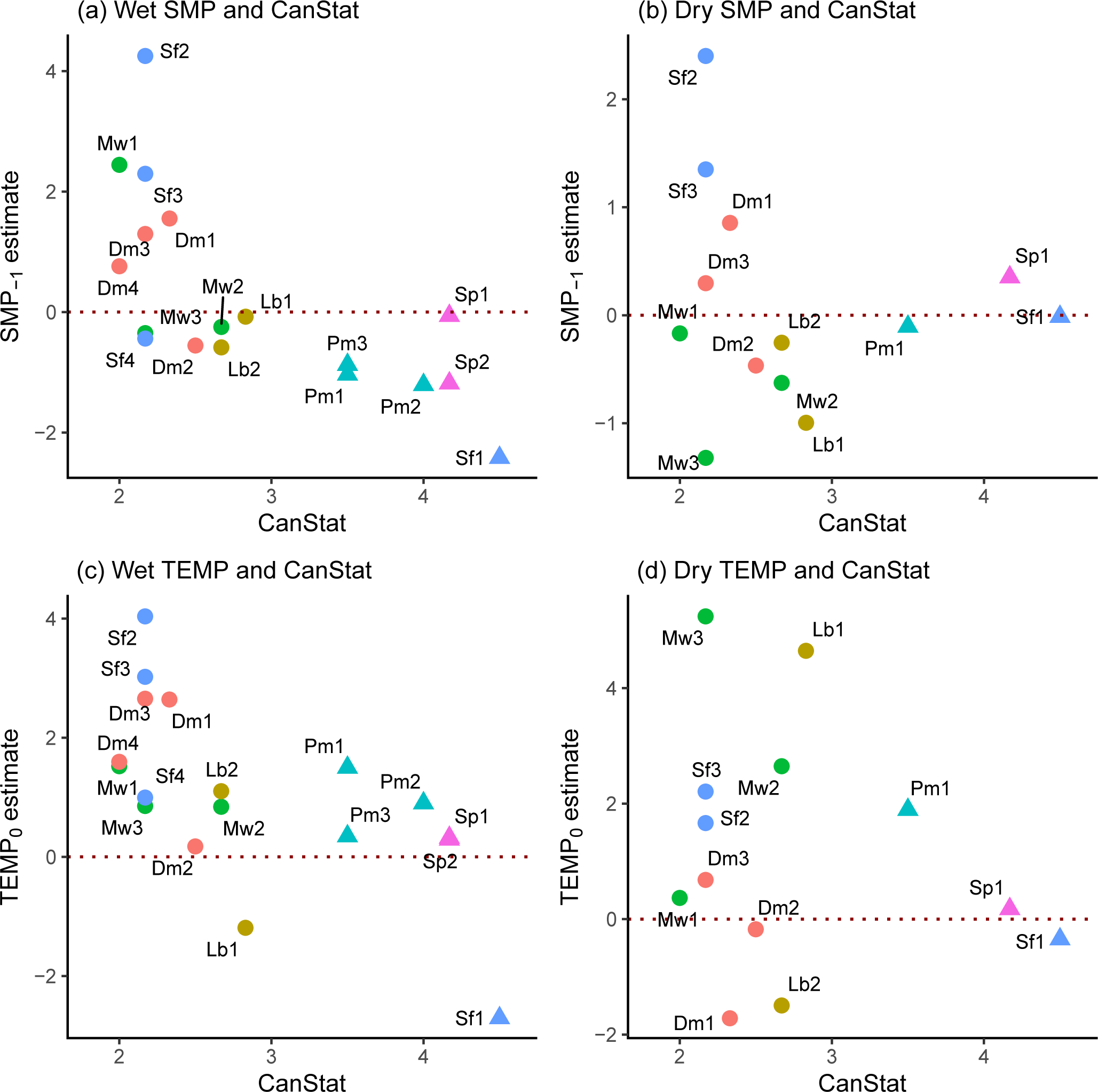
Standardized SMP_−1_ and TEMP_0_ coefficients plotted in relation to canopy status (*CanStat*), in (a, c) wet (*n* = 18), and (b, d) dry (*n* = 13), periods. Symbols as in Fig.6.

Neither the *gbh*-scaled coefficients of SMP_−1_ nor of TEMP_0_ in the wet and dry periods differed significantly between stations (*F* = 0.10 to 0.88, df = 3,14 or 2,10; *P* > 0.45), although mean SMP_−1_ coefficient did decline the most from 1.05, 0.64, −0.05 down to −0.59 between stations 1 to 4 in the wet period (Appendix 1: Fig. S5). In the corresponding dry period mean SMP_−1_ coefficients were 0.31, 0.37, and −0.28 across stations 1 and 3. There was therefore some tendency for SMP coefficients to be more negative on the ridge (stations 3 and 4) than the lower slope stations (1 and 2).

### Relationships between coefficients and diurnal changes

A large variability, not only with respect to tree size (*gbh*, *CanStat*, or *scl*) but also within-species — even where tree sizes were similar, is shown in the relationships between the two time-series coefficients (*gbh*-scaled) and *gthch* statistics (Figs 9 and 10). Since ‘replicate’ trees were scattered widely across the panels of these figures, phylogenetic influence is likely to be very low, and a high level of tree individuality is indicated.

**Fig. 9.**
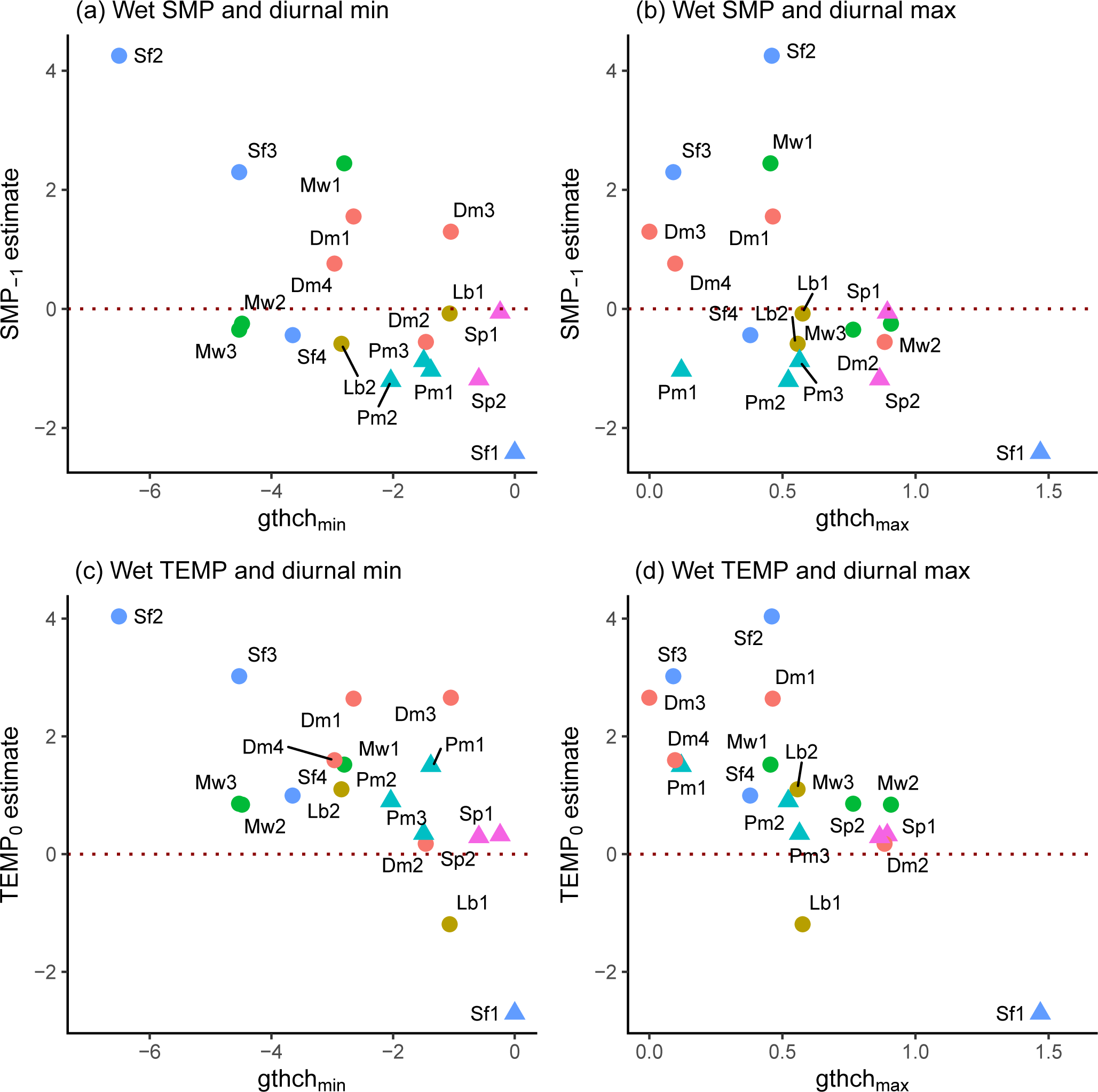
Standardized SMP_−1_ and TEMP_0_ coefficients in the wet period in relation to the corresponding (a, c) minimum and (b, d) maximum diurnal girth changes (*gthch*) of the periods (*n* = 18). Symbols as in Fig.6.

**Fig. 10.**
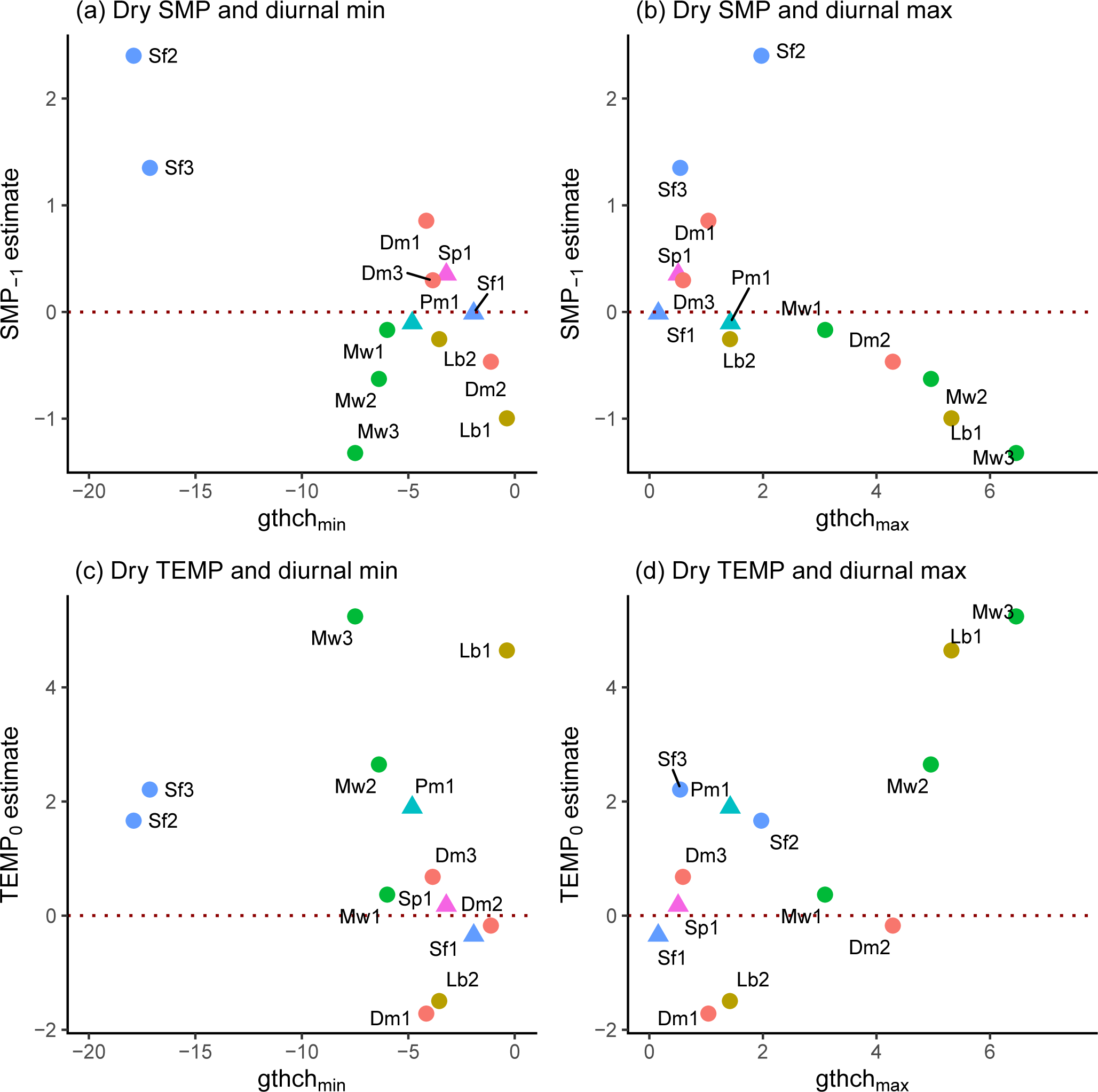
Standardized SMP_−1_ and TEMP_0_ coefficients in the dry period in relation to the corresponding (a, c) minimum and (b, d) maximum diurnal girth changes (*gthch*) of the periods (*n* = 13). Symbols as in Fig.6.

Correlations were negative and significant for SMP_−1_ (*r* = –0.640, *P* ≤ 0.01; Fig. 9a) and TEMP_0_ (*r* = –0.658, *P* ≤ 0.01; Fig. 9c) versus *gthch_min_* in the wet period, though notably the three trees of *S. fallax* were the most outlying graph points; likewise, for SMP_−1_ (*r* = –0.523, *P* ≤ 0.05; Fig. 9b) and TEMP_0_ (*r* = –0.735, *P* ≤ 0.001; Fig. 9d) versus *gthch_max_*.

In the dry period, the relatively strong correlation for SMP_−1_ versus *gthch_min_* (*r* = – 0.715, *P* ≤ 0.01; Fig. 10a) was due in part to the negatively skewed values for the two small trees of *S. fallax,* and was also the case for TEMP_0_ versus *gthch_min_* (*r* = –0.229, *P* = 0.45; Fig. 10c). By contrast, whilst the correlation for SMP_−1_ versus *gthch_max_* was strong (*r* = –0.645, *P* ≤ 0.02; Fig. 10b; note the outlying `Sf2`), and for TEMP_0_ versus *gthch_min_* it was equally strong but now positive (*r* = 0.683, *P* = 0.01; Fig. 10d). Considering just small trees (*scl* = 1), correlations of SMP_−1_ and TEMP_0_ with *gthch_min_* and *gthch_max_* were very similar despite the smaller sample size. The contrast between the relationship of TEMP_0_ with *gthch_max_*, in both wet and dry periods, is remarkable: and the more so that this was little changed when using only small trees, especially of true understorey species. In Figs 9 and 10, the graphed coordinates of the replicate trees of the same species were not close to one another, in either period; the species being well mixed within panels.

Average SMP_−1_ coefficients decreased between the wet and dry periods, twice as much for *D. muricatus* as for *M. wrayi* (Table 4). Those for TEMP_0_ decreased even more for *D. muricatus* though hardly at all for *M. wrayi*. For both variables the replication at the tree level is too low for formal tests of significance, although bearing in mind the significance of each estimated coefficient, there is an interesting indication that *D. muricatus* was more resistant to the drying than *M. wrayi*. One difficulty was that among the three to four trees of these two species there was one in each case each that behaved quite differently from the others.

### Prediction of failing time-series estimates

No estimates from time series regressions were possible for five trees in the dry period at station 4. This led to an unfortunate imbalance when comparing trees and species across the wet and dry periods. To partly compensate for this lack, SMP and TEMP coefficients (for lags 0, 1 and 2 days) were predicted from their relationships with *gthch_max_* (Fig. 10b, d), since these had the better and more robust correlations of *gthch_min_* and *gthch_max_* (reported above).

Simple linear regressions allowed the missing trees’ coefficients to be predicted (with SEs; Appendix 1: Table S5). The regressions used the *gth*-scaled values of SMP and TEMP estimates, so predictions required back-scaling. The new values were incorporated into Figs 4 and 5. The regressions for SMP had similarly high negative fits: those for TEMP were all positive but a little less significant. Important to note is that the SEs on the GLS-arima- derived estimates are different in origin from those involving also the extra step of *gthch-* coefficients regressions. The same approach was applied for the two-term GLS-arima models (no interaction term) for SMP and TEMP estimates with 0 and 1-day lags and these likewise added into Appendix 6: Figs S1 and S2 (regression fits in Appendix 1: Table S5).

### Influence of local tree basal area abundance

For each of the 12 small (*scl* = 1) trees, a measure of neighbourhood competition is the sum of basal areas (*BA*) of all trees ≥ 10 cm *gbh,* within a 5-m radius (*d*) and with each tree’s BA inversely weighted by its distance (d) to the focus one (*BA/d,* cm_2_/m). Relevant estimates were abstracted from calculations using the 2007 main plots census data (Newbery and Ridsdale 2016). In neither the wet nor dry periods were coefficients of SMP_−1_ and TEMP_0_ significantly correlated with the BA/d-values (*P* = 0.22-0.47). At 5-m radius trees made *BA/d* =167-2684 cm_2_/m (two high outliers: g42 [`Dm4`] and g25 [`Sf3`] of 1286 and the maximum cm_2_/m). To compare the effects of *BA/d* on species in a balanced way, data for just small trees (*scl* = 1) which had *BA/d* ≤ 1500 cm^2^/m, and at least two trees in this range were taken. Up to 500 cm^2^/m all SMP_−1_ and TEMP_0_ coefficients consistently steeply increased in the wet period, but then variably dropped away at > 500 cm^2^/m. Larger BA/d-values most likely resulted from larger trees with deeper less-overlapping root zones. In the dry period there was also an increase for SMP_−1_ with *BA/d* < 500 cm_2_/m, but for TEMP_0_ a completely opposite trend, i.e. the coefficient decreased with increasing *BA/d* (Appendix 1: Fig. S6).

## DISCUSSION

### Dendrobands, bark and microclimate

Barks of moist tropical forest trees are typically thin, hard, smooth and light-colored, especially those in the shaded understorey (Richards 1998). Thickness varies mostly between 3 and 8 mm for trees 50-100 cm gbh (Roth 1981, Hegde et al. 1998, Paine et al. 2010, Poorter et al. 2014, Rosell 2016). Occasionally thicker barks do occur especially on larger trees. The outer bark, or rhytidome, is non-living material, being composed of one or more cork (phellum) layers (Evert et al. 2006), and has the main functions of water-proofing the stem, providing defense against pests and pathogens, and protecting the inner living bark from mechanical damage. The outermost layers (epidermis) slowly decompose, crumble and fall away, more often as very fine material. It is this outer dead material that can absorb some humidity and possibly affect dendroband recordings. This partial effect was well catered for by the statistical analysis in this paper. It is essential to realize, moreover, that in the main tropical small-tree bark dimensions are very different in scale from the very thick and rough barks of many oaks and pines studied in very dry habitats of the temperate biome (e.g. Zweifel et al. 2005, Mencuccini et al. 2017, Oberhuber et al. 2020). Trees in the shade tend to have thinner barks than those in the open in general (Thomas 2000). Roth (1981) recorded cork layer widths of only 0.1 to 0.5 mm on many trees in Venezuelan rain forest: in moist environments, thin bark facilitates better aeration outwards to the lenticels than thick bark. In contrast to understorey species, small trees of the Dipterocarpaceae begin to form fissured thicker barks with their many periderm layers, ones that become clear features of the adult canopy trees (Whitmore 1962).

During the course of the day, in marked contrast to the upper forest canopy, RH changes very little at about 1 m above the forest floor in the understorey. It may fall from ∼100% at night to 85-90% in the mid-afternoon in continuously wet periods (Cachan 1963, Aoki et al. 1978, Hardwick et al. 2015, Song et al. 2021). Comparable data for short dry periods appear not to be available in the literature. Otherwise, useful RH profiles for primary forest are scarce, largely because the data often come from large towers in clearings, e.g. at BCI in Panama (Herrmann et al. 2016), or secondary forests which are more open.

Unfortunately, no RH data were collected for the Danum site in the forest understorey for both wet and dry periods. The data from the climate station in the large DVFC clearing outside the forest showed falls in mid-afternoon RH to ∼70% in dry periods, but it is more likely that the closed forest (with very few tree fall gaps) substantially buffered this change and RH probably remained higher within the understorey. This contention was supported by RH falling from over 90% to close to 80% in the dry seasons of three years in lowland rain forest understorey in Panama (Wright 1991). Longman and Jenik (1987) showed for clearings versus understorey in Ghanaian forest, RH fell from 90 to 60% and 90 to 80% respectively in the afternoons. Tymen et al. (2017) established by a broad sampling of understorey microclimatic conditions of forest in French Guiana, that minimum RH was 25% lower in clearings than in the closed understorey, and furthermore this ratio altered very little across seasons. Hence, at Danum when minimum RH reached 60% outside (Appendix 2: Table S1) the Guiana study would predict a minimum of 80% RH around the dendrobands used inside the forest, both in wet and dry periods on hot days. In the present study growth across wet and dry periods was not considered as e.g. in the study of Stahl et al. (2010). Once the 30-day rft had moved below 100 mm, into the dry period, *gthi* variation was being measured *within* that period. Lastly, it must be recognized that water will to some degree be moving radially from an outer shower-wetted bark to the inner living bark (Mencuccini et al. 2013), and the dynamics of water flux at these times is very intricate. Accordingly, the statistical analysis in the present paper pointed to RH change not having a significant effect on dendroband recordings made *within* each period.

Rainfall at Danum is most frequent in the mid- to late-afternoon. Chappell et al. (2001) found that 43% of diurnal rainfall fell between 14:00 and 16:00. This would have been just after the time of day when perhaps a tree’s bark had shrunk and a shower would then counteract the direction of *gth* change. Much would depend on the rate at which bark shrinks and swells within a day and this is not known, and it is likely that longer lags are involved which differ between tree species. In addition, not all rain in showers reaches the forest understorey. In typical wet period in tropical rain forest c. 10-20% of rainfall is intercepted by the canopy (Muzylo et al. 2009), and in a seasonally dry or semi-deciduous forest interception can reach 35-40% (Junqueira et al. 2019, Lopes et al. 2020). Tani et al. (2003) recorded 17% interception at Pasoh, West Malaysia, in lowland rain forest for 1996-1999, but at the times of peak dryness of the 1997-98 ENSO event it rose to c. 40%. At Danum in the primary forest in wet periods interception was on average 17% (Douglas 2022), and by projection may have been similar at Danum in a dry period. These insights furthermore explain the quadratic relationship found in the present study between *gthi* and ln(rainfall), when only substantial showers had much effect on band readings. The wide scatter of points for days in these relationships is likely to have been a result of the precise time of day or night when rain fell too. A wetted stem early in the morning would partly dry during the rest of the day of a dry period, one wetted mid-afternoon would stay wet until the next day. Different sides of trees might also have been facing various canopy gaps for different parts of the day, which could have led to variability in any bark drying. Any stem flow after a shower on a relatively smooth and dry tree stem will probably simply run off without absorption.

### SMP and TEMP effects on large and small trees

The relationships from the temporal-autocorrelated adjusted time-series, were summarized in the slopes or coefficients of *gthi* on either SMP_−1_ and TEMP_0_. *Gthi* was a forward variable (i −> i+1), and SMP_−1_ and TEMP_0_ were backward ones (i−1 and i respectively). TEMP_0_ was an external variable to the trees, unaffected by their water fluxes (unless evapotranspiration was also cooling the forest), so it could be considered a cause of *gthi*, because it largely preceded *gthi* (there was approximately half of a day of overlap).

Although logger temperature was recorded inside the forest, it correlated well with external temperature at DVMC. SMP_-1_ was only in part an external variable though, because it was determined by both rain fall into the top 20 cm of soil (expressed as a total over the *n* preceding days), and by the uptake of water driven by evapotranspiration in the canopy (i.e. by TEMP_0_). SMP_-1_ was averaged across the four stations, that is it was a variable common to plot and forest (as was TEMP_0_). On hot sunny days (with high TEMP_0_) strong evapotranspiration pull will have acted on all trees, those being recorded *and* their neighbours. Thus, each tree’s response would have been in part influenced by trees in its own neighbourhood, although the data set does not allow an analysis at that level of SMP spatial resolution. Soil sensors in any case were not placed exactly close to particular banded trees, but distributed among them at random. SMP_-1_ was therefore a potential cause of *gthi,* but at the same time it was also determined by it when water flux was being driven by evapotranspiration.

It was SMP_-1_, and not SMP_0_, that was mostly the better predictor of *gthi*. TEMP_0_ could not have been a direct cause of SMP_−1_ since that would have implied causality acting in a backwards manner. It is possible though that TEMP_−1_, and even TEMP_−2_, being temporarily correlated with TEMP_0_, were acting on SMP_−1_, although their effects were not so strong as indicated by the two-term GLS model fits. It appears further that SMP_−1_ and TEMP_0_ were acting fairly independently and were therefore additive. SMP largely affected *gthi* the day before TEMP did. From this it must be concluded that had TEMP (i.e. evapotranspiration), and/or *gthi*, any indirect effect on SMP_−1_ this would have been prior (at a negative lag) to the direct effect of SMP_−1_ on *gthi*. A simple interpretation is that rainfall (positively) and evapotranspiration (negatively) together determined SMP_−1_.

In the wet period, large trees had notably the strongest negative SMP_−1_ coefficients, and these were similarly strong for SMP_0_, SMP_−1_, and SMP_−2_. The smaller trees were not only much more mixed in sign of the coefficients but also much more variable across the lags in SMP. Likewise, for TEMP the large trees had almost always positive coefficients in the wet period, especially for TEMP_0_ but more weakly for TEMP_−1_ and TEMP_−2_. These large trees were driving the evapotranspiration of the forest through water uptake, which was lowering SMP for all trees, large and small. It was unfortunate that four of the six large trees lacked dry-period SMP data (station 4). It is tentatively assumed that they would have responded like g12 and g22 to also have *gthi* reduced close to zero in the dry period, and as the predictions suggested. In the wet period, g12 behaved rather differently from the other five large trees, however. Large trees were affecting the SMP in the soil much more, also responding differently between periods, than small ones. Smaller understorey trees were mostly well shaded so their demands for water use were less than those of the canopy trees.

### Feedback and response

The aim of the present paper was not to advance mechanistic tree water relations models at the detailed level of (Zweifel et al. 2005, Steppe et al. 2006, 2008, Mencuccini et al. 2013) because complete series of climate variables were unavailable for Danum. Ideally radiation could have been used to estimate potential evapotranspiration, and RH recorded in the forest understorey. SMP was measured only in the top soil profile layer, and TEMP as ambient (logger) temperature. The data are then best used at the system level, in a phenomenological way, when asking whether different species had characteristically different responses that were possibly indicative of any water-use traits. Hidden within the daily mean changes in girth will have also been hysteresis (Garnier and Berger 1986, Simonneau et al. 1993), an asymmetric relationship between *gthi* and soil moisture potential, picked up as part of the diurnal patterns. Furthermore, to separate short-interval growth increments from water-flux-induced stem changes (Zweifel et al. 2001, Zweifel et al. 2005) would have been very difficult from visual inspection of the *gthi* time-series. There were rarely any clear ‘ledges’ over several days below which *gth* could be inferred to dip due to shrinkages, followed by an unambiguous ‘step-up’ in increment that could be attributable to intrinsic growth (Zweifel 2016). Each increment (*gthi*) was part swelling-shrinkage, due to changes in rates of water flux, and very small part growth, even when *gthi* was negative, in both wet or dry periods.

Most trees did have net positive changes over the entireties of the two periods which indicated net absolute growth.

An aspect of systems analysis is how feedback is expressed within a time-series: each day’s response as *gthi* will have depended on responses the day before, the system (tree) state, and the environmental conditions before and on the day considered. Evapotranspiration output, represented by TEMP, used the water supplied via the vascular network, and created a decrease in SMP which was balanced over time by an increase in potential as a result of rainfall input. Different species may have been more or less sensitive to decreases in SMP_−1_ (or what it represents) conditional on their different evapotranspiration demands. What was happening diurnally (Fig. 3) provided a set of averaged ‘reference pictures’ of the trees’ rhythms (Zweifel et al. 2006). The patterns became modified from day to day as SMP_−1_ and TEMP_0_ changed. A central question was whether different types of diurnal pattern related to the different forms of response described by the time-series regression coefficients.

How would different groups of understorey species have been expected ‘hypothetically’ to respond in a forest free of neighbours with varying influences (a common neighbourhood)? All species would have responded positively to increased TEMP_0_ (more radiation, more photosynthesis, more water usage and flux) but differently when water became limiting. For a small tree of the drought-tolerant understorey guild transiting from wet to dry period this might have led the positive dependence of *gthi* on SMP_-1_ to become less steep or close to zero. Being able to tolerate the lowest SMP_-1_ levels *gthi* was little affected, although possibly at the cost that there was no advantage when SMP_-1_ was high (under a form of ecophysiological trade-off operating). Conversely, a non-tolerant understorey species would probably have had a steeper coefficient than average, more affected when SMP_-1_ became very low yet not necessarily responding better when relieved of the stress. Drought-intolerant small trees of overstorey species would have been expected to reduce *gthi* at very low SMP_-1_ but responding appreciably better when SMP_-1_ was near zero. Additionally, there is perhaps a fourth guild, not obviously represented in the data set, composed of small trees of overstorey species that were more tolerant of drought (e.g. *Parashorea*). These trees would have been less affected by SMP_−1_ and responded to its increase at a slower rate than the intolerant species, again involving a possible trade-off.

An interesting feature of girth variation across the 18 trees and six species was that the coefficients of SMP_-1_ and TEMP_0_ as measures of response were better correlated with average diurnal maxima (*gthch_max_*) than minima (*gthch_min_*) of girth change. For instance, trees with small *gthch_max_* has positive SMP_-1_ coefficients, but those with large *gthch_max_* had more negative ones. Maxima indicate the proportional extents to which the stems were able to swell, and could be interpreted as the highest ‘elasticities’ of their stem tissues could achieve in conducting water. Minima, by contrast, showed the extents to which stems could shrink, and they appear to be less important than maxima for predicting tree responses to SMP and TEMP. Whilst detailed wood anatomy might well explain differences between species in this regard, it is pertinent to note that these maxima differed importantly between trees within any species, suggesting that they, the maxima, were contained by what their individual locations would physically allow. Large trees tended to have much smaller diurnal variations than small trees, and smaller *gthch_max_*-values.

Understanding how the internal cycle within the tree, expressed as response in *gthi*, was affected over time by the two external factors rainfall and TEMP, requires a recognition of reciprocal feedback control between TEMP (as proxy for evapotranspiration) and SMP. This was postulated to differ between wet and dry periods. It requires examining whether the time-series regression coefficients were positive or negative – and how significantly, and whether they accorded with the physics of tree-water relations for different forest structural guilds. When SMP declined its values became more negative: when TEMP declined it became less positive. Large overstorey trees are generally expected to exert much greater evapotranspiration pull on the soil water than smaller understorey ones due to their larger biomasses. It is important here to re-emphasize that SMP_-1_ and TEMP_0_ had coefficients in almost all cases of the same sign as, but stronger (and usually more significant) than, the alternative lags considered, viz. SMP_0_ and SMP_-2_, and TEMP_-1_ and TEMP_-2_. This is not to say that those latter variables were not acting at all. But, if these other variables were causing, even in part, *gthi* variation, it is inferred that they were doing so more weakly than the two most-significant ones highlighted. There were a few unexplained exceptions, however (Figs 4 and 5).

Grainger testing was not too informative because it could not test specific hypotheses about where the defining time interval between cause and effect was essential. On the other hand, it picked up significant causal effects among the lagged X-terms (SMP_0_, SMP_−1_ or SMP_−2_) and hence trees where SMP_−1_ was not the only best correlate. Granger-causation testing furthermore assumes unidirectional causality and does not guarantee that only ‘X caused Y’ — any more than GLS regression does, because both variables may have been driven by a third variable Z; X and Y were merely correlated (Eichler 2013). Cartwright (2007) and Kleinberg (2013) provide interesting discussions on the detection of causality from time-series data, and on the limitations of Granger’s approach. An important lack in the Danum stem girth data was not knowing all the other variables that influenced a focal tree’s SMP_−1_ and its *gthi*. Two such factors, highlighted above, were (1) how SMP changed with rooting depth, and (2) the influences of neighbour trees’ water uptake, and their <u>own</u> SMP_−1_-*gthi* relationships. Without having hierarchically nested models the method could not point to *which* lag in X was most important for predicting Y. Temporal autocorrelation in Y otherwise remained embedded in the Granger regression model. The GLS regression approach in this paper used just one measure of *gthi*, and whilst accounting for the temporal auto-correlation, it did relate the response variable to defined lags in SMP and TEMP. More onerous to compute than the fast omnibus Granger approach, piece-wise GLS regressions came closer to detecting and testing the real processes.

### Wet and dry periods

In the wet period, the basic idea is that rainfall adequately replaced the water removed by trees in upward transport supporting evapotranspiration. Rainfall, indirectly through soil water supply (and some canopy wetting), met the demands of photosynthesis and growth. The system longer-term must achieve stability. When SMP_-1_ increased, i.e. became less negative, and *gthi* also increased, the coefficient was positive. Water became more readily available to the trees, and this can be interpreted as having led to a relaxation of any water stress limitation on girth increment. But when SMP_-1_ increased and *gthi* decreased, the coefficient was negative, and seemingly girth increment was being reduced even though the stress was being apparently relaxed. It cannot have been that increasing *gthi* lowered SMP_-1_ because SMP_-1_ preceded *gthi*, but water uptake was lowering SMP in the days(s) before, and this not at that earlier time showing an increase on *gthi*, constituted a ‘delayed effect’. This explanation may go some way to solving the paradox.

Alternatively, for a small tree to have a negative *gthi*-SMP relationship indicated that increments at low SMP were larger than those at high SMP, i.e. the effect increasing SMP lowered *gthi*. This appears counter to the physics of water flow and stem expansion (Zweifel et al. 2001). It is highly unlikely that an understorey tree would lower SMP from high uptake (increased *gthi*) as would a large canopy one. Changing SMP with depth may be an explanation if the SMP that these small trees experienced was the inverse of what was recorded at 20 cm depth, i.e. where SMP was high at the surface it was low at lower depths and vice versa. The SMP at 20 cm depth acts as a reference level. This could be feasible if rain events were largely only wetting the top soil, or when dried out in the upper layers there was more water in the soil reachable by roots at lower depth. This would argue again for the critical dependence of use of water on rooting depth. Further, it would mean that *gthi*-SMP correlations, positive and negative were reflecting ‘normal’ and inverse SMP scales. To accommodate this hypothesis, it is necessary also to consider large and small trees sharing the upper 20-40 cm of soil and SMP being well correlated across this depth range, then below, e.g. 40-100 cm small trees differentially separate out their roots to avoid competition, and lastly below 1-2 m the deeper roots of the larger trees (Jenik 1978).

When *gthi* increased with increased TEMP_0_ though, the coefficient being positive, TEMP_0_ may have caused the *gthi* change as a result of strong demand and uptake of water, later leading to swelling of the stem but only when, presumably, there was a sufficient supply coming from the soil (i.e. SMP_-1_ was near zero). Otherwise, when *gthi* decreased with increase in TEMP_0_, the coefficient being negative, TEMP_0_ could also have caused this too, but this was a situation where water was being rapidly taken up and supply was <u>not</u> being sufficiently met (SMP_-1_ was likely already low), and this would have caused the stems to shrink under the demand (Sheil 2003).

When water availability was non-limiting, that is SMP was quite high and near zero, evapotranspiration would easily have proceeded, and when temperature was high, with little cloud cover and photosynthesis maximal, the girth would expand (Zimmermann 1983). In the dry period as SMP decreased evapotranspiration and water became more limiting in the soil, but also the air in the canopy was drier, then stomatal closure would presumably have limited water loss, though nevertheless what water was being used would have caused a positive *gthi* when evapotranspiration increased. It might then be expected that the effect of evapotranspiration bypassed *gthi* (i.e. *gthi* did not alter) and directly affected SMP, so that lower rainfall and water uptake together affected SMP, and this latter then affected *gthi*. The control of water use in the crowns might explain why there was practically no significant interaction between SMP and TEMP on *gthi* in the wet period. Predictions were that in the dry period the coefficient of *gthi* on TEMP was lower than that in the wet period, because less water was fluxing. Then, in addition to that, the lower SMP would have decreased *gthi*, and the feedback between lower leaf potential and stomatal control would have gradually stepped down the demand over several days (Jones 1992). In the short dry periods this presumably led to interactions which were variously transient. These lagged controls were subsumed within the time-series and very difficult to distinguish from effects of changing external factors.

In the dry period species’ responses would be expected to have differed as those species running short of water attempted to still access it, and as supplies fell due to lower rainfall (*rft*), increased TEMP_0_ would have resulted in lowered *gthi* (shrinkage). Lowered SMP_-1_ would have made the situation more acute. From wet to dry period the expectation was that coefficients for TEMP_0_ became more negative, and coefficients for SMP_−1_ became more positive, compared with the wet period. This was not always the case, however. Other factors are important to consider here. (i) Some trees may have been accessing water at greater depths in the dry compared with wet period, and this would not have been reflected in top-soil SMP changes, these latter largely governed by recently declining rainfall. (ii) Trees which were large, with higher open crowns, might have been more influenced by the dry period than smaller trees with lower shaded crowns in the understorey.

Although there were slight differences on average in slopes of *gthi* on SMP_−1_ and TEMP_0_ between ridge and lower-slope stations, these stations did not differ significantly in SMP, particularly in the dry period. Nevertheless, there was a tendency for the coefficients for SMP_−1_ to shift from positive on lower slopes to negative on ridges. In an earlier study in the same plots, Gibbons and Newbery (2003) obtained very similar results and likewise reached the same conclusions. Differences in *gthi* responses between stations were much more likely to have been due to how root distribution changed vertically and horizontally with respect to the SMP profile. Differences in leaf water potentials in wet and dry periods were explained for the drought-tolerant *Dimorphocalyx muricatus*, growing mainly on ridges, by the fact that it had relatively little surface rooting and more roots with depth, whilst the drought-intolerant *Mallotus wrayi*, growing across the whole topographic gradient, had conversely more roots near the surface and relatively few deeper ones (Gibbons and Newbery 2003). The SMP recorded at 20 cm depth in the present study was but an indicator of soil moisture status: SMP at lower depths were likely related to surface-layer SMP, albeit slightly differently for lower slopes and ridges depending on the maximum soil depths possible.

### Possible role of competition

The small trees were forced to respond positively to increased TEMP like the larger ones, but they would additionally, presumably, have suffered from competition for water with them as their neighbours. The different species’ time-series were in parallel, so the small tree increment changes are exactly concurrent with the those of the large trees. Hence what may appear matching in terms of the driving variables (TEMP_0_ and SMP_-1_) for the larger trees may not have been the case for the smaller understorey trees. Larger trees usually have both dense surface rooting, primarily for nutrient and water uptake, but also large deeper ones for access to additional water supplies in drier periods (e.g. Stahl et al. 2013). Smaller understorey trees with their less deep roots were therefore highly likely, even inevitably, competing with the larger overstorey ones and this would have lowered the SMP they experienced locally.

Competition for below-ground resources is generally considered to be symmetric, i.e. trees take up water in proportion to their biomasses. However, differences between storeys lie not only in tree size but that the overstorey ones are much more lighted than understorey ones, and this would lead competition for water to become asymmetric. The increase in the coefficients with relatively modest nearest-neighbour distance-weighted basal area values suggested that competition was operating. At night, though, water may have been laterally redistributed within the upper soil layers, which would ameliorate longer-term effects of competition, particularly in the wet period. Deeper water could also have been available to the shallower-rooted smaller trees by the process of hydraulic lift (Caldwell et al. 1991, Dawson 1993). Conversely, water may be redistributed by roots downwards from rewetted surface soil to drier layers below (Burgess et al. 1998, 2001). The relative strength of these equalizing processes would likely vary from time-to-time and place-to-place within the Danum plots depending on the precise neighbourhood of each tree. These processes would reinforce the competitive symmetry in water usage.

Above ground neighbour basal area may nevertheless not have been a good reflection of how root biomasses were distributed. Trees that then showed little difference between periods in their coefficients for SMP_−1_ and TEMP_0_ would appear tolerant to the dry period: in the dry period they will have had higher evapotranspiration than in the wet one, this causing – at least in the short term – attempts to increase water uptake. This may not necessarily have lowered SMP_−1_ though. An understorey tree which was not close to a large canopy one might have had a positive SMP_−1_ coefficient, whilst when the trees were close to one another, in competition for water, it had a negative coefficient because <u>locally</u> it was suffering water limitation, although the overall, large-tree dominated, <u>average</u> SMP_−1_ was indicating otherwise. In other words, where small and large trees would have had similar *gthi* at low SMP_−1_, when SMP_−1_ was closer to zero *gthi* for large trees was higher than at low SMP_−1_ (positive slope), whilst that for small trees *gthi* dropped lower (negative slope). SMP sensor values averaged across stations were effectively being re-weighted locally by the larger-tree effects. These factors may account for the almost consistently positive coefficients in wet and dry periods for all trees’ *gthi* with regard to TEMP but the mixed set of positive and negative coefficients (between and within species) in them with regard to SMP_−1_.

If increasing *gthi* of small trees should always, as a principle, be positively related to the increasing SMP where its roots mostly access water, then the problematic might be ‘stood on its head’, to show that those trees with decreasing relationships indicated rooting at deeper than the 20-cm reference. This might be a way of indirectly identifying rooting depths.

### Complex water flux dynamics

In the dry period stomatal closure would doubtless have played a role in slowing evapotranspiration and restricting water uptake as SMP decreased (Jones 1992). However, the relatively slow dynamics involved are spread over many days, as first the tree water stores were utilized and then the potentials began to decrease throughout the tree from crown to roots. With insufficient new rainfall the low SMP would not have been relieved, and the interplay with *gthi* would, if conditions remained constant for a while, equilibrate at a new state, albeit a likely unstable one. Different trees at different locations in the forest would have reduced the soil moisture content to different degrees locally, and to different depths, resulting in a highly heterogeneous 3D distribution. This might have equilibrated spatially, particularly at night for the top 20 (-40) cm of the profile, as mentioned. For this reason, the responses of different individuals in the dry period would be expected to have been even more variable than in the wet one, the more stressful conditions in the dry, exacerbating the relatively more stable situation in the wet period. The corresponding wet and dry period responses were clearly not well correlated, even though the data were coming from the same individuals over time.

Using the same trees across the whole study will have inevitably confounded differences *gthi* and SMP between the wet and dry periods with changes in neighbourhood tree dynamics in one or other of the periods, e.g. if a tree died nearby or lost large branches, parts of the canopy changed due to large-tree deaths even further away. In a related way, the short periods of volatility in *gthi* shown by four of the banded trees were most likely due to either the measured tree itself losing a branch or otherwise suffering damage and readjusting its water flux balance, or just temporarily being affected by local neighbour tree dynamics.

If the hydraulic architecture of trees maps to a simple linear system, over time input ‘on average’ will balance output, with lags (Chatfield 2004). The swelling and shrinking of the stem is in a feedback loop with the increase and decrease in SMP in the soil. TEMP, as a proxy for photosynthesis and evapotranspiration pull, externally controls water uptake, and this water is replaced by inputs from rainfall. Rainfall therefore drove *gthi* change via SMP: “*rft* −> SMP <−> *gthi* <− TEMP”. SMP and *gthi* were reciprocally coupled, and driven and stabilized by these two main external variables, *rft* and TEMP. If *gthi* is seen as an inverse measure of tree moisture potential (TMP), it entails an internal feedback loop between SMP and *gthi*. The two variables may be displaying a Lorenz manifold (Sugihara et al. 2012, Ye et al. 2015) in a way that mirage correlations are more or less strong on one or the other fold depending on whether *rft* or TEMP are having the relatively stronger external influence. The mix of positive and negative coefficients found within and between understorey species when fitting *gthi* to SMP (but not for TEMP) in the GLS regressions — positive and negative correlations essentially, may have been a result of different individuals being ‘caught’ on different parts of a general attractor due to their particular set-up system circumstances in the time period studied. Some series showed a positive, others negative, correlation. This would appear a candidate set-up for convergent cross mapping. The shape of such a manifold might be different in wet and dry periods.

### Conclusion: confronting hypotheses

It was previously proposed that the understorey trees might be divided into (a) the small juveniles of overstorey species, and (b) all trees of true understorey species, subdivided into (b1) drought-tolerant and (b2) drought-intolerant groups (Newbery et al. 1999). The time- series analyses here did not provide clear confirmation of traits related to water-flux related stem changes that would distinguish species in the proposed structural-dynamic guilds of understorey tree species at Danum (Newbery et al. 1999, Newbery et al. 2011). Individual trees within species behaved quite idiosyncratically. This is interpreted to mean that the relationship between *gthi* and changing SMP (driven by temperature on evapotranspiration and by rainfall input) depended on the exact constellation of each and every forest location and its conditions, particularly in this case the water fluxes of neighbouring trees. It could also mean that the proposed manifold dynamics were interacting with neighbour competitive effects.

There was moreover no case to be made for the predicted species in the drought- intolerant guild (*M. wrayi*) consistently showing water stress whilst those in the drought- tolerant one (*D. muricatus, L. beccarianum*) indicating different means of coping with the shortages, even though these species show markedly differing spatial distributions with respect to plot topography (Newbery et al. 1996). Possibly more indicative responses would have been recorded under much stronger drought conditions (Newbery and Lingenfelder 2004), ones that matched the stronger events in the past when selection may have operated also more strongly (Newbery et al. 2011). The varying responses by different individuals of the understorey species have been also due to high plasticity in form and response within species, determined by local tree neighbourhood in terms of the complexities of tree rooting and soil moisture three dimensional distributions. Downward water redistribution would favour the deeper-rooted *D. muricatus*, upward the shallower-rooted *M. wrayi*.

Determining species traits by reduction from the structure and dynamics of their structural guilds, and detecting these traits in the field from long time series of measurements of individuals did prove both highly challenging and inconclusive. Investigations are stymied by the severe lack of detailed knowledge of the spatial extents of the constituent species’ individual tree rooting systems and their physiologies. The technicalities involved are still quite primitive, and the need for replication of matching SMP recordings below ground is formidable. Knowledge of tropical forest root systems remains extremely limited. In recent decades, little attempt was made to advance beyond the framework laid by Jenik (1978). One study to estimate approximate rooting depths and lateral root distributions of trees in dipterocarp forest was made by Baillie and Mamit (1983) on road-side cuttings.

The highly interrelated processes running in a time series of even such a relatively small model system with one feedback loop and two external drivers, had primary effects that were likely variably and asynchronously lagged; and there were secondary effects applying. This led to very complicated dynamics at the individual tree level. The changing neighbourhood variables added another important source of spatial and temporal variability (noise) at the population-community level, a form of stochasticity. The forest overall though was functioning in physical and physiological terms at the whole ecosystem level with relatively stable dynamics, subsuming and equilibrating all the minor instabilities and variations of the two lower (population/individual) levels. The physics of water redistribution by equilibration along gradients, diurnally and over several days with lags, may contribute to some amelioration of competition for water below ground.

On the basis of the findings in this work the hypothesis that apparent drought-tolerant tree species in the understorey have distinctly different responses to SMP from non-drought-tolerant ones, and to small trees of overstorey species, must be rejected. Apart from using larger random samples of trees of each species and recording over several dry events, the key complicating factor is local variability in root competition resulting from neighbourhood stochasticity. This makes tree growth responses highly individualistic and difficult to predict.

## Acknowledgements

This work was part of the Royal Society of London’s SE Asian Rain Forest Research Programme 1985-2015. We thank the Sabah Biodiversity Council and the Danum Valley Management Committee for permission to continue research at Danum. Dr R. C. Ong (Sabah Forest Department) kindly provided host support. We appreciate the assistance of Jamil Hanapi in monitoring the stations and handling data downloads and transfer on several occasions, and of Saidih Samat and Anthony Karolus with the installation and maintenance of field equipment. Reviewers are thanked for their comments.

## Author contributions

The research was conceived jointly by DMN and ML. ML undertook bulk of the field work and prepared the raw data files. DMN statistically analysed the data set, and very largely wrote the paper with inputs from ML. Both authors read, discussed and corrected the various versions.

## Data Accessibility

Newbery and Lingenfelder (2020).

## List of appendices as electronic supplementary material

1. Rainfall, soil moisture potential and temperature relationships; tree variables, stations and topography, model sensitivity analysis, causality test results; predictive response equations.

2. Tests of dependence of girth increment on relative humidity, conditioned on temperature.

3. Assessment of the effect of temperature on the dendrometer band mechanism.

4. Analysis of effects of showers in the dry period on girth increment measurement.

5. Tables of GLS-arima model fits, one and two-term, for wet and dry periods.

3. Bar graphs of SMP and TEMP coefficients from two-term GLS-arima regressions.

