## Appendices 1-6 for "Stem girth changes in response to soil water potential in lowland dipterocarp forest in Borneo: a phenomenological and individualistic time-series analysis"

**Appendix 1: Table S1**. Details of the 18 selected trees and their electronic bands (eDDS), with species codes, reference tags, three measures of canopy exposure (layer, space and light level – defined in the footnotes), these last averaged into a canopy status (canStat) value, and relative growth rates from main plot census, 2001-2007 (*rgr*) for the wet and dry periods.

| | | | Canopy | | | | Rgr in $P_3$ |
|---|---|---|---|---|---|---|---|
| Band | Tag | Code | Layer[1] | Space[2] | Light[3] | CanStat | (mm/m/yr) |
| g11 | 560 | Mw | 2 | 2 | 2 | 2.00 | 14.01 |
| g12 | 567 | Sf | 4 | 4.5 | 5 | 4.50 | 49.70 |
| g14 | 577 | Lb | 2.5 | 3 | 3 | 2.83 | 20.62 |
| g15 | 1084 | Sf | 2 | 3 | 1.5 | 2.17 | 10.84 |
| g22 | 1662 | Pm | 3.5 | 3.5 | 3.5 | 3.50 | 8.96 |
| g23 | 1732 | Dm | 2 | 3 | 2 | 2.33 | 9.36 |
| g24 | 1685 | Mw | 2 | 3 | 3 | 2.67 | 10.69 |
| g25 | 2405 | Sf | 2 | 2 | 2.5 | 2.17 | 23.85 |
| g31 | 7680 | Mw | 2.5 | 2 | 2 | 2.17 | 9.33 |
| g32 | 6892 | Sp | 4 | 4 | 4.5 | 4.17 | 55.42 |
| g33 | 6900 | Lb | 3 | 2.5 | 2.5 | 2.67 | 33.91 |
| g34 | 7436 | Dm | 2 | 2.5 | 3 | 2.50 | 2.11 |
| g35 | 7586 | Dm | 2 | 3 | 1.5 | 2.17 | 0.00 |
| g41 | 8077 | Sf | 2 | 2.5 | 2 | 2.17 | 0.00 |
| g42 | 8060 | Dm | 2.5 | 2 | 1.5 | 2.00 | 0.00 |
| g43 | 8321 | Pm | 4 | 4 | 4 | 4.00 | 12.30 |
| g44 | 8848 | Sp | 4 | 4 | 4.5 | 4.17 | 18.94 |
| g45 | 8810 | Pm | 3 | 4 | 3.5 | 3.50 | 8.92 |

[1] Layer of tree: 1 tree-let; 2 understorey; 3 mid-level; 4 top canopy; 5 emergent. [2] Space of crown: 1, completely suppressed; 2, mixed/entangled from all sides; 3, significantly overlapping most sides; 4, stems touching from some sides or above; 5, crown free, unshaded; by neighbours. [3] Light level: 1 < 30%; 2, 30 - < 50%; 3, 50 - < 70%; 4, 70 - < 90; 5, ≥ 90% of crown receiving full light

**Appendix 1: Table S2**. Means, with standard deviations and lower and upper quartiles, of the 1-day girth increment, *gthi*, for the wet (n = 550) and dry (n = 93-98; see text) periods. Values are *gthi* in cm x 1000; sd is standard deviation; q1 and q3 are lower and upper quartiles. Codex, are the codes in Table 2, with individuals within species numbered.

| nr | station | tree | Codex | band | wet | | | | dry | | | |
|---|---|---|---|---|---|---|---|---|---|---|---|---|
| | | | | | mean | sd | q1 | q3 | mean | sd | q1 | q3 |
| 1 | 1 | 1 | Mw1 | g11 | 0.589 | 9.681 | −4.0 | 6.5 | 1.623 | 8.564 | −2.9 | 5.5 |
| 2 | 1 | 2 | Sf1 | g12 | 8.320 | 30.850 | −9.6 | 23.2 | 1.128 | 4.692 | −2.1 | 4.6 |
| 3 | 1 | 4 | Lb1 | g14 | 0.540 | 23.500 | −4.9 | 6.0 | 0.770 | 13.110 | −6.4 | 2.8 |
| 4 | 1 | 5 | Sf2 | g15 | −0.451 | 14.875 | −9.4 | 9.5 | −1.480 | 14.950 | −12.0 | 8.3 |
| 5 | 2 | 2 | Pm1 | g22 | 0.840 | 23.860 | −5.0 | 5.4 | −0.628 | 8.915 | −6.3 | 3.1 |
| 6 | 2 | 3 | Dm1 | g23 | 0.097 | 9.347 | −4.0 | 4.6 | 0.210 | 6.591 | −4.6 | 5.0 |
| 7 | 2 | 4 | Mw2 | g24 | −0.170 | 4.264 | −2.7 | 1.9 | 1.369 | 8.674 | −3.3 | 4.4 |
| 8 | 2 | 5 | Sf3 | g25 | −0.240 | 10.505 | −6.0 | 6.5 | −0.560 | 11.560 | −7.1 | 5.2 |
| 9 | 3 | 1 | Mw3 | g31 | −0.057 | 4.655 | −2.7 | 2.3 | 2.450 | 18.180 | −5.6 | 6.9 |
| 10 | 3 | 2 | Sp1 | g32 | 7.397 | 10.933 | 0.6 | 14.6 | 5.996 | 5.298 | 3.0 | 10.2 |
| 11 | 3 | 3 | Lb2 | g33 | 2.470 | 4.047 | −0.2 | 4.6 | 3.571 | 7.780 | −1.3 | 7.4 |

| | | | | | | | | | | | | |
|---|---|---|---|---|---|---|---|---|---|---|---|---|
| 12 | 3 | 4 | Dm2 | g34 | −0.240 | 4.919 | −3.3 | 2.7 | 0.929 | 5.310 | −3.0 | 5.2 |
| 13 | 3 | 5 | Dm3 | g35 | 0.035 | 13.139 | −5.6 | 6.6 | 0.277 | 7.099 | −4.0 | 5.6 |
| 14 | 4 | 1 | Sf4 | g41 | −0.090 | 3.493 | −1.9 | 1.7 | – | – | – | – |
| 15 | 4 | 2 | Dm4 | g42 | 0.001 | 4.024 | −2.3 | 2.3 | – | – | – | – |
| 16 | 4 | 3 | Pm2 | g43 | 3.831 | 22.445 | −8.1 | 13.1 | – | – | – | – |
| 17 | 4 | 4 | Sp2 | g44 | 3.641 | 9.459 | −1.9 | 8.7 | – | – | – | – |
| 18 | 4 | 5 | Pm3 | g45 | −0.933 | 10.674 | −6.3 | 3.7 | – | – | – | – |

Appendix 1: Table S3. Mean and SEs of the soil moisture potential (SMP with lag 1 d) and logger temperature (no lag) coefficients from the finally selected GLS time-series regressions, that are plotted in Figs. 4 and 5. Here 'code' in Table 3 is expanded to 'codex' to show the tree number per species as labelled on the Figs 6-10. Note: the coefficients are not here standardized for girth, as they are in Figs 6-10.

| number | station | tree | codex | gth | $SMP_{-1}$ wet | | $TEMP_0$ wet | | $SMP_{-1}$ dry | | $TEMP_0$ dry | |
|---|---|---|---|---|---|---|---|---|---|---|---|---|
| | | | | | est | se | est | se | est | se | est | se |
| 1 | 1 | 1 | Mw1 | g11 | 1.517 | 0.218 | 0.945 | 0.310 | -0.103 | 0.125 | 0.230 | 0.439 |
| 2 | 1 | 2 | Sf1 | g12 | −4.688 | 0.923 | −5.249 | 0.915 | -0.026 | 0.096 | -1.176 | 0.267 |
| 3 | 1 | 4 | Lb1 | g14 | −0.046 | 0.431 | −0.736 | 0.504 | -0.617 | 0.245 | 2.882 | 0.674 |
| 4 | 1 | 5 | Sf2 | g15 | 2.297 | 0.345 | 2.182 | 0.457 | 1.298 | 0.247 | 0.900 | 0.706 |
| 5 | 2 | 2 | Pm1 | g22 | −1.531 | 0.221 | 2.219 | 0.365 | -0.155 | 0.141 | 2.805 | 0.424 |
| 6 | 2 | 3 | Dm1 | g23 | 0.934 | 0.249 | 1.587 | 0.294 | 0.514 | 0.155 | -1.028 | 0.390 |
| 7 | 2 | 4 | Mw2 | g24 | −0.166 | 0.078 | 0.574 | 0.130 | -0.425 | 0.111 | 1.803 | 0.402 |
| 8 | 2 | 5 | Sf3 | g25 | 1.517 | 0.245 | 1.997 | 0.316 | 0.893 | 0.126 | 1.460 | 0.629 |
| 9 | 3 | 1 | Mw3 | g31 | −0.235 | 0.056 | 0.584 | 0.120 | -0.897 | 0.218 | 3.567 | 0.880 |
| 10 | 3 | 2 | Sp1 | g32 | −0.094 | 0.320 | 0.504 | 0.339 | 0.549 | 0.091 | 0.274 | 0.309 |
| 11 | 3 | 3 | Lb2 | g33 | −0.455 | 0.104 | 0.862 | 0.136 | -0.197 | 0.174 | -1.905 | 0.466 |
| 12 | 3 | 4 | Dm2 | g34 | −0.430 | 0.100 | 0.141 | 0.158 | -0.362 | 0.113 | -0.134 | 0.247 |
| 13 | 3 | 5 | Dm3 | g35 | 1.248 | 0.291 | 2.553 | 0.419 | 0.287 | 0.095 | 0.230 | 0.439 |
| 14 | 4 | 1 | Sf4 | g41 | −0.271 | 0.094 | 0.619 | 0.181 | – | – | – | – |
| 15 | 4 | 2 | Dm4 | g42 | 0.504 | 0.073 | 1.055 | 0.138 | – | – | – | – |
| 16 | 4 | 3 | Pm2 | g43 | −2.825 | 0.480 | 2.111 | 0.693 | – | – | – | – |
| 17 | 4 | 4 | Sp2 | g44 | −2.291 | 0.277 | 0.567 | 0.302 | – | – | – | – |
| 18 | 4 | 5 | Pm3 | g45 | −1.205 | 0.221 | 0.480 | 0.307 | – | – | – | – |

Appendix 1: Table S4. *F*-ratio statistics from the Granger causality test for the *gthi* time series of the 18 bands, in the wet and dry periods. 'gth' is the band identifier, 'Codex' for species is as listed in Appendix 1: Table S2. The two independent variables were soil moisture potential (SMP) and logger temperature (TEMP), as used before in the GLS-arima models.

| Number | Station | Tree | Codex | gth | Wet period | | Dry period | |
|---|---|---|---|---|---|---|---|---|
| | | | | | $SMP_0$ | $TEMP_0$ | $SMP_0$ | $TEMP_0$ |
| 1 | 1 | 1 | Mw1 | g11 | 30.06*** | 13.33*** | 3.86* | $0.94^{ns}$ |
| 2 | 1 | 2 | Sf1 | g12 | 9.04*** | $2.49^{o}$ | $2.17^{ns}$ | $1.32^{ns}$ |
| 3 | 1 | 4 | Lb1 | g14 | 3.66* | $2.20^{ns}$ | 6.13** | 3.95* |
| 4 | 1 | 5 | Sf2 | g15 | 25.76*** | 6.67*** | 14.77** | 14.25*** |
| 5 | 2 | 2 | Pm1 | g22 | 7.94*** | $2.03^{ns}$ | 5.45** | 3.90* |
| 6 | 2 | 3 | Dm1 | g23 | 8.03*** | 5.61** | 8.37*** | 7.91** |
| 7 | 2 | 4 | Mw2 | g24 | 5.10** | $1.49^{ns}$ | 6.86** | 4.05* |
| 8 | 2 | 5 | Sf3 | g25 | 25.43*** | 11.16*** | 5.41** | 3.70* |
| 9 | 3 | 1 | Mw3 | g31 | 15.85*** | 12.05*** | 7.47** | 4.58* |
| 10 | 3 | 2 | Sp1 | g32 | $0.10^{ns}$ | $1.06^{ns}$ | 16.38*** | 10.54*** |
| 11 | 3 | 3 | Lb2 | g33 | 8.30*** | $0.18^{ns}$ | 4.79* | 4.11* |
| 12 | 3 | 4 | Dm2 | g34 | 9.93*** | $1.98^{ns}$ | 8.13*** | $1.76^{ns}$ |
| 13 | 3 | 5 | Dm3 | g35 | 10.40*** | $0.50^{ns}$ | 4.00* | 5.80** |
| 14 | 4 | 1 | Sf4 | g41 | $2.52^{o}$ | $1.95^{ns}$ | -- | -- |
| 15 | 4 | 2 | Dm4 | g42 | 24.42*** | 13.43*** | -- | -- |
| 16 | 4 | 3 | Pm2 | g43 | 15.35*** | $2.37^{o}$ | -- | -- |
| 17 | 4 | 4 | Sp2 | g44 | 27.89*** | $2.77^{o}$ | -- | -- |
| 18 | 4 | 5 | Pm3 | g45 | 15.12*** | $2.25^{ns}$ | -- | -- |

***, $P \leq 0.001$; **, $P \leq 0.01$; *, $P \leq 0.05$; $^{o}$, $P \leq 0.10$, $^{ns}$, $P > 0.10$.

Appendix 1: Table S5. Linear regression statistics of SMP and TEMP estimates from GLS-arima regressions on relative maximum diurnal change in stem girth (*gthch*$_{max}$) of the trees with bands ($\beta_0$, intercept; $\beta_1$slope) in the dry period (g15 excluded, *n* = 12), for (a) single-term models with the three lags of no and one or two days, and (b) two-term models with lags of no and one day crossed. These regressions were applied to predict the missing estimates for station 4 in the dry period.

(a) Single-term models

| variate | lag | $\beta_0$ | $\beta_1$ | *F*-value[a] | P(*F*) | $R^2$(%)[b] |
|---|---|---|---|---|---|---|
| SMP | 0 | 0.593 | –0.302 | 27.2 | < 0.001 | 62.6 |
| | 1 | 0.604 | –0.279 | 23.0 | < 0.001 | 58.2 |
| | 2 | 0.546 | –0.232 | 24.8 | < 0.001 | 58.8 |
| TEMP | 0 | –0.523 | 0.686 | 9.1 | 0.013 | 25.2 |
| | 1 | –1.128 | 0.490 | 9.0 | 0.013 | 21.9 |
| | 2 | –1.420 | 0.876 | 35.1 | < 0.001 | 67.5 |

[a] df = 1, 10, [b] adjusted.

(b) Two-term models

| variate | SMP-lag | TEMP-lag | $\beta_0$ | $\beta_1$ | *F*-value[a] | P(*F*) | $R^2$(%)[b] |
|---|---|---|---|---|---|---|---|
| SMP | 0 | 0 | 0.582 | –0.259 | 17.73 | 0.002 | 60.3 |
| | 0 | 1 | 0.498 | –0.284 | 37.17 | < 0.001 | 76.7 |
| | 1 | 0 | 0.602 | –0.241 | 16.19 | 0.002 | 58.0 |
| | 1 | 1 | 0.530 | –0.252 | 27.13 | < 0.001 | 70.4 |
| TEMP | 0 | 0 | –0.095 | 0.411 | 4.68 | 0.056 | 25.1 |
| | 0 | 1 | –0.604 | 0.136 | 0.86 | 0.375 | –1.3 |
| | 1 | 0 | –0.316 | 0.511 | 7.20 | 0.023 | 36.1 |
| | 1 | 1 | –0.629 | 0.261 | 4.99 | 0.050 | 26.6 |

[a] df = 1, 10, [b] adjusted.

Appendix 1: Fig. S1.  Relationship between soil moisture potential (SMP, sqrt-transformed) and the 20-day running rainfall total in the wet and dry periods. The fitted curves were: wet, Y = −5.61 − 8.80·(0.9824$^X$) + 0.00439·X ($R^2$= 34.3%, n = 550); dry, Y = −1.30 − 30.9·(0.9837$^X$) − 0.0270·X ($R^2$= 68.4%, n = 98).

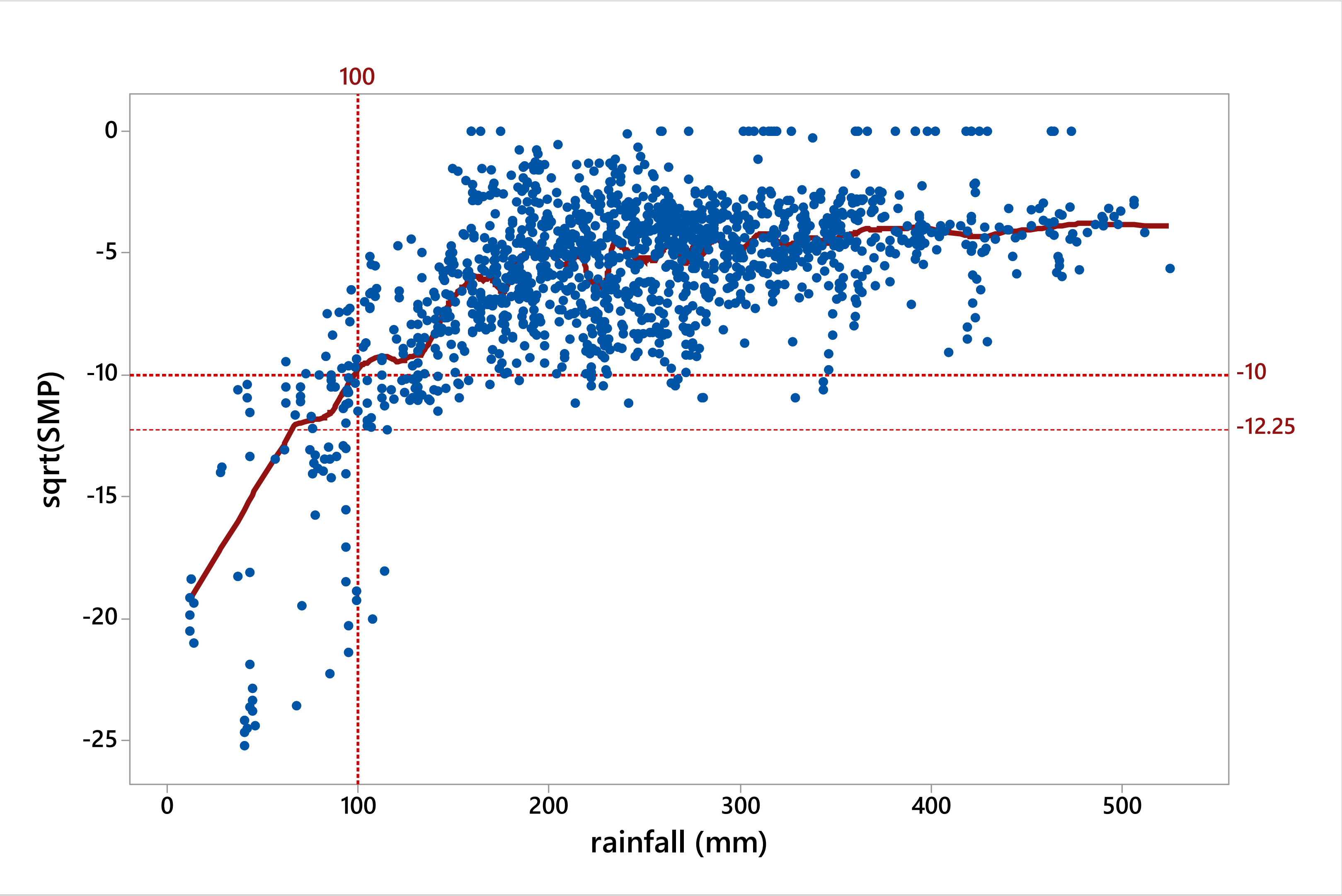

Appendix 1: Fig. S2. Non-linear regression model fits of the relationship between square-root transformed soil moisture potential (SMP_sqrt) and the 20-day rainfall totals (rft), for the wet and dry periods at Danum.

WET: SMP_sqrt = −5.611 − 8.80*(0.9824^rft) + 0.00439*rft; $F$ = 96.7, df = 3, 546, $P$ < 0.001.

DRY: SMP_sqrt = −1.30 – 30.9*(0.9837^rft) – 0.0270*rft; $F$ = 71.0, df = 3, 94, $P$ < 0.001.

Fig. S2a

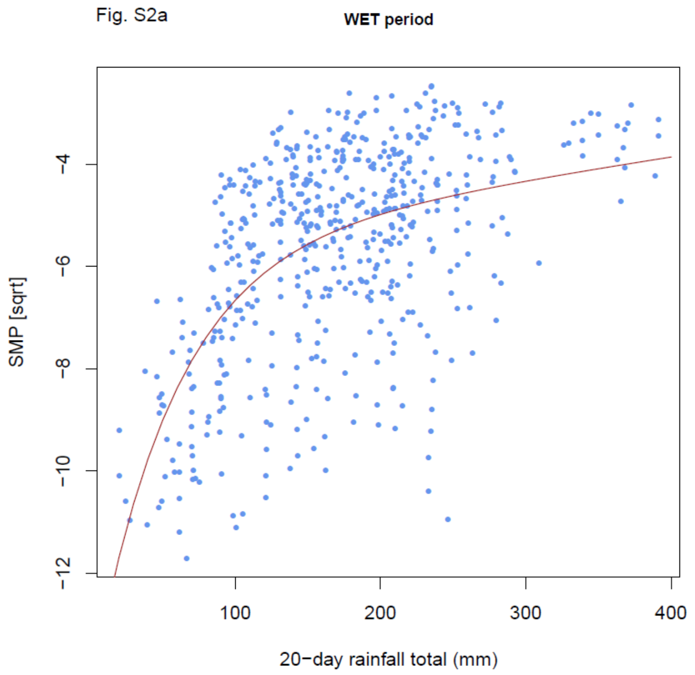


Fig. S2b

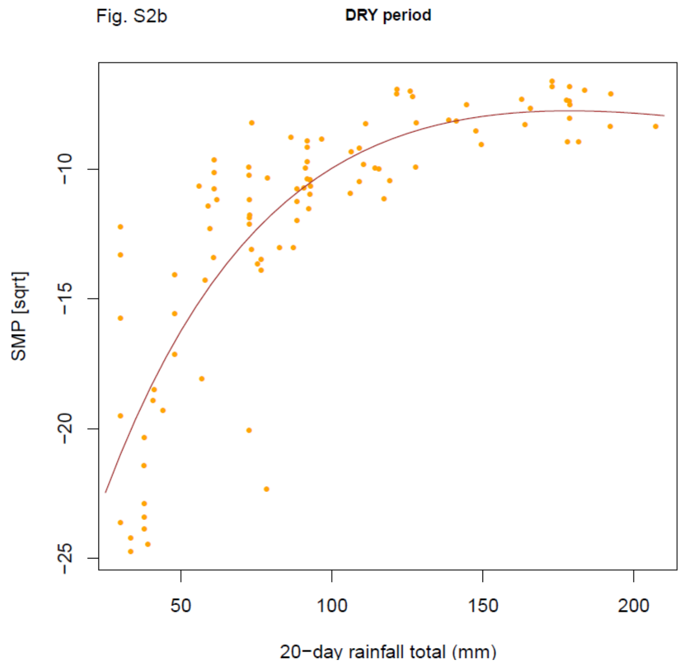

Appendix 1: Fig.S3. Mean diurnal courses of (a) soil moisture potential (SMP), across all stations; and (b) logger temperature (TEMP), for station 1 as example, in the wet and dry periods. For SMP, means are for the 30 successive wet sub-periods (LL and UL – the 95% confidence limits), but for the three dry sub-periods separately.

(a) SMP

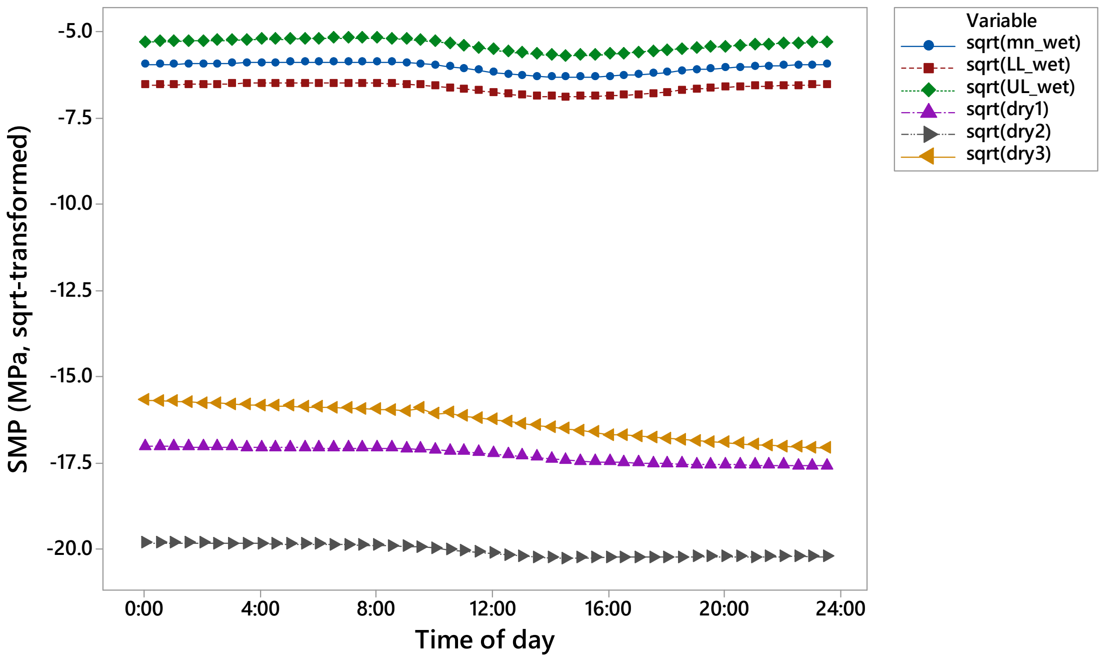


(b) TEMP

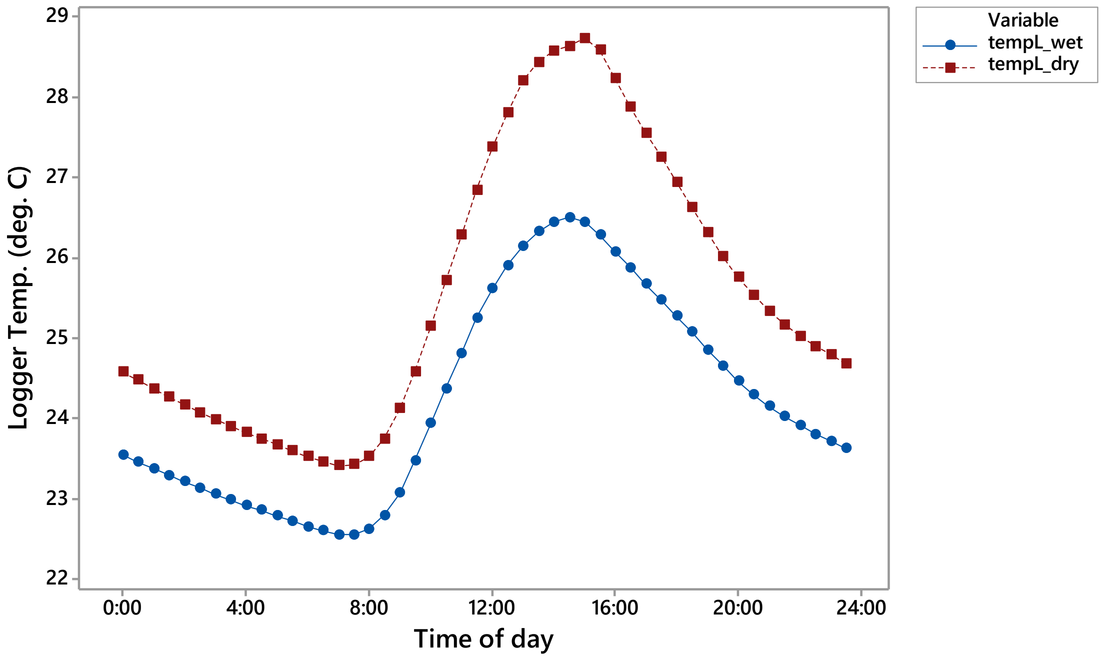

Appendix 1: Fig. S4. Comparison of the $SMP_{-1}$ estimates from GLS regression models with or without 'rain' (20-d-rft, sqrt-transformed) as an addition term, for (a) the wet (excluding the four bands where GARCH models were required (see main text), and (b) the dry, periods. Excluding the strong and unusually behaving outlier (g12) in the wet period, and the two bands with atypical 1-day lagged rainfall dependencies in the dry one, corresponding non-linear curve fits are shown in (c) and (d).

(a) Wet

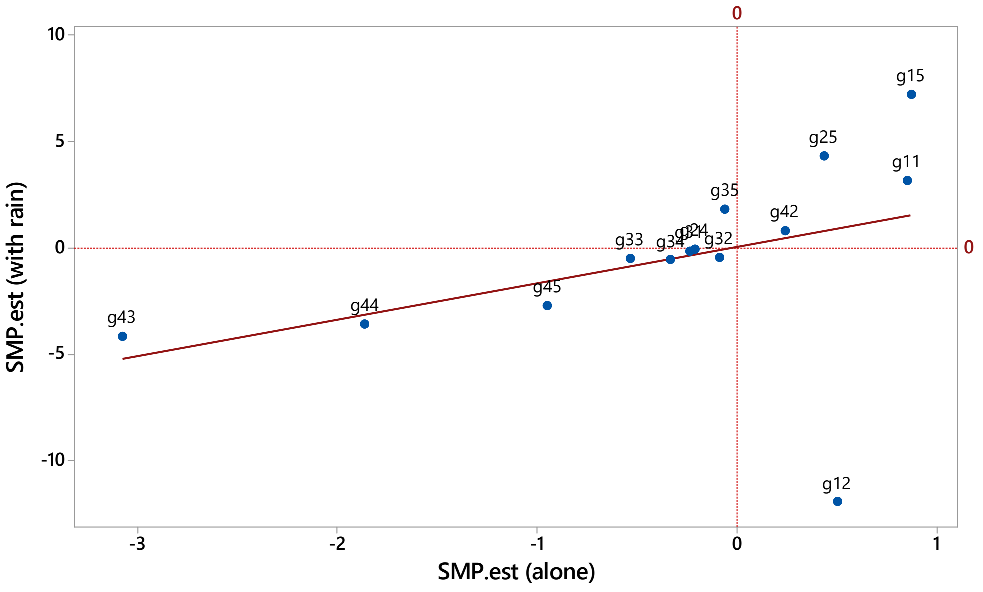


(b) Dry

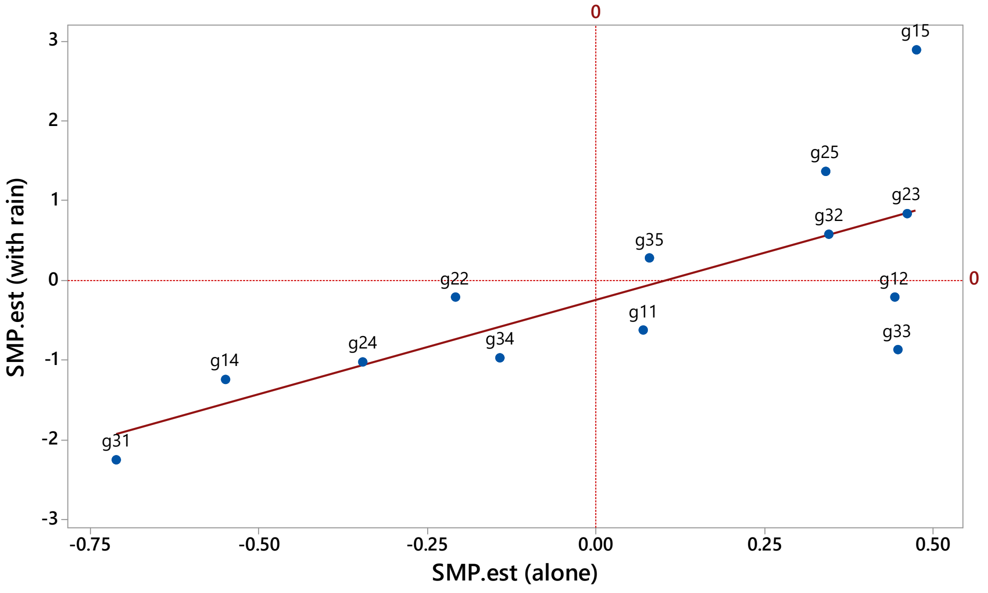

(c) Wet - fitted curve
[Y = 0.9429 + 4.286 X + 0.8654 X^2;  R^2adj = 85.9%]

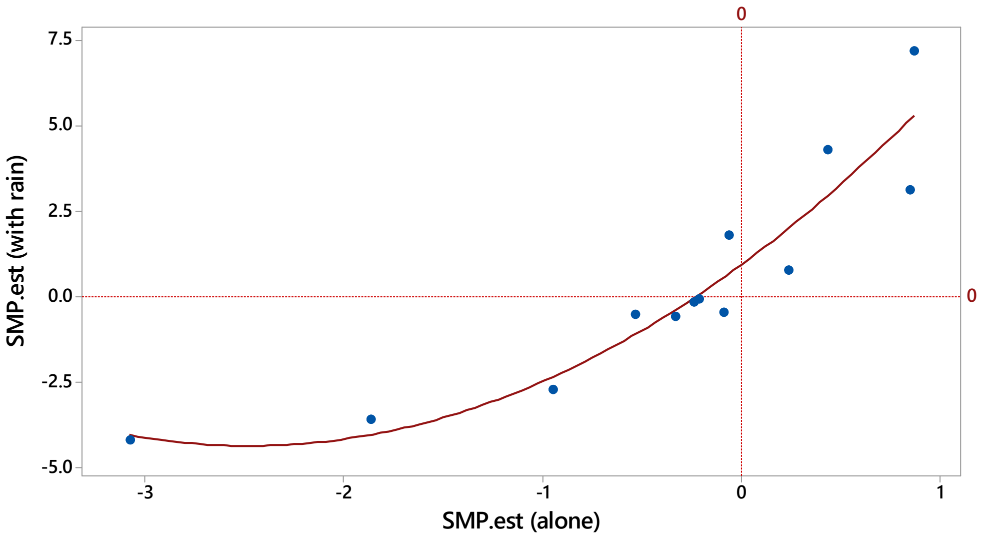


(d) Dry - fitted curve
[Y  = - 0.2025 + 3.352 X + 1.442 X^2;  R^2adj = 77.1%]

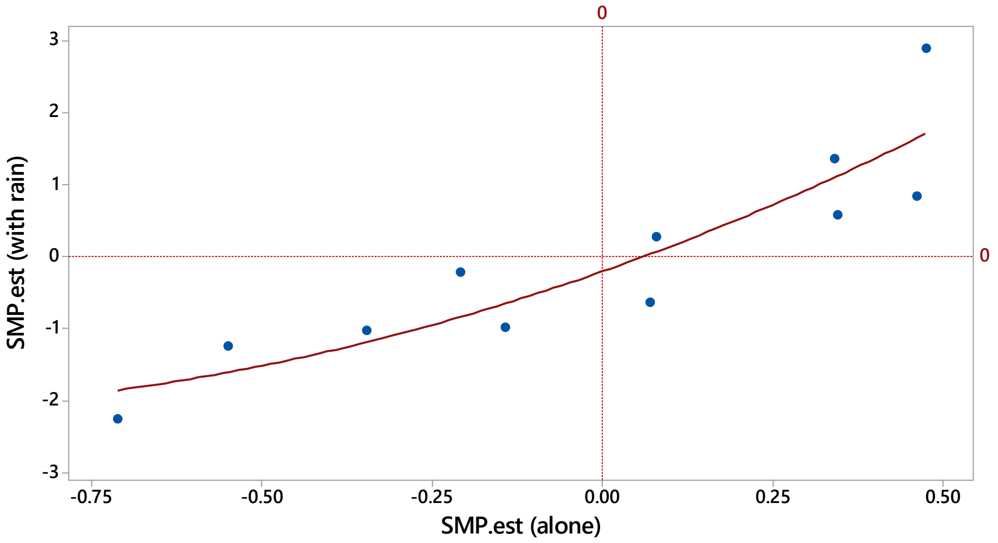

Appendix 1: Fig. S5. Standardized SMP−1 versus TEMP0 coefficients graphed together for (a) wet and (b) dry periods for the 18 respective 13 trees analyzed for their gthi time-series, and their corresponding dry- and wet-period (c) SMP−1 and (d) TEMP0 values against one another, annotated with the station numbers. Large trees (dcl = 1), filled circles; small trees (dcl = 2), filled triangles. This is equivalent to Fig. 7 in the main text where species labels are shown. Station locations are shown in Fig. 1 in the main text.

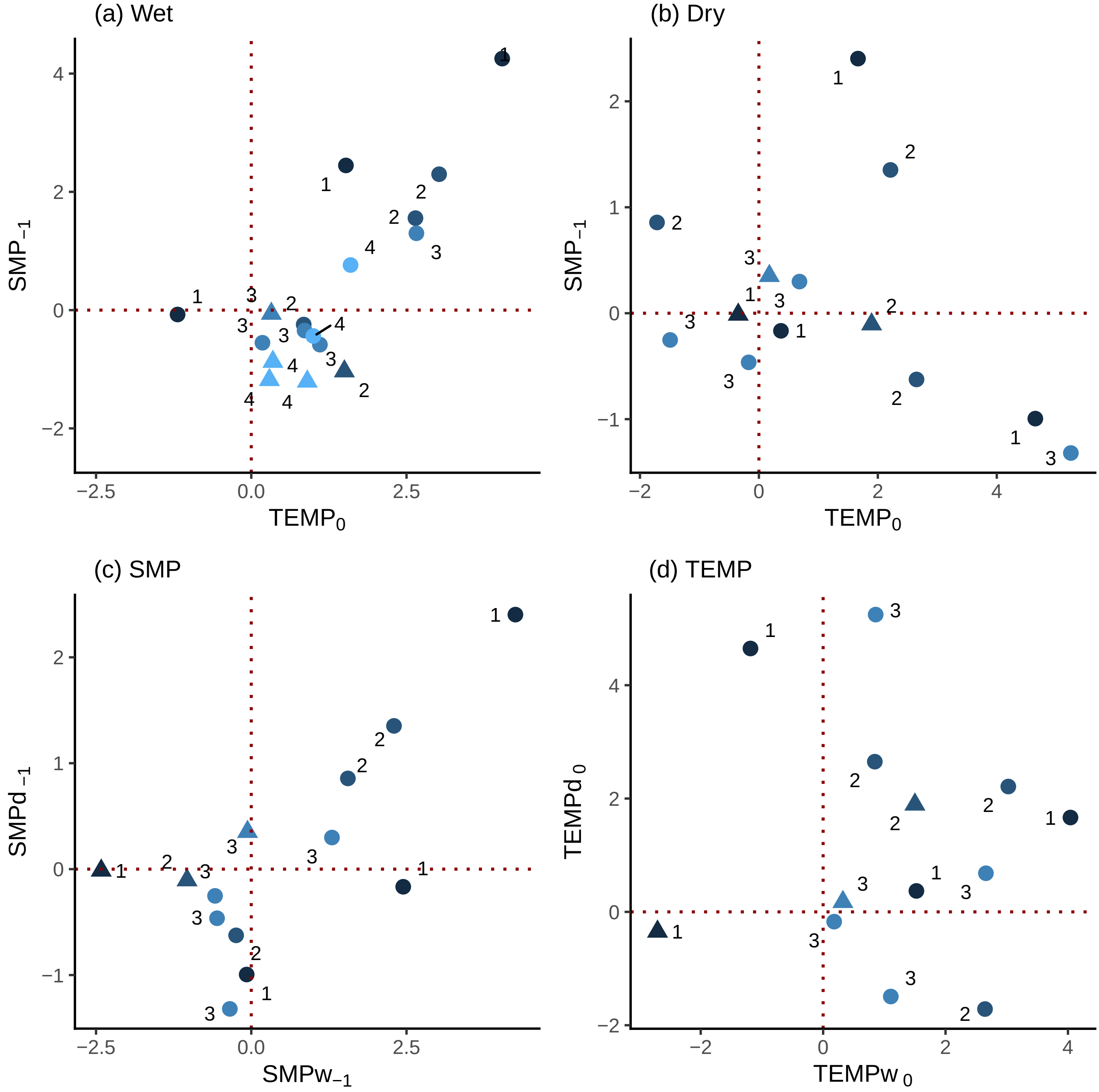

Appendix 1: Fig. S6. Dependence of the SMP–1 and TEMP0 estimates from the single-term GLS-arima regression models on the inverse-distance weighted basal area abundance (BA/d) of neighbouring trees to the banded one with a 5 m radius, for the available small trees (scl = 1), in the wet and dry periods. Species codes are as shown in Table 2 of the main text (or in Appendix 1: Table S1).

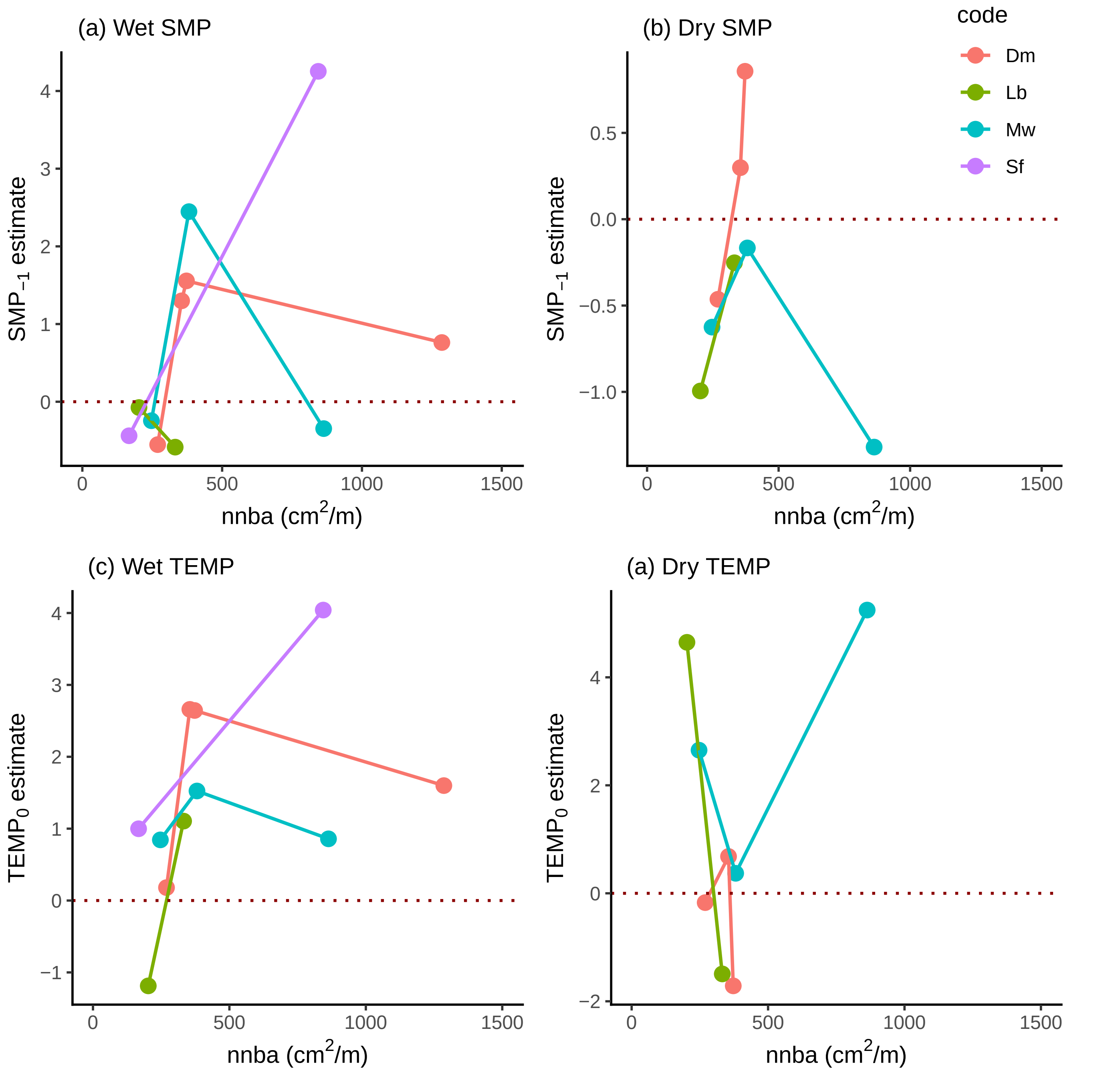

Appendix 2: Table S1. Statistics for the days within the studied wet and dry periods for which records of dry-bulb temperature and relative humidity at 14::00 h (Tdry14, RH14) were available at the Danum Valley Field Centre.

| | n days | mean ± SD | median | minimum | maximum |
|---|---|---|---|---|---|
| Wet | | | | | |
| Tdry14 ($^oC$) | 357 | 30.0 ± 2.1 | 30.4 | 24.6 | 34.6 |
| RH14 (%) | 352 | 80.2 ± 8.4 | 79.7 | 61.3 | 98.3 |
| Dry | | | | | |
| Tdry14 ($^oC$) | 67 | 31.4 ± 2.5 | 32.2 | 25.1 | 34.9 |
| RH14 (%) | 69 | 76.7 ± 10.8 | 75.0 | 59.5 | 100.0 |

Appendix 2: Table S2. Tables of estimates (± SE), with *t*-values and their probability levels, for GLS-arima regressions of daily stem girth increment on relative humidity recorded at 14:00 (RH14). (a) Coefficient of single fitted term 'RH14 residuals', taken from first regressions of RH14 on dry-bulb temperature at 14:00 (Tdry14), and (b) Coefficients of the interaction term in the nested model 'Tdry14 + Tdry14·RH14'. RH14 and Tdry14 were recorded at the DVFC climate station.

| | | | (a) Residual RH term from one-term RH model | | | | (b) Interaction term from two-term TEMP and RH model | | | |
|---|---|---|---|---|---|---|---|---|---|---|
| season | gth | spec | est | se | t(est) | P(t) | est | se | t(est) | P(t) |
| w | g14 | Lb | 0.4332 | 0.2694 | 1.068 | 0.109 | 0.01533 | 0.00852 | 1.799 | 0.073 |
| d | g14 | Lb | -0.6477 | 0.2652 | -2.442 | 0.017 | -0.02032 | 0.00769 | -2.641 | 0.010 |

Appendix 2: Fig. S1. Girth increment, *gthi*, of each tree in the wet period plotted against relative humidity (RH14, %), limited to three, thin temperature (Tdry14, °C) 'slices' — for those days on which the DVFC climate station did have recordings of both variables.

### (a) Slice 1 (29.3 – 29.9 deg.C)

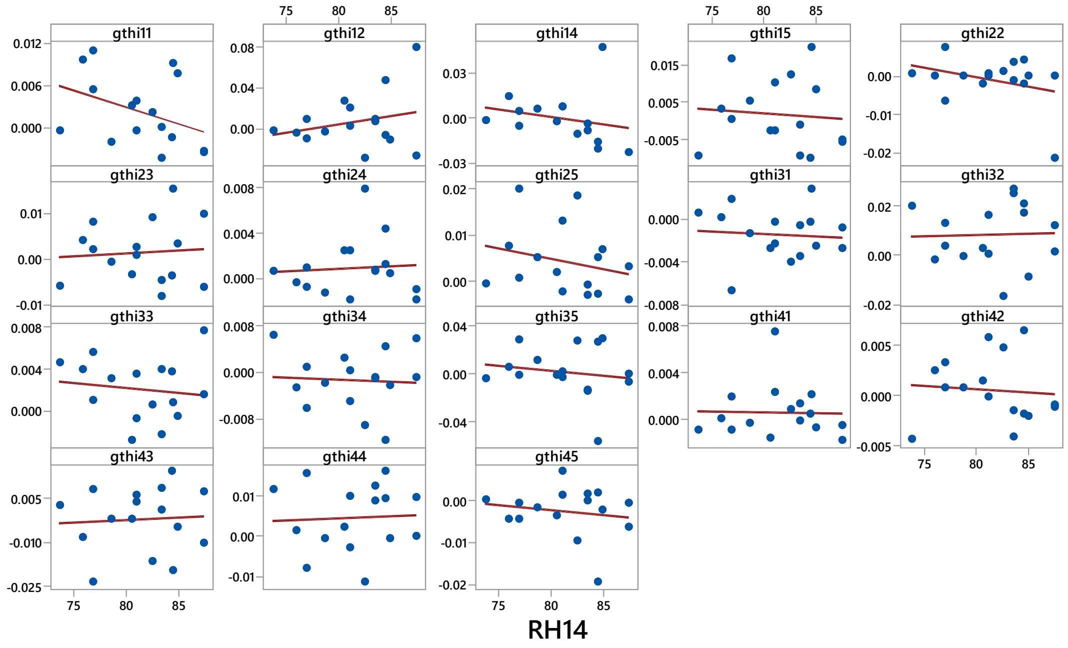


### (b) Slice 2 (30.0 – 30.6 deg.C)

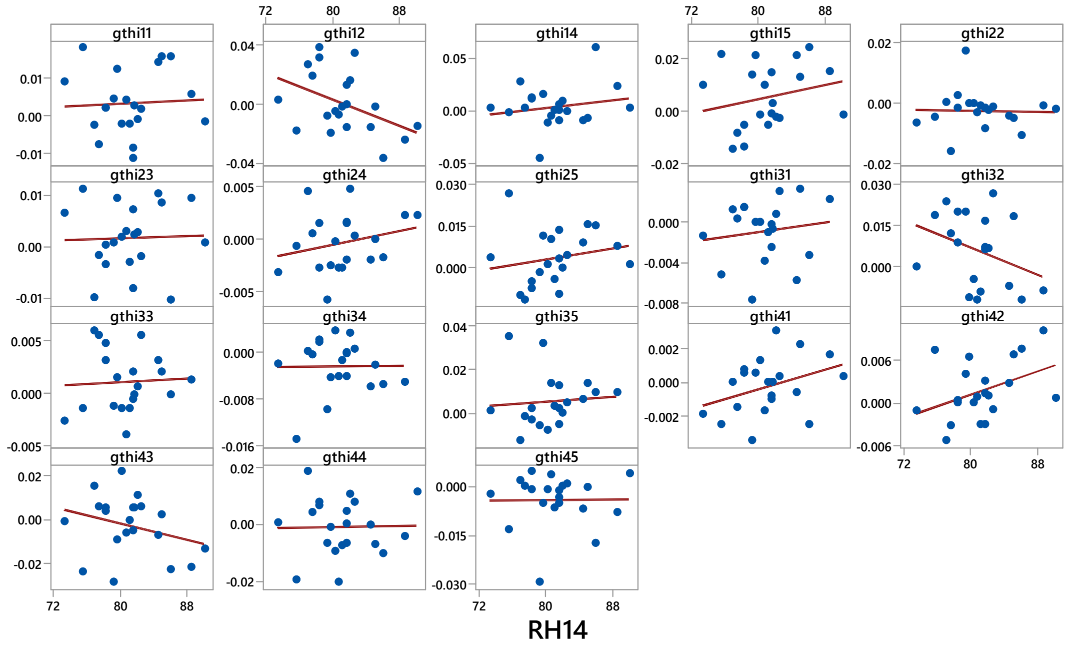

### (c) Slice 3 (30.7 – 31.3 deg.C)

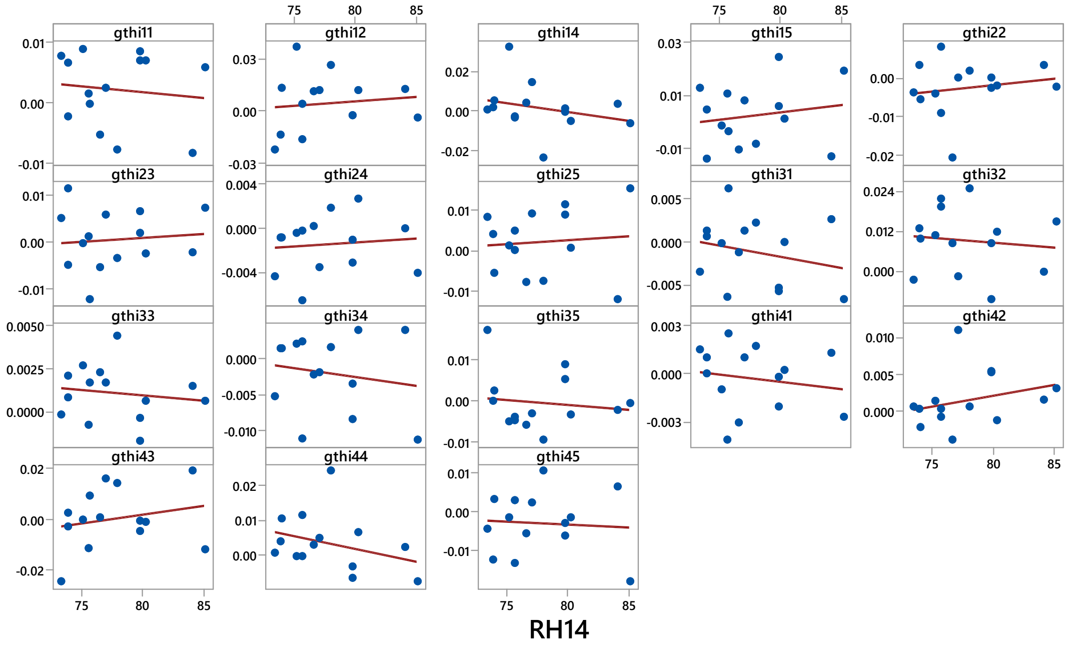

Appendix 3: *Change in dendrometer band length due to temperature change*

The UMS Manual and Data Sheet for the D6 band (UMS, München, Germany) suggest a coefficient of < 1 µm/m/°K for the Invar steel cable and state a 'temperature error' of < 4 µm/°K' for the whole dendrometer. The expansion of the dendrometer is made of two components: (a) the cable itself ($\lambda_1$ = 1 µm/m/°K) and (b) the 'block-and-plate' (expansion value unstated by UMS), which together give the < 4 µm/m/°K overall. Since the cables (bands) are of different lengths on the various trees, a correction for each tree's band can be found for them independently. The 'block-and-plate' (which is the same construction for each band) should, by deduction, account for a < 4 - < 1 = ~ 3 µm/m/°K difference ($\lambda_2$).

Daily temperature differences ($\Delta T = T_{t+1} - T_t$) were found for each tree's band, corresponding to the 1-day increments in *gbh*. When ΔT was positive the dendrometer would have expanded and therefore adjusted the *gthi* should be smaller, and when the ΔT was negative the converse the case. The adjustment was: $gthi_{adj} = gthi - \{((mgbh/100) \cdot \lambda_1 \cdot \Delta T) + (\lambda_2 \cdot \Delta T)\}/k$, where k = 10 is a rescaling factor to convert the adjustment term in µm to cm·$10^{-3}$ (m·$10^{-5}$), the unit for *gthi*. GLS regressions (p = 2, q = 2) were rerun using $ghti_{adj}$, for SMP with no lag, and separately for logger-TEMP with no lag, for the trees/bands in the wet and dry periods. Omitting the four trees where volatility was shown, and two very outlying, low, SMP- (slopes < 1.0) and two logger-TEMP-values (slopes < 1.5), regressions for the *n* = 25 species' adjusted versus unadjusted slope-values (estimates for wet {14} and dry {11} periods together) were very strong ($R^2$ = 99.8 and 99.4% respectively, and these two regression slopes very close to unity {0.978 and 1.015} with intercepts very near zero). The two outliers were also close to the lines for the 25 points but afforded high leverages).

Could diurnal changes in the bands have been in part due to daily temperature change? From the DVMC climate station, dry-bulb temperatures at 08:00 and 14:00 h in the wet period were 24.3 and 30.0 °C on average, and in the dry period correspondingly 25.3 and

31.7 ºC; ΔT being then 5.7 and 6.4 ºC for wet and dry periods. Although there are no records for temperatures at DVMC at midnight, Appendix 1: Fig. S4 shows for example that minimums were usually around 08:00 h. However, the *gthch* variable was calculated with 00:00 h as reference, and the mean logger temperatures at this time (t = 1) and the maximum at station 1 were 23.5 and 26.5 ºC in the wet, and 24.6 and 28.7 ºC in the dry, period; ΔT being 3.0 and 4.1 ºC for the two periods. Logger temperatures at 00:00 h were 1.0 to 1.1 ºC higher that those at 08:00 h. In the daytime the temperature sensor would, furthermore, have been somewhat insulated, from especially the higher temperature extremes, within the logger housing. The temperature curves for the other three stations were very similar. Taken together an average upper maximum diurnal ΔT experienced by the bands can be estimated at ~5.5 º C. Applying this in the formula for band expansion given above, a tree of 30 cm *gbh* would have maximally expanded 0.62 *gthch* units, and one of 100 cm *gbh* by 0.22 (the change being relative to *gbh* at $t_1$; most of the expansion in the 'block-and-plate'). These are very small values when judged against diurnal band fluctuations. Temperature change therefore affected the species' diurnal *gthch* patterns very minimally, at least as far as the bands themselves were concerned.

Appendix 4: Table S1. Statistics for the quadratic regression fits of daily girth increment (gthi) of dendrobands on 13 trees as a function of rainfall in the current, and in the previous, day (on ln-scale) in the dry period, for days with ≥ 1 mm in the day. Explanation of the series is found in the main text. Probability values ≤ 0.02 are shown in bold font. Bands where the relationship is significantly stronger for the previous than current day are indicated in red.

| series | band | code | Gbh | Rain on current day | | | Rain on previous day | | |
|---|---|---|---|---|---|---|---|---|---|
| | | | (cm) | $R^{2\|a}$ | $F^b$ | P(F) | $R^{2\|a}$ | $F^c$ | P(F) |
| A(B) | g11 | Mw | 31 | 0.332 | 15.65 | **<0.001** | 0.071 | 3.22 | 0.048 |
| | g12 | Sf | 97 | 0.034 | 2.03 | 0.141 | 0.235 | 9.92 | **<0.001** |
| | g14 | Lb | 31 | 0.170 | 7.05 | **0.002** | –0.007 | 0.80 | 0.457 |
| | g15 | Sf | 27 | 0.345 | 16.52 | **<0.001** | 0.003 | 1.09 | 0.344 |
| B | g22 | Pm | 74 | 0.148 | 5.67 | **0.006** | 0.069 | 2.97 | 0.060 |
| | g23 | Dm | 30 | 0.137 | 5.28 | **0.008** | –0.030 | 0.24 | 0.790 |
| | g24 | Mw | 34 | 0.308 | 13.02 | **<0.001** | –0.013 | 0.65 | 0.527 |
| | g25 | Sf | 33 | 0.014 | 1.37 | 0.263 | –0.024 | 0.38 | 0.689 |
| | g31 | Mw | 34 | 0.249 | 9.95 | **<0.001** | –0.021 | 0.44 | 0.647 |
| | g32 | Sp | 78 | 0.097 | 3.92 | 0.026 | –0.022 | 0.43 | 0.653 |
| | g33 | Lb | 39 | –0.033 | 0.13 | 0.876 | 0.440 | 21.81 | **<0.001** |
| | g34 | Dm | 39 | 0.460 | 23.97 | **<0.001** | 0.002 | 1.07 | 0.352 |
| | g35 | Dm | 48 | 0.132 | 5.10 | **0.009** | 0.006 | 1.16 | 0.322 |

a, adjusted $R^2$-value; df: [b], A(B) 2, 57; B 2, 52. [c] , A(B) 2,56; B 2, 51. [The 1 df less for the lagged regression error term is because the day before the starts of the series had no rain.]

Appendix  4: Fig. S1.

Daily girth increment (gthi) as a function of rainfall in the current day,'rain', (on ln-scale) in the dry period. The 13 bands are presented pairwise in rows: on the left all days (i.e. those with zero rain included) are shown with a 'lowess' regression fit, and on the right for those days receiving ≥ 1 mm rainfall with a quadratic regression fit.

Appendix 4: Fig. S2.

Daily girth increment (gthi) as a function of rainfall in the day before, 'rain_1', (on ln-scale) in the dry period. The 13 bands are presented pairwise in rows: on the left all days (i.e. those with zero rain included) are shown with a 'lowess' regression fit, and on the right for those days receiving ≥ 1 mm rainfall with a quadratic regression fit.

Fig. S1

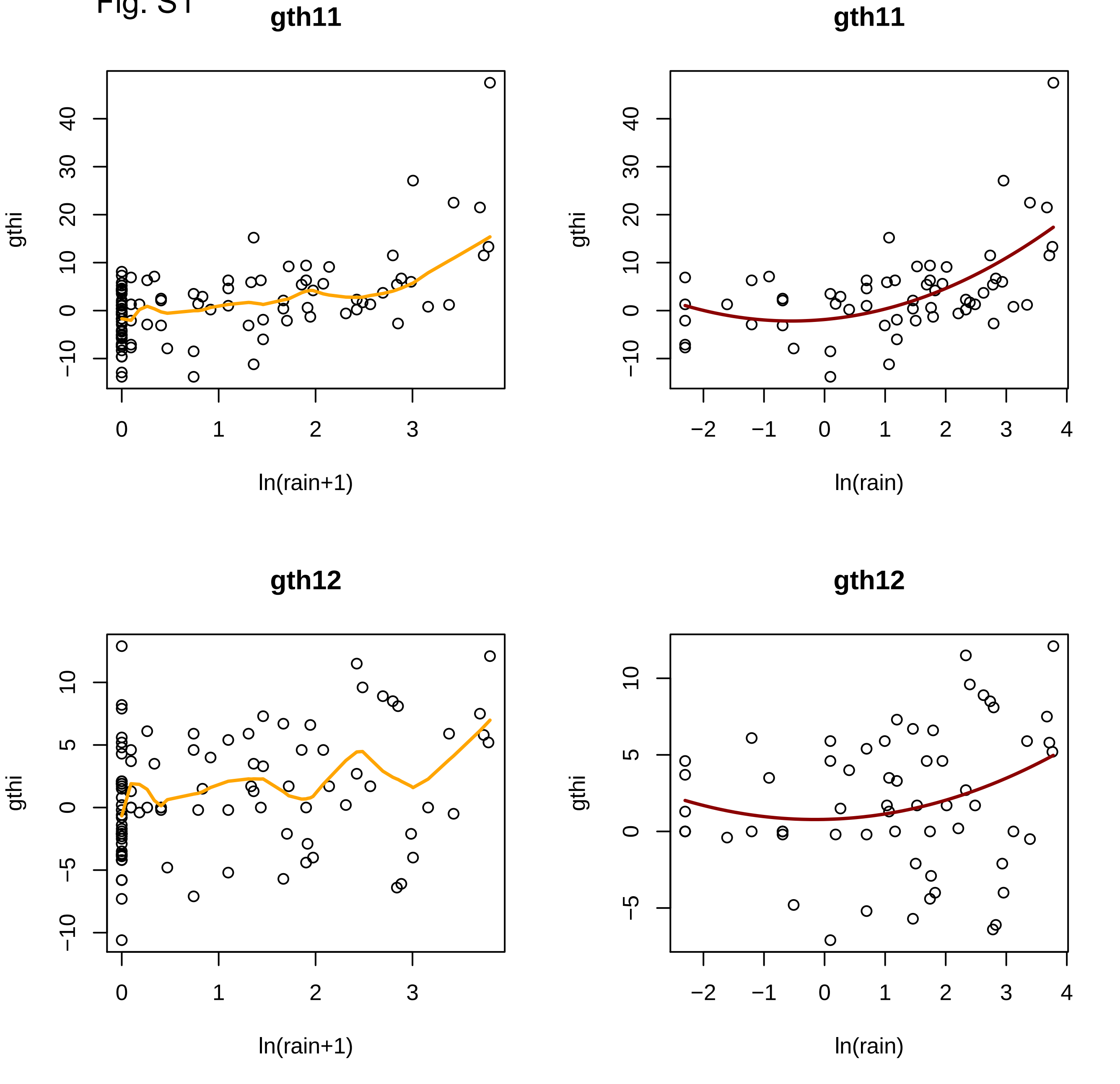

gth11
gthi
ln(rain+1)
gth11
gthi
ln(rain)
gth12
gthi
ln(rain+1)
gth12
gthi
ln(rain)

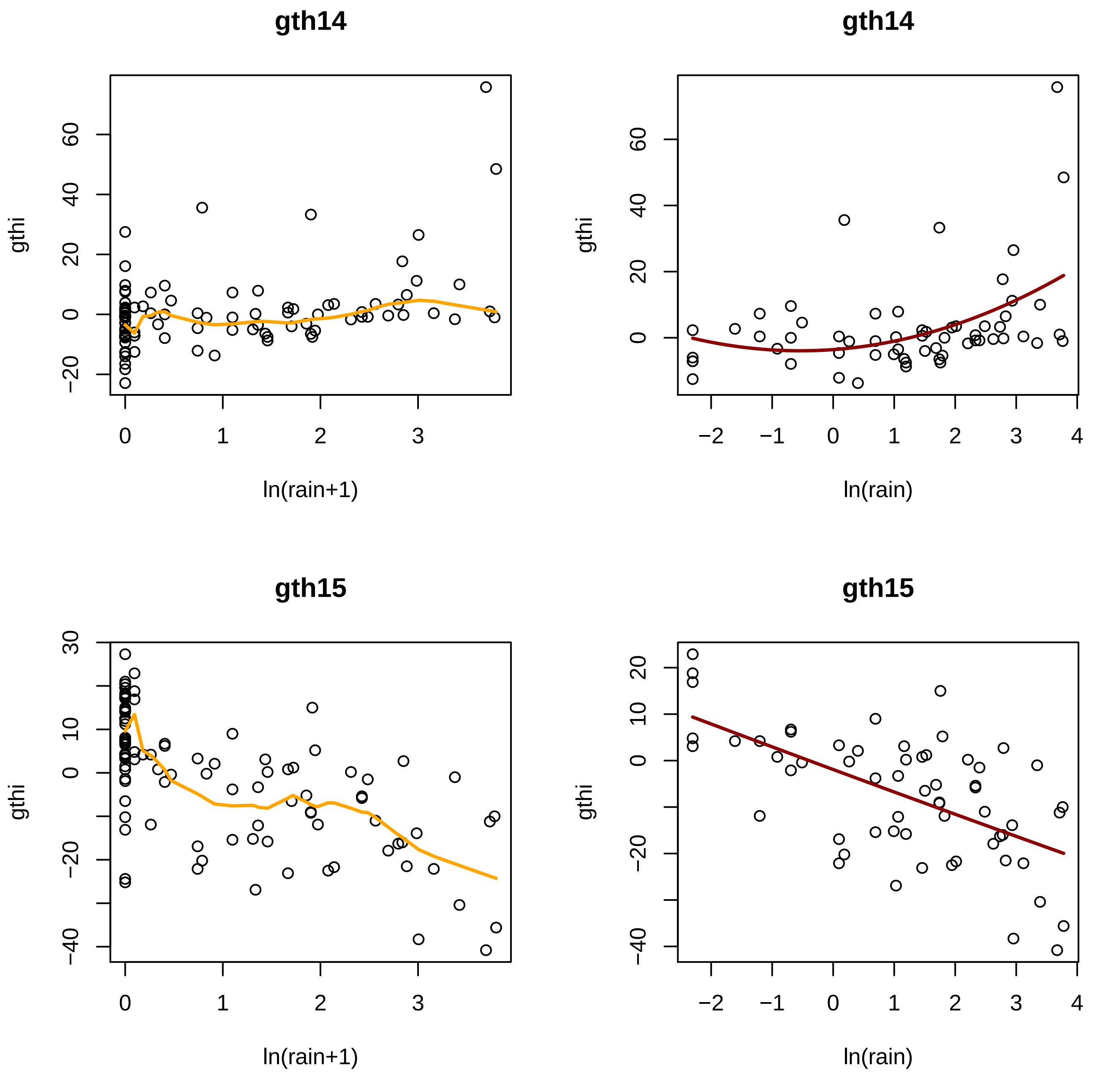
gth14
gthi
ln(rain+1)
gth14
gthi
ln(rain)
gth15
gthi
ln(rain+1)
gth15
gthi
ln(rain)

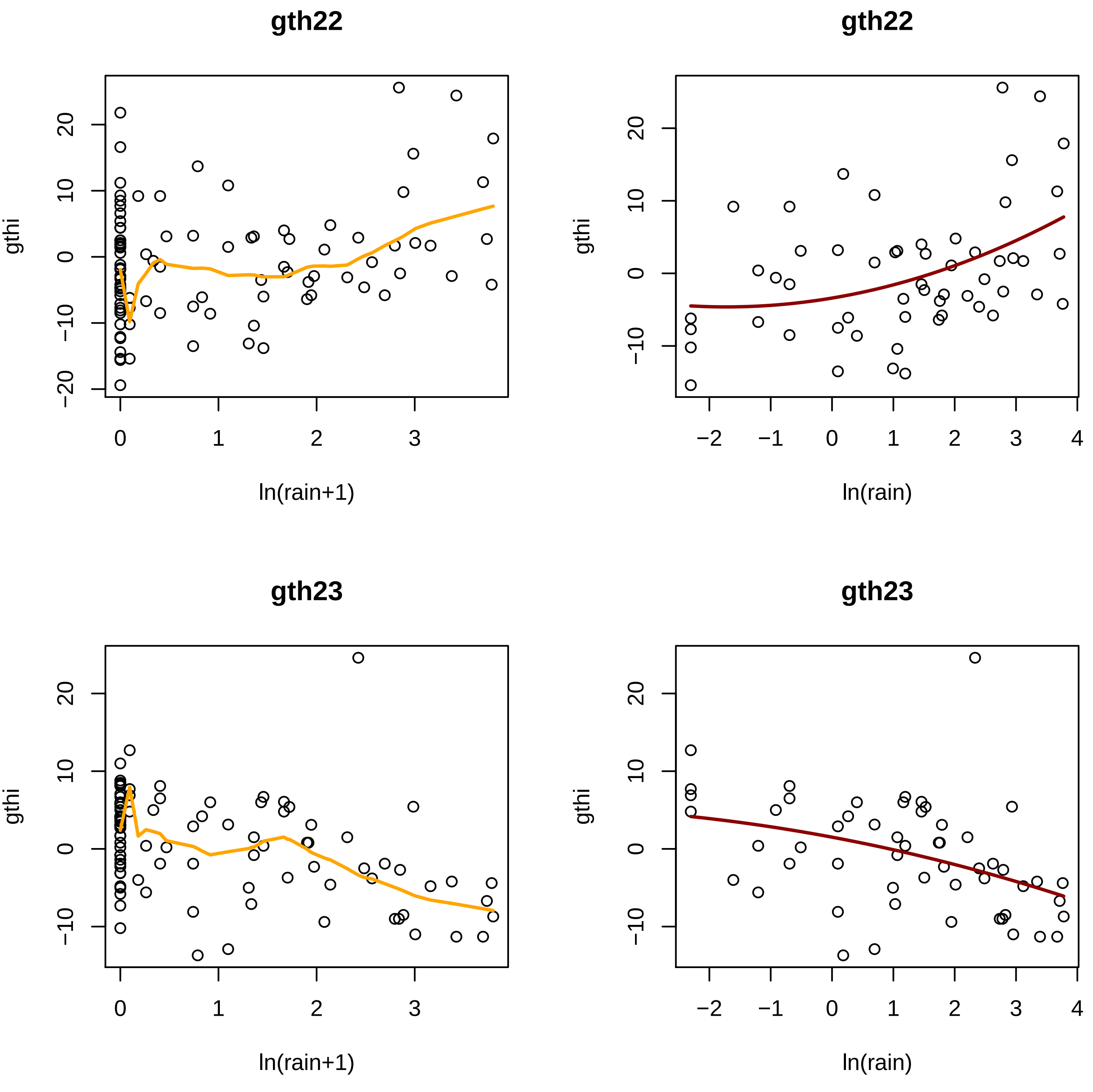

gth22
gth22
gth23
gth23
gthi
ln(rain+1)
ln(rain)

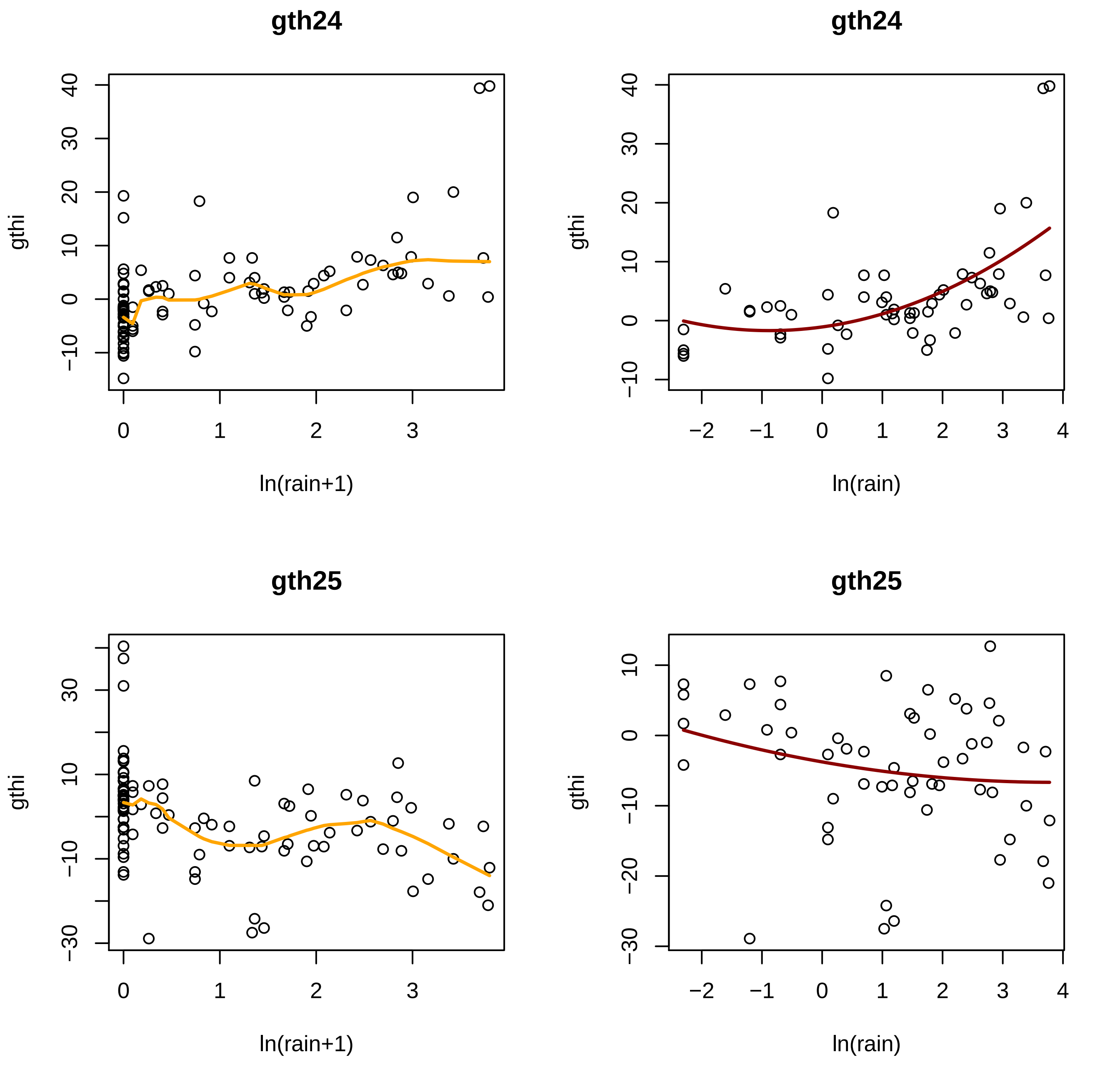
gth24
gthi
ln(rain+1)
gth24
gthi
ln(rain)
gth25
gthi
ln(rain+1)
gth25
gthi
ln(rain)

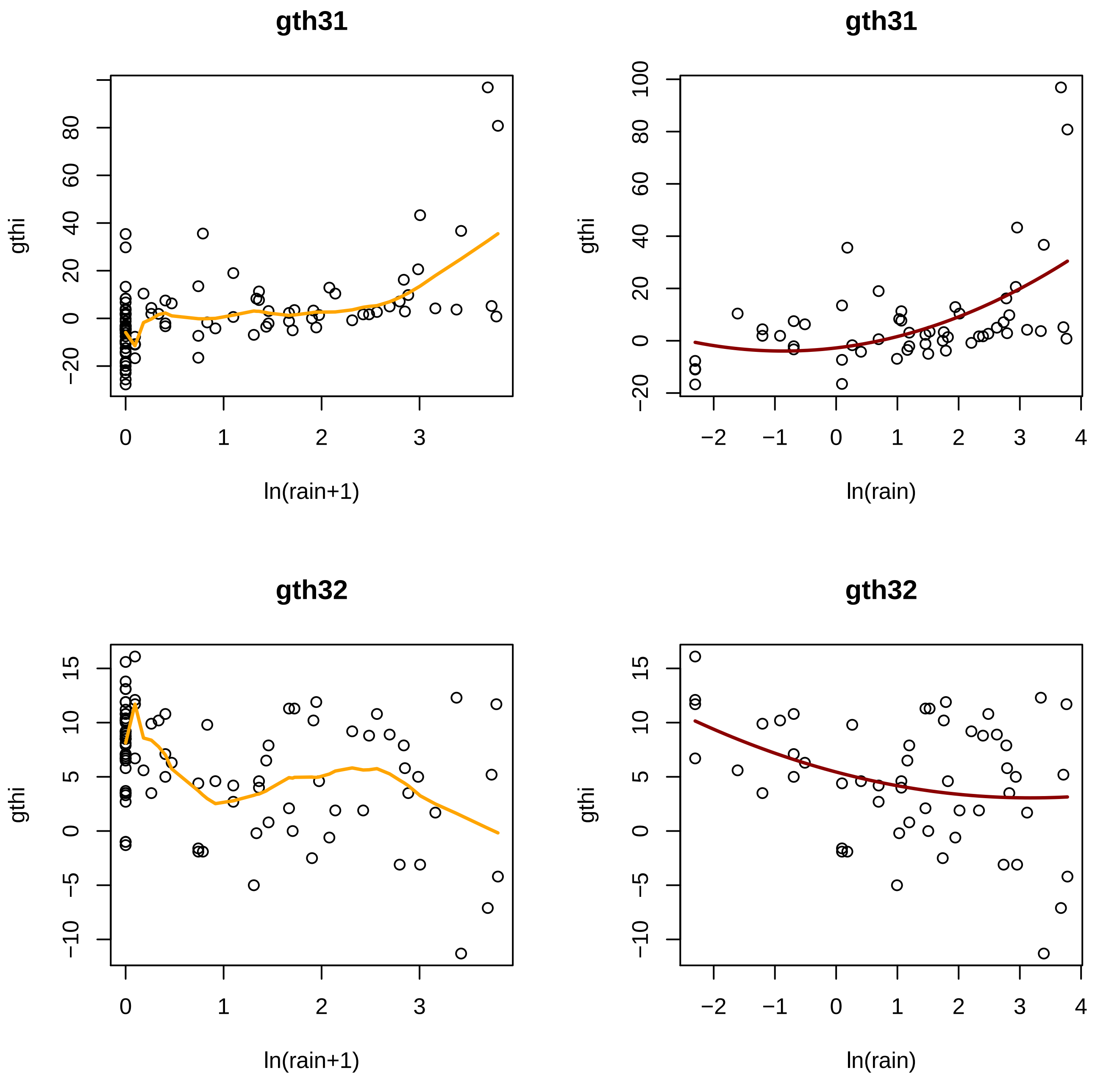
gth31
gth31
gth32
gth32
gthi
ln(rain+1)
ln(rain)

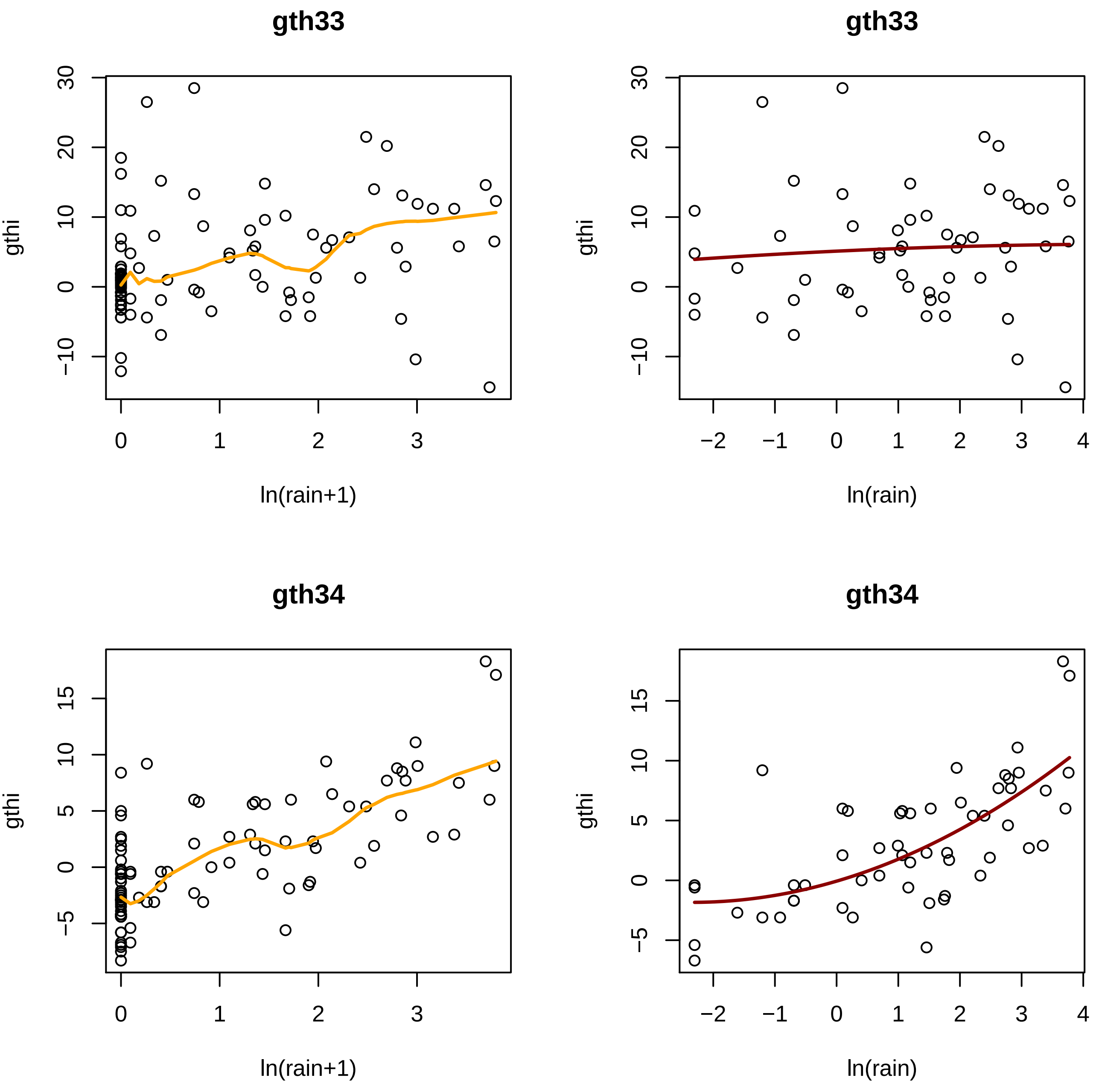
gth33
gthi
ln(rain+1)
30
20
10
0
−10
0
1
2
3
gth33
gthi
ln(rain)
−2
−1
4
gth34
gthi
ln(rain+1)
15
5
−5
gth34
gthi
ln(rain)

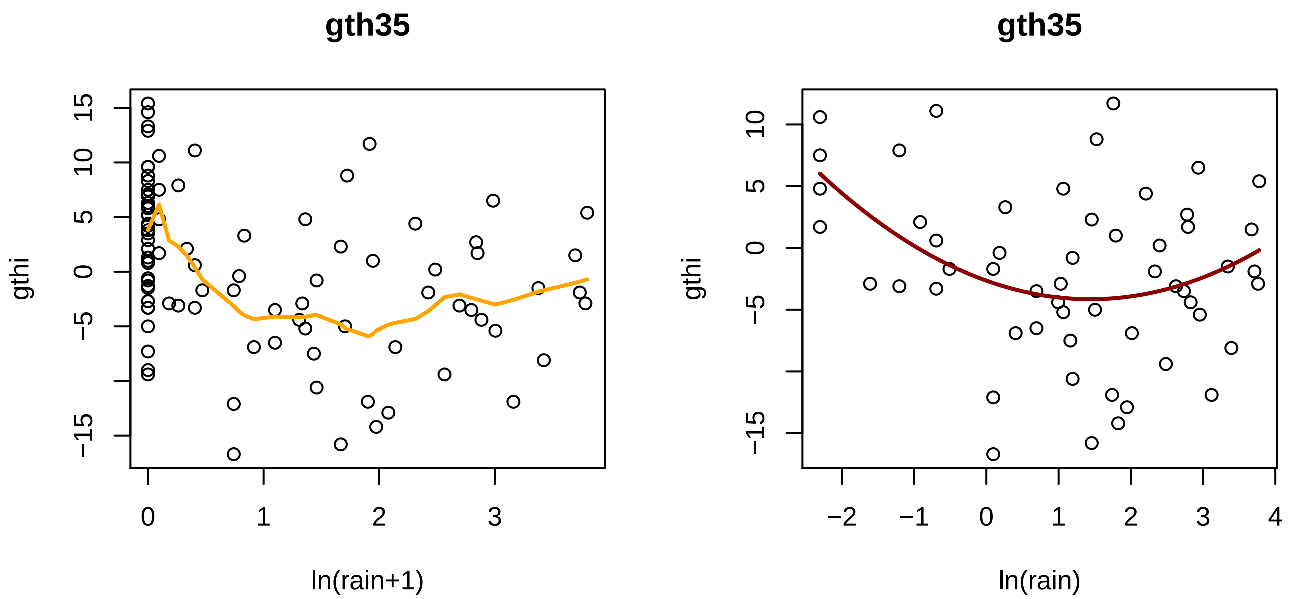
gth35
gthi
ln(rain+1)
gth35
gthi
ln(rain)

Fig. S2

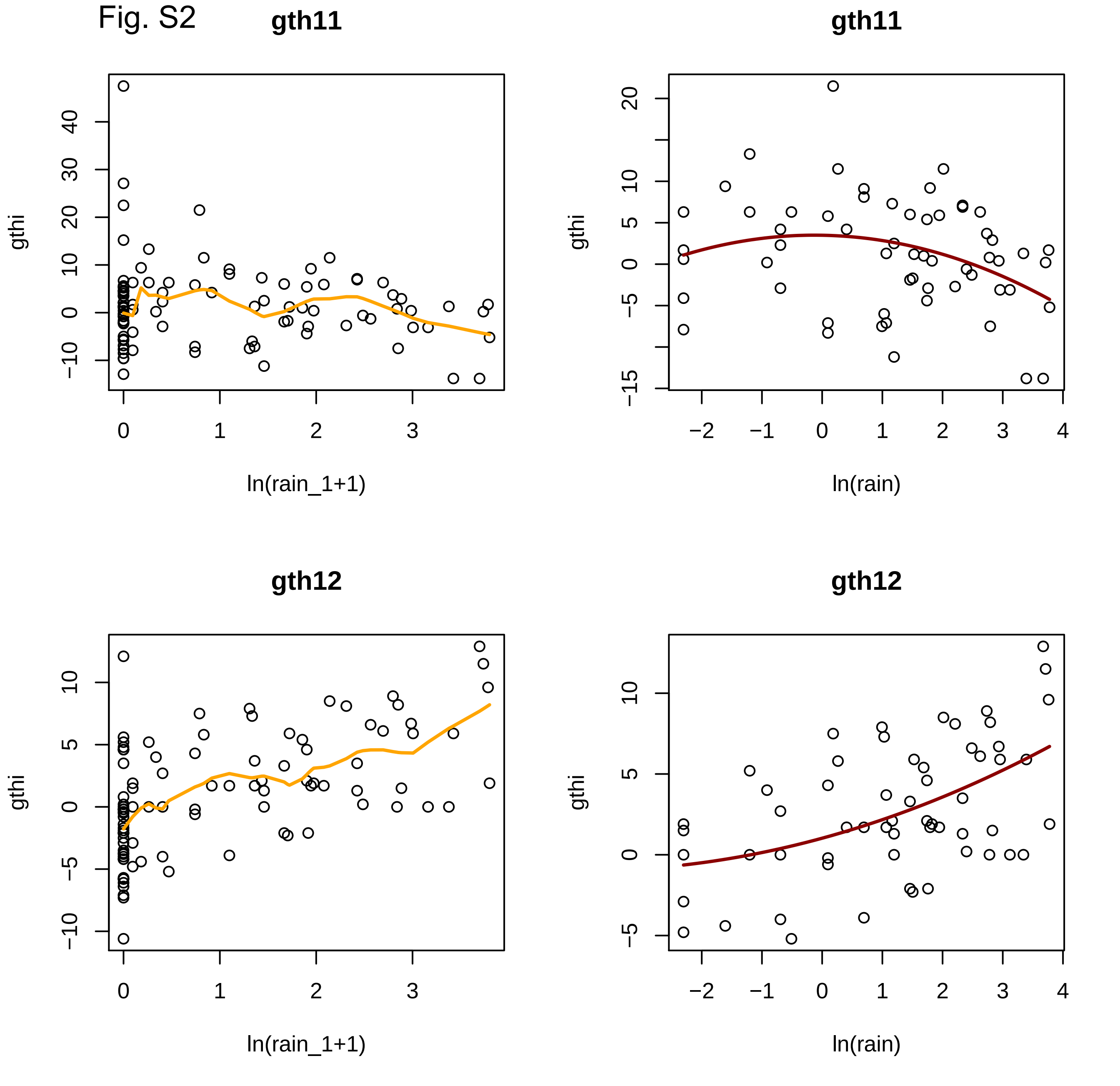
gth11
gthi
ln(rain_1+1)
gth11
gthi
ln(rain)
gth12
gthi
ln(rain_1+1)
gth12
gthi
ln(rain)

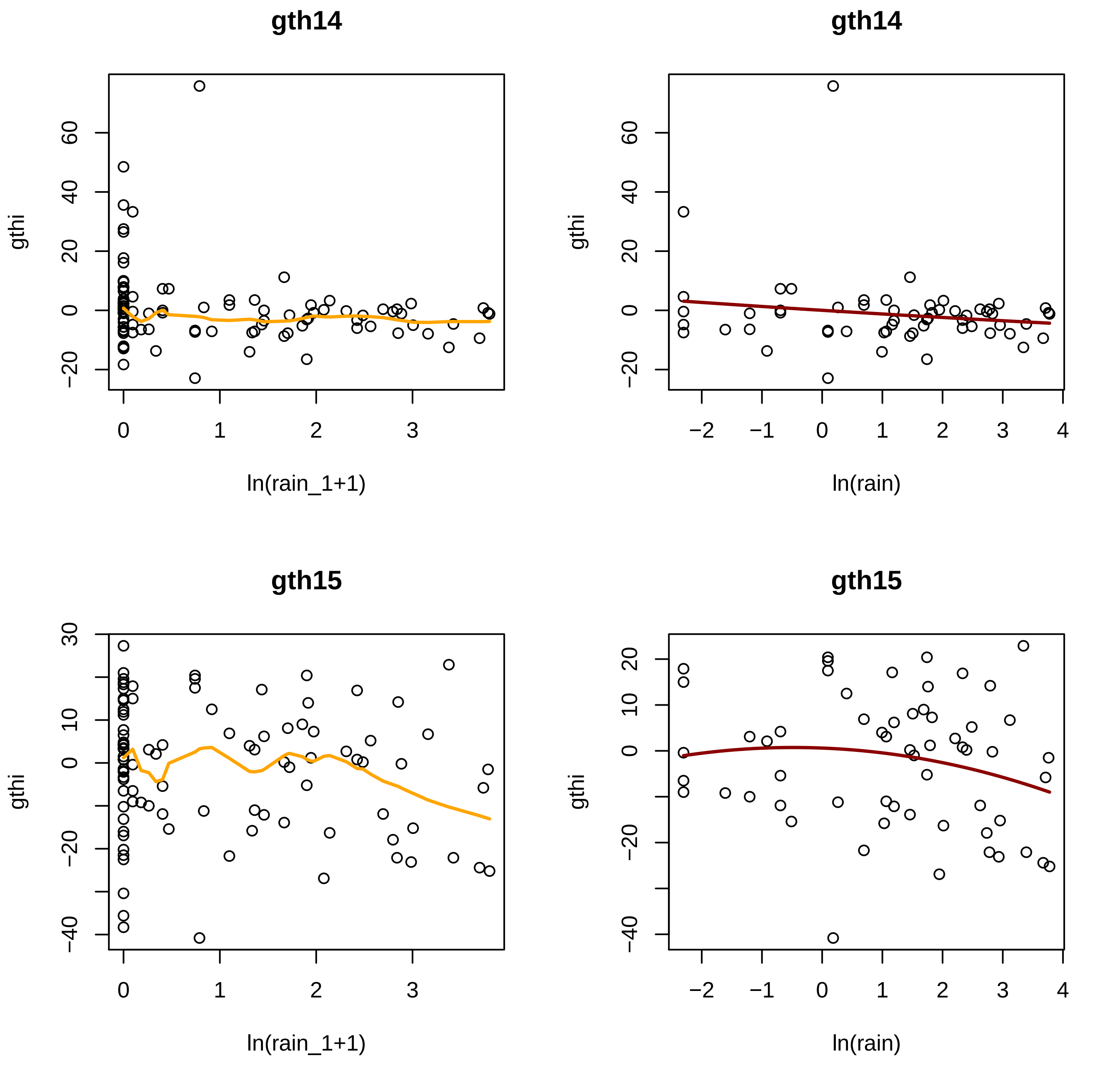
gth14
gthi
ln(rain_1+1)
gth14
gthi
ln(rain)
gth15
gthi
ln(rain_1+1)
gth15
gthi
ln(rain)
−20
0
20
40
60
−40
−20
0
10
30
−2
−1
0
1
2
3
4

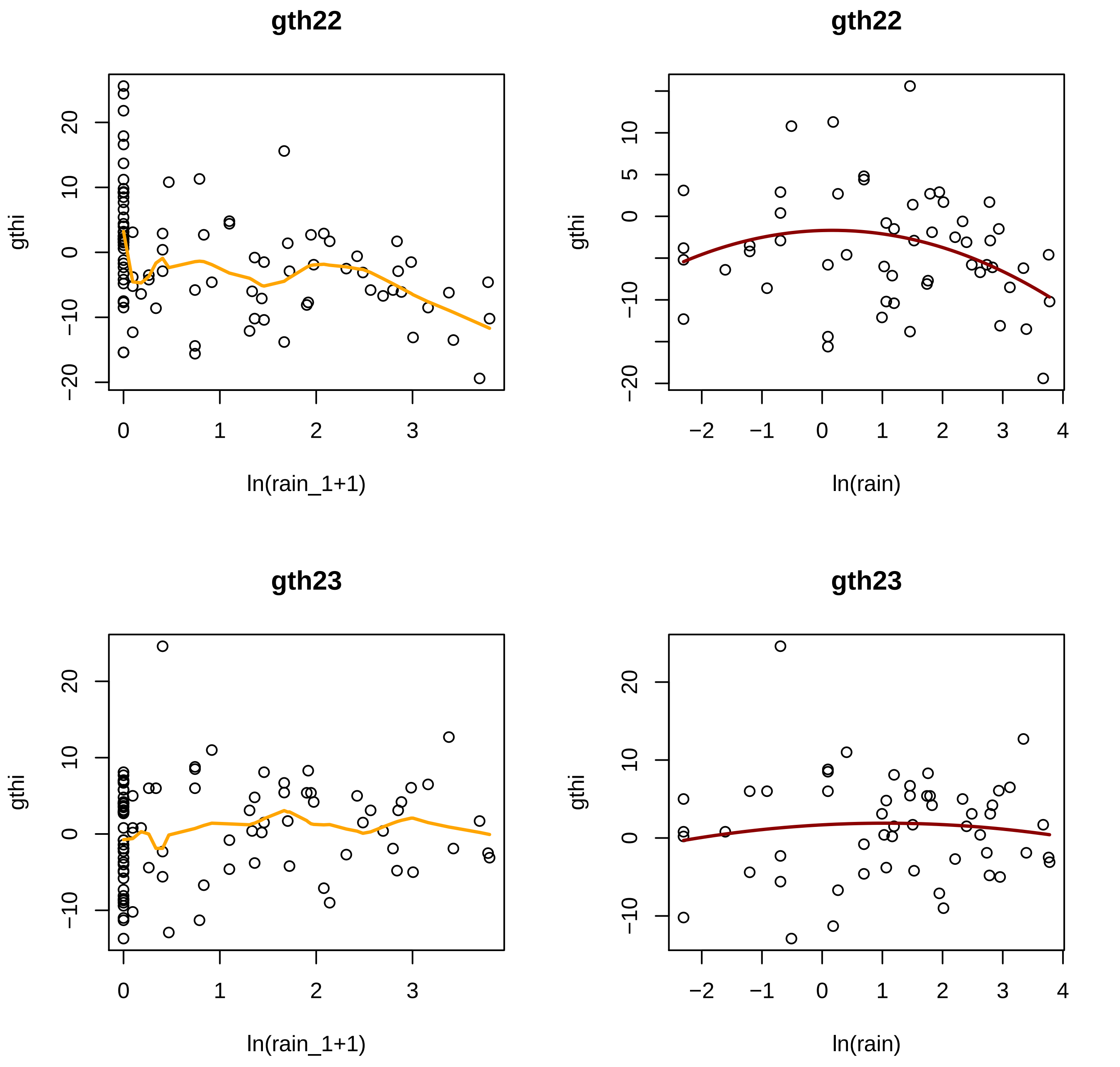
gth22
gthi
ln(rain_1+1)
gth22
gthi
ln(rain)
gth23
gthi
ln(rain_1+1)
gth23
gthi
ln(rain)

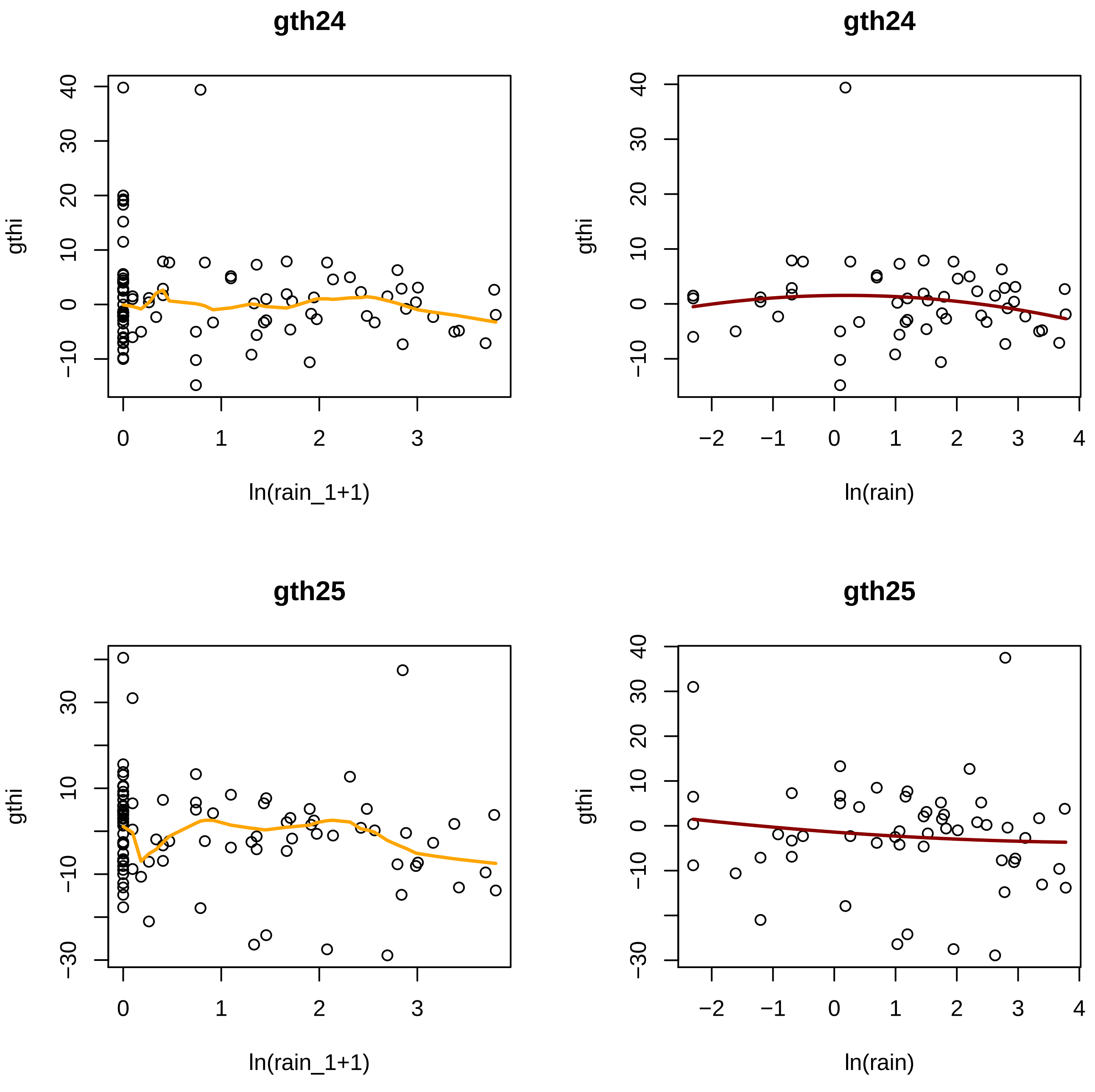
gth24
gth24
gth25
gth25
gthi
ln(rain_1+1)
ln(rain)

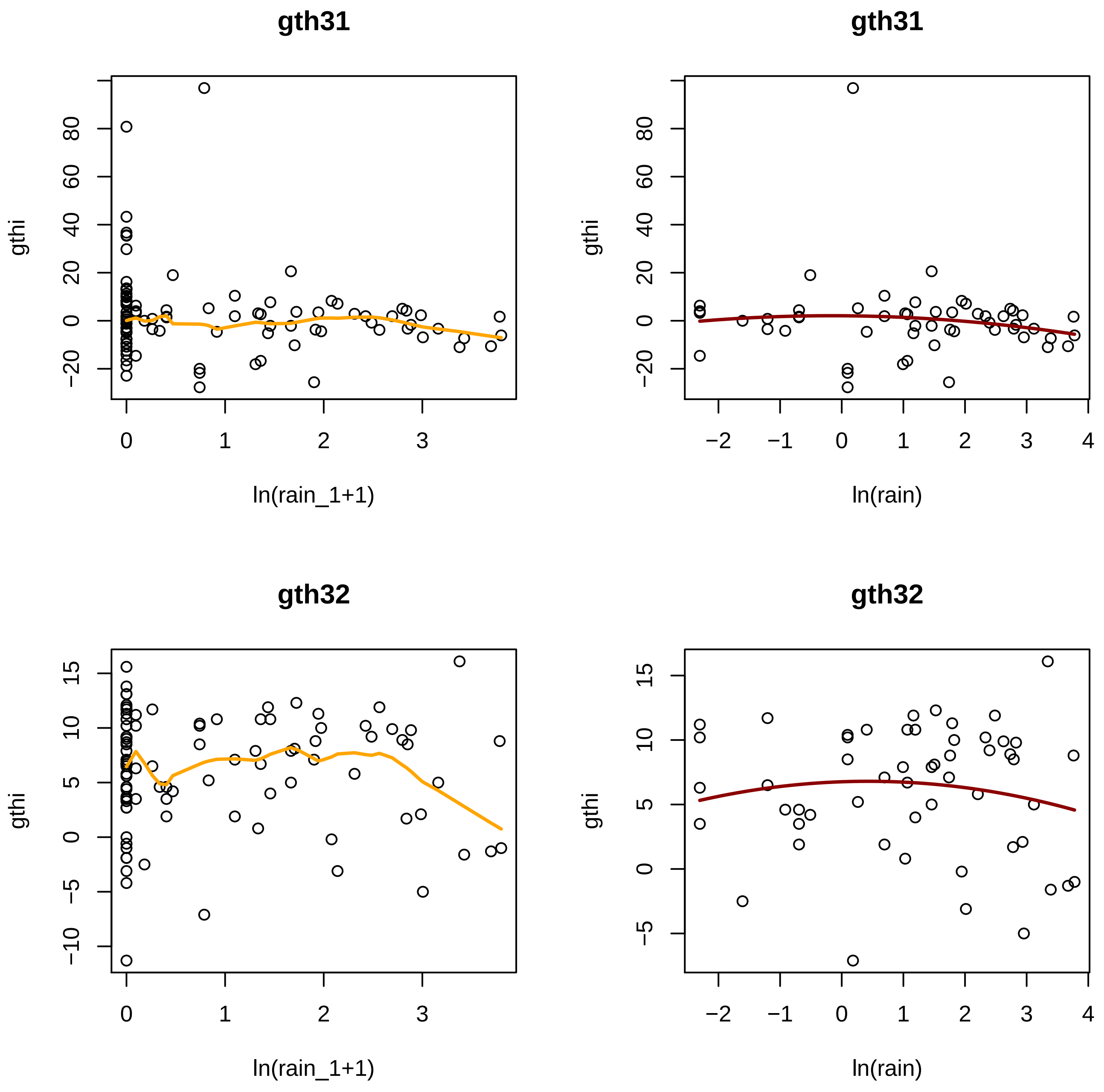
gth31
gthi
ln(rain_1+1)
gth31
gthi
ln(rain)
gth32
gthi
ln(rain_1+1)
gth32
gthi
ln(rain)

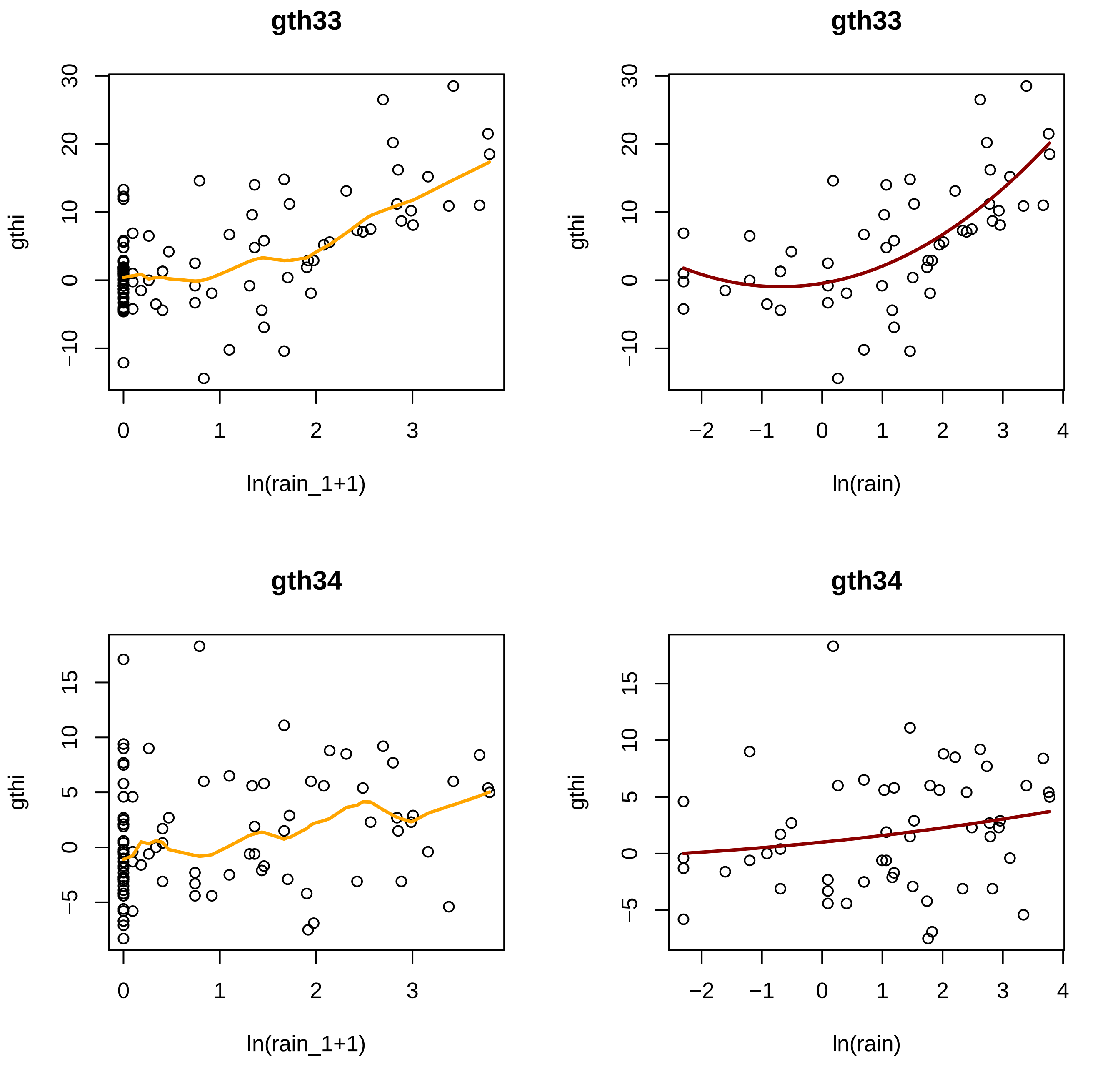
gth33
gthi
ln(rain_1+1)
gth33
gthi
ln(rain)
gth34
gthi
ln(rain_1+1)
gth34
gthi
ln(rain)

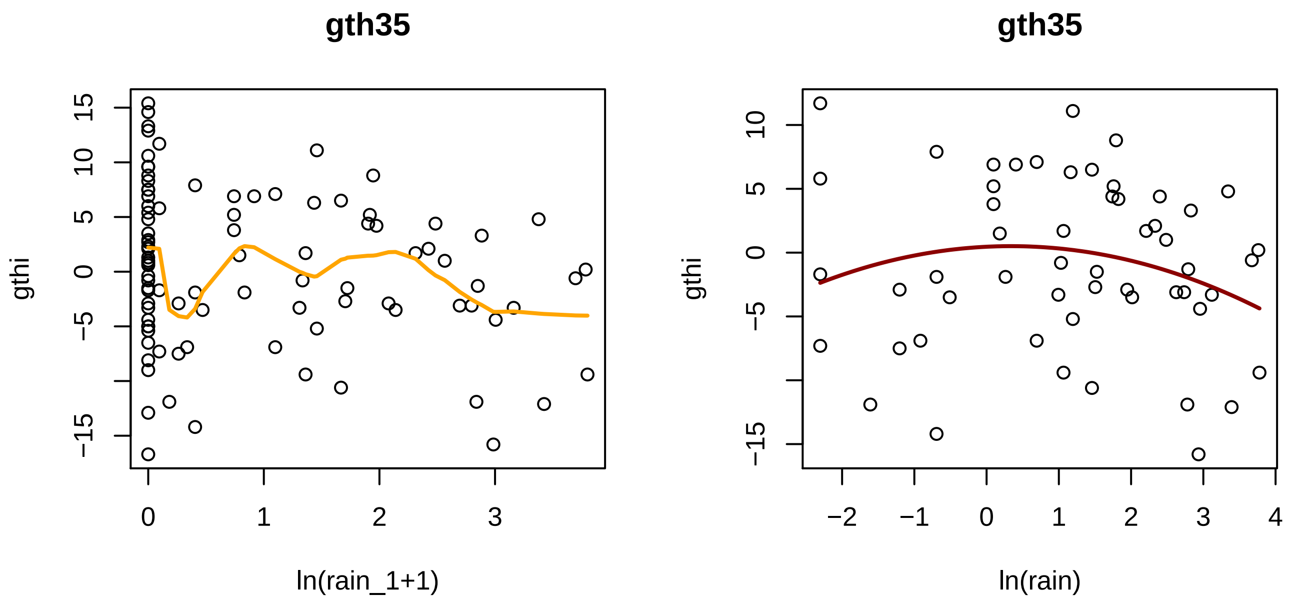

gth35
gthi
ln(rain_1+1)
gth35
gthi
ln(rain)

Appendix 4: Fig. S3. Estimates of (a) SMP, and (b) TEMP, coefficients from the GLS-arima single-term model fits for the 13 dendrobands recorded in the dry period, each for three lags (0, 1 and 2 days), plotted for analyses that used the original daily growth increments, *gthi*, ('dry') versus those that used the residuals from fitting *gthi* to rainfall ('rdry'). The two most important outliers in each case are identified.

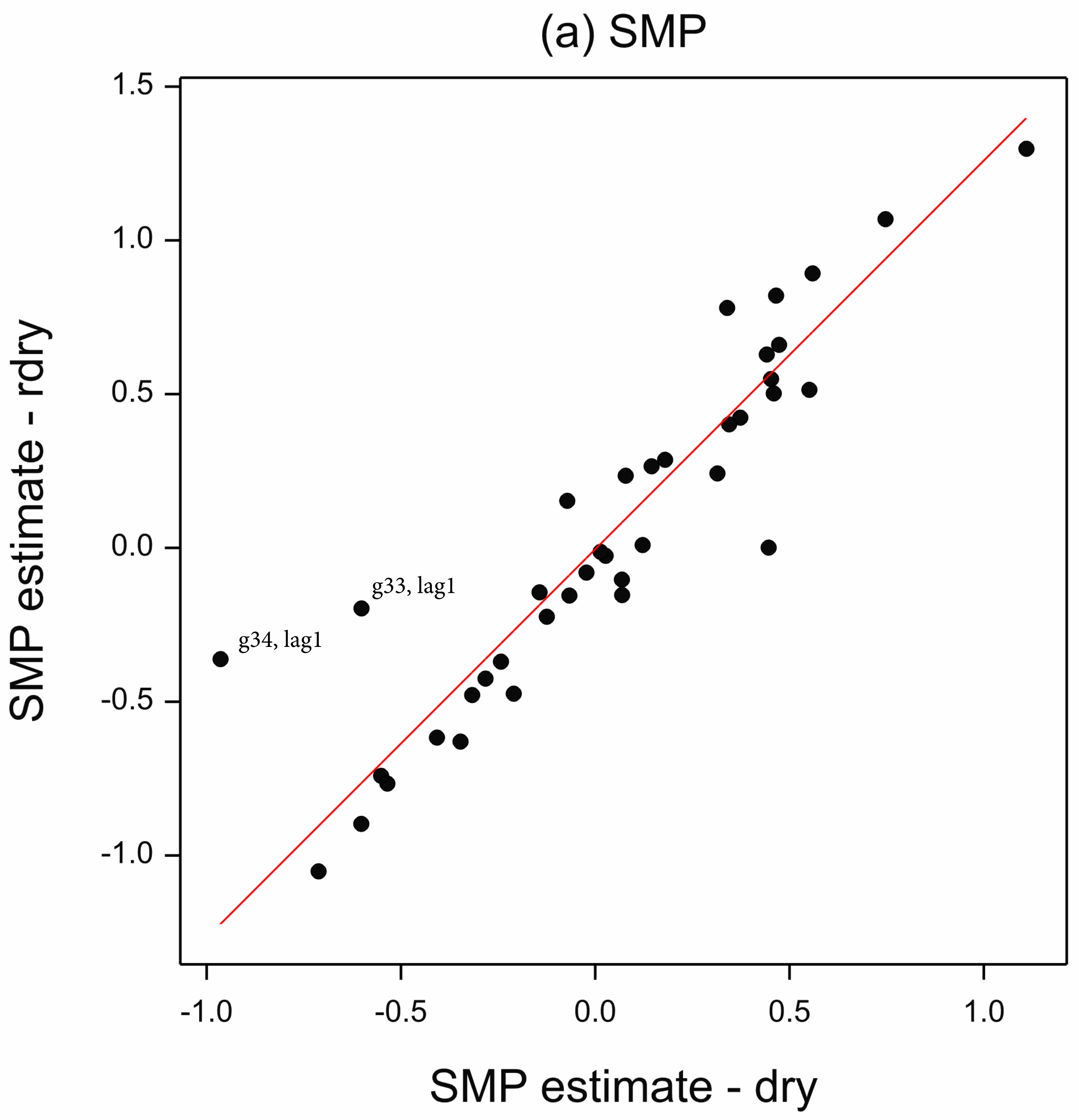


Appendix 4: Fig. S3. Estimates of (a) SMP, and (b) TEMP, coefficients from the GLS-arima single-term model fits for the 13 dendrobands recorded in the dry period, each for three lags (0, 1 and 2 days), plotted for analyses that used the original daily growth increments, gthi, ('dry') versus those that used the residuals from fitting gthi to rainfall ('rdry'). The two most important outliers in each case are identified.

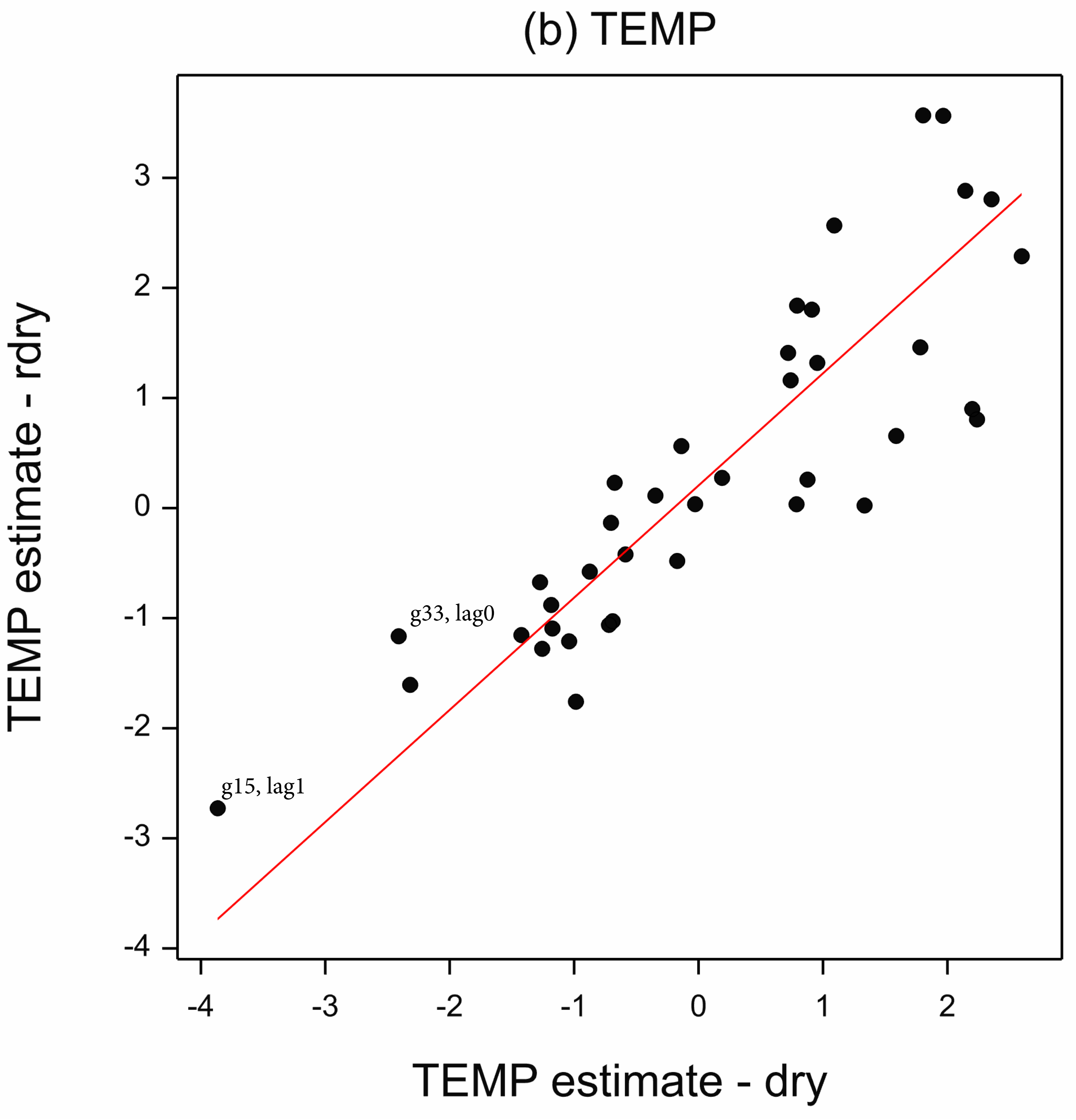

(b) TEMP
TEMP estimate - rdry
TEMP estimate - dry
g33, lag0
g15, lag1
3
2
1
0
-1
-2
-3
-4
-4
-3
-2
-1
0
1
2

Appendix 5: Table S1. Tables of *t*-values and their probability levels for three time lags (0, 1, and 2 days) for GLS-regression coefficients (slopes) of daily stem girth increment versus soil moisture potential (SMP). In these regression fits SMP was the only independent variable. The coefficients and their standard errors are represented in Fig. 4 of the main paper.

| season | gth | spec | t(est)_0 | P(t)_0 | t(est)_1 | P(t)_1 | t(est)_2 | P(t)_2 |
|---|---|---|---|---|---|---|---|---|
| w | g11 | Mw | 3.848 | 0.00013 | 6.963 | <0.00001 | 5.359 | <0.00001 |
| w | g12 | Sf | 0.673 | 0.50102 | −5.080 | <0.00001 | −2.071 | 0.03879 |
| w | g14 | Lb | 0.150 | 0.44100 | −0.277 | 0.39100 | 0.216 | 0.41400 |
| w | g15 | Sf | 2.464 | 0.01406 | 6.652 | <0.00001 | 5.371 | <0.00001 |
| w | g22 | Pm | −7.239 | <0.00001 | −6.921 | <0.00001 | −5.697 | <0.00001 |
| w | g23 | Dm | 0.311 | 0.37800 | 3.071 | 0.00100 | 2.434 | 0.00800 |
| w | g24 | Mw | −2.710 | 0.00693 | −2.128 | 0.03375 | −1.289 | 0.19803 |
| w | g25 | Sf | 1.735 | 0.08324 | 6.190 | <0.00001 | 4.667 | <0.00001 |
| w | g31 | Mw | −4.140 | 0.00004 | −4.180 | 0.00003 | −3.752 | 0.00019 |
| w | g32 | Sp | −0.289 | 0.77268 | −0.293 | 0.76926 | −0.170 | 0.86508 |
| w | g33 | Lb | −4.972 | <0.00001 | −4.365 | 0.00002 | −2.450 | 0.01460 |
| w | g34 | Dm | −3.237 | 0.00128 | −4.288 | 0.00002 | −3.848 | 0.00013 |
| w | g35 | Dm | −0.216 | 0.82876 | 4.288 | 0.00002 | 4.464 | 0.00001 |
| w | g41 | Sf | −2.404 | 0.00900 | −2.736 | 0.00300 | −2.662 | 0.00400 |
| w | g42 | Dm | 4.069 | 0.00005 | 6.917 | <0.00001 | 6.589 | <0.00001 |
| w | g43 | Pm | −5.920 | <0.00001 | −5.884 | <0.00001 | −5.673 | <0.00001 |
| w | g44 | Sp | −6.367 | <0.00001 | −8.258 | <0.00001 | −5.722 | <0.00001 |
| w | g45 | Pm | −4.508 | 0.00001 | −5.458 | <0.00001 | −4.787 | <0.00001 |
| d | g11 | Mw | −1.176 | 0.24254 | −0.825 | 0.41135 | 0.079 | 0.93696 |
| d | g12 | Sf | 0.291 | 0.77178 | −0.268 | 0.78921 | −0.872 | 0.38520 |
| d | g14 | Lb | −2.851 | 0.00533 | −2.516 | 0.01355 | −2.718 | 0.00780 |
| d | g15 | Sf | 2.335 | 0.02162 | 5.258 | <0.00001 | 5.035 | <0.00001 |
| d | g22 | Pm | −2.599 | 0.01090 | −1.101 | 0.27367 | −0.093 | 0.92650 |
| d | g23 | Dm | 3.155 | 0.00217 | 3.320 | 0.00130 | 1.492 | 0.13921 |
| d | g24 | Mw | −3.838 | 0.00023 | −3.816 | 0.00025 | −3.477 | 0.00078 |
| d | g25 | Sf | 4.995 | <0.00001 | 7.109 | <0.00001 | 6.225 | <0.00001 |
| d | g31 | Mw | −4.416 | 0.00003 | −4.117 | 0.00008 | −3.595 | 0.00053 |
| d | g32 | Sp | 4.034 | 0.00011 | 6.049 | <0.00001 | 4.945 | <0.00001 |
| d | g33 | Lb | 0.006 | 0.99555 | −1.128 | 0.26210 | 0.998 | 0.32082 |
| d | g34 | Dm | −1.254 | 0.21308 | −3.205 | 0.00186 | −2.111 | 0.03750 |
| d | g35 | Dm | 2.384 | 0.01920 | 3.006 | 0.00342 | 2.921 | 0.00440 |

Coefficients significant at the familywise α ≤ 0.05, 0.01 and 0.002 levels are background colored in increasing intensities of green for the wet series and orange for the dry one. Simple Bonferroni adjustment was applied by dividing the family α-value by the number of bands/trees per series, α/18 and α/13 respectively. The adjusted α's for wet were *P* = 0.00278, 0.00056 and 0.00011; and for dry *P* = 0.00385, 0.00077 and 0.00015.

Appendix 5: Table S2. Tables of *t*-values and their probability levels for three time lags (0, 1, and 2 days) for GLS-regressions coefficients (slopes) of daily girth increment versus ambient temperature (TEMP). In these regression fits TEMP was the only independent variable. The coefficients and their standard errors are represented in Fig. 5 of the main paper.

| season | gth | spec | t(est)_0 | P(t)_0 | t(est)_1 | P(t)_1 | t(est)_2 | P(t)_2 |
|---|---|---|---|---|---|---|---|---|
| w | g11 | Mw | 3.053 | 0.00238 | −5.143 | <0.00001 | −1.658 | 0.09783 |
| w | g12 | Sf | −5.737 | <0.00001 | 1.786 | 0.07462 | −1.383 | 0.16725 |
| w | g14 | Lb | −0.980 | 0.32860 | 0.490 | 0.62440 | −0.461 | 0.64500 |
| w | g15 | Sf | 4.773 | <0.00001 | −4.112 | 0.00005 | −0.514 | 0.60742 |
| w | g22 | Pm | 6.084 | <0.00001 | 2.140 | 0.03293 | 2.585 | 0.01004 |
| w | g23 | Dm | 5.228 | 0.00001 | −2.816 | 0.00541 | −0.482 | 0.63040 |
| w | g24 | Mw | 4.423 | 0.00001 | 1.076 | 0.28239 | 1.219 | 0.22347 |
| w | g25 | Sf | 6.309 | <0.00001 | −5.214 | <0.00001 | 0.444 | 0.65739 |
| w | g31 | Mw | 4.869 | <0.00001 | 1.077 | 0.28185 | 3.234 | 0.00129 |
| w | g32 | Sp | 1.485 | 0.13811 | −0.416 | 0.67789 | −1.753 | 0.08008 |
| w | g33 | Lb | 6.324 | <0.00001 | 0.934 | 0.35072 | 1.591 | 0.11212 |
| w | g34 | Dm | 0.891 | 0.37355 | 1.013 | 0.31146 | 1.985 | 0.04761 |
| w | g35 | Dm | 6.091 | <0.00001 | −0.750 | 0.45387 | −0.114 | 0.90965 |
| w | g41 | Sf | 3.037 | 0.00275 | 0.057 | 0.95460 | 1.357 | 0.17660 |
| w | g42 | Dm | 7.662 | <0.00001 | −4.963 | <0.00001 | −3.780 | 0.00017 |
| w | g43 | Pm | 3.047 | 0.00242 | −0.172 | 0.86375 | 1.337 | 0.18176 |
| w | g44 | Sp | 1.880 | 0.06064 | 1.906 | 0.05714 | 2.691 | 0.00733 |
| w | g45 | Pm | 1.564 | 0.11838 | 2.439 | 0.01504 | 2.088 | 0.03722 |
| d | g11 | Mw | 0.523 | 0.60215 | 0.080 | 0.93658 | 1.259 | 0.21116 |
| d | g12 | Sf | −2.754 | 0.00704 | −1.555 | 0.12327 | −1.775 | 0.07903 |
| d | g14 | Lb | 4.276 | 0.00004 | 1.367 | 0.17472 | 3.497 | 0.00071 |
| d | g15 | Sf | 1.275 | 0.20538 | −4.207 | 0.00006 | −2.510 | 0.01375 |
| d | g22 | Pm | 6.616 | <0.00001 | 2.758 | 0.00703 | 3.064 | 0.00288 |
| d | g23 | Dm | −2.636 | 0.00988 | −3.077 | 0.00276 | −3.414 | 0.00096 |
| d | g24 | Mw | 4.483 | 0.00002 | 0.053 | 0.95807 | 4.551 | 0.00002 |
| d | g25 | Sf | 2.321 | 0.02253 | −2.494 | 0.01442 | −0.870 | 0.38669 |
| d | g31 | Mw | 4.054 | 0.00011 | 2.386 | 0.01911 | 3.921 | 0.00017 |
| d | g32 | Sp | 0.887 | 0.37748 | −2.969 | 0.00382 | −3.847 | 0.00022 |
| d | g33 | Lb | −3.094 | 0.00263 | 3.015 | 0.00333 | 0.088 | 0.92986 |
| d | g34 | Dm | −0.543 | 0.58859 | 1.056 | 0.29367 | 0.456 | 0.64942 |
| d | g35 | Dm | 1.683 | 0.09573 | −3.223 | 0.00176 | −2.822 | 0.00586 |

See footnote to Table S1 for colour coding.

Appendix 5: Table S3. Tables of *t*-values and their probability levels for two time lags (0 and 1 days) for GLS-regression SMP coefficients (slopes) of daily stem girth increment versus soil moisture potential (SMP) and ambient temperature (TEMP). In these regression fits there were two independent terms, SMP and TEMP, but no interaction term. The coefficients and their standard errors are represented in Fig. S1 of Appendix 4.

| season | gth | spec | t(est)_00 | P(t)_00 | t(est)_01 | P(t)_01 | t(est)_10 | P(t)_10 | t(est)_11 | P(t)_11 |
|---|---|---|---|---|---|---|---|---|---|---|
| w | g11 | Mw | 5.294 | <0.00001 | 2.266 | 0.02385 | 7.854 | <0.00001 | 5.455 | <0.00001 |
| w | g12 | Sf | −1.285 | 0.19918 | 1.177 | 0.23985 | −4.237 | 0.00003 | −2.768 | 0.00583 |
| w | g14 | Lb | −0.048 | 0.96200 | 0.191 | 0.84900 | −0.114 | 0.90940 | −0.211 | 0.83340 |
| w | g15 | Sf | 4.582 | 0.00001 | 1.306 | 0.19205 | 8.188 | <0.00001 | 5.592 | <0.00001 |
| w | g22 | Pm | −5.996 | <0.00001 | −6.894 | <0.00001 | −6.124 | <0.00001 | −6.529 | <0.00001 |
| w | g23 | Dm | 1.854 | 0.06534 | −0.462 | 0.64440 | 3.665 | 0.00033 | 2.279 | 0.02386 |
| w | g24 | Mw | −1.124 | 0.26152 | −2.502 | 0.01264 | −0.957 | 0.33879 | −1.865 | 0.06270 |
| w | g25 | Sf | 4.331 | 0.00002 | 0.195 | 0.84545 | 7.759 | <0.00001 | 4.727 | <0.00001 |
| w | g31 | Mw | −2.463 | 0.01410 | −4.177 | 0.00003 | −2.648 | 0.00833 | −4.124 | 0.00004 |
| w | g32 | Sp | 0.151 | 0.88012 | −0.393 | 0.69457 | −0.231 | 0.81748 | −0.444 | 0.65720 |
| w | g33 | Lb | −3.518 | 0.00047 | −4.773 | <0.00001 | −4.083 | 0.00005 | −4.128 | 0.00004 |
| w | g34 | Dm | −3.172 | 0.00160 | −3.090 | 0.00210 | −4.340 | 0.00002 | −4.175 | 0.00003 |
| w | g35 | Dm | 2.199 | 0.02828 | −0.547 | 0.58483 | 5.873 | <0.00001 | 4.367 | 0.00002 |
| w | g41 | Sf | −1.603 | 0.11062 | −2.564 | 0.01117 | −1.892 | 0.06004 | −2.768 | 0.06228 |
| w | g42 | Dm | 6.455 | <0.00001 | 1.535 | 0.12525 | 9.109 | <0.00001 | 5.386 | <0.00001 |
| w | g43 | Pm | −5.237 | <0.00001 | −6.333 | <0.00001 | −5.396 | <0.00001 | −6.253 | <0.00001 |
| w | g44 | Sp | −4.834 | <0.00001 | −6.167 | <0.00001 | −8.113 | <0.00001 | −7.876 | <0.00001 |
| w | g45 | Pm | −4.288 | 0.00002 | −3.978 | 0.00008 | −5.435 | <0.00001 | −4.975 | <0.00001 |
| d | g11 | Mw | −1.075 | 0.28530 | −1.294 | 0.19871 | −0.688 | 0.49343 | −0.870 | 0.38666 |
| d | g12 | Sf | −0.874 | 0.38439 | 0.767 | 0.44491 | −1.271 | 0.20695 | −0.494 | 0.62216 |
| d | g14 | Lb | −1.809 | 0.07354 | −2.581 | 0.01139 | −1.893 | 0.06142 | −2.208 | 0.02966 |
| d | g15 | Sf | 2.234 | 0.02782 | 0.888 | 0.37695 | 5.460 | <0.00001 | 4.152 | 0.00007 |
| d | g22 | Pm | −0.539 | 0.59131 | −1.779 | 0.07864 | 0.727 | 0.46910 | −0.156 | 0.87676 |
| d | g23 | Dm | 2.803 | 0.00620 | 2.187 | 0.03137 | 3.587 | 0.00054 | 2.479 | 0.01502 |
| d | g24 | Mw | −2.823 | 0.00585 | −4.448 | 0.00002 | −2.347 | 0.02114 | −3.096 | 0.00261 |

| | | | | | | | | | | |
|---|---|---|---|---|---|---|---|---|---|---|
| d | g25 | Sf | 6.488 | <0.00001 | 3.111 | 0.00250 | 8.229 | <0.00001 | 5.208 | <0.00001 |
| d | g31 | Mw | −2.968 | 0.00384 | −3.818 | 0.00025 | −2.678 | 0.00881 | −3.349 | 0.00119 |
| d | g32 | Sp | 3.993 | 0.00013 | 2.970 | 0.00382 | 5.977 | <0.00001 | 5.132 | <0.00001 |
| d | g33 | Lb | −1.375 | 0.17262 | 0.549 | 0.58457 | −1.147 | 0.25429 | −0.505 | 0.61508 |
| d | g34 | Dm | −1.298 | 0.19750 | −1.404 | 0.16382 | −3.194 | 0.00194 | −2.995 | 0.00354 |
| d | g35 | Dm | 3.091 | 0.00266 | 0.567 | 0.57205 | 3.565 | 0.00059 | 1.782 | 0.07819 |

See footnote to Table S1 for colour coding.

Appendix 5: Table S4. Tables of *t*-values and their probability levels for two time lags (0 and 1 days) for GLS-regression TEMP coefficients (slopes) of daily stem girth increment versus soil moisture potential (SMP) and ambient temperature (TEMP). In these regression fits there were two independent terms, SMP and TEMP, but no interaction term. The coefficients and their standard errors are represented in Fig. S2 of Appendix 4.

| season | gth | spec | t(est)_00 | P(t)_00 | t(est)_01 | P(t)_01 | t(est)_10 | P(t)_10 | t(est)_11 | P(t)_11 |
|---|---|---|---|---|---|---|---|---|---|---|
| w | g11 | Mw | 4.724 | <0.00001 | −4.134 | 0.00004 | 4.506 | 0.00001 | −2.930 | 0.00353 |
| w | g12 | Sf | −5.816 | <0.00001 | 2.047 | 0.04115 | −6.492 | <0.00001 | 0.825 | 0.40975 |
| w | g14 | Lb | −1.022 | 0.30820 | 0.406 | 0.68560 | −1.078 | 0.28220 | 0.356 | 0.72200 |
| w | g15 | Sf | 6.226 | <0.00001 | −3.521 | 0.00047 | 6.734 | <0.00001 | −1.988 | 0.04732 |
| w | g22 | Pm | 3.898 | 0.00011 | 0.188 | 0.85083 | 4.973 | <0.00001 | 0.594 | 0.55255 |
| w | g23 | Dm | 5.649 | 0.00001 | −2.792 | 0.00581 | 5.600 | 0.00001 | −1.928 | 0.05544 |
| w | g24 | Mw | 3.609 | 0.00034 | −0.089 | 0.92889 | 4.004 | 0.00007 | 0.216 | 0.82887 |
| w | g25 | Sf | 7.500 | <0.00001 | −4.947 | <0.00001 | 7.798 | <0.00001 | −3.269 | 0.00115 |
| w | g31 | Mw | 3.249 | 0.00123 | −1.199 | 0.23110 | 3.465 | 0.00057 | −0.966 | 0.33458 |
| w | g32 | Sp | 1.463 | 0.14414 | −0.492 | 0.62285 | 1.483 | 0.13871 | −0.532 | 0.59465 |
| w | g33 | Lb | 5.128 | <0.00001 | −0.235 | 0.81416 | 6.101 | <0.00001 | 0.104 | 0.91757 |
| w | g34 | Dm | −0.493 | 0.62233 | −0.267 | 0.78925 | −0.555 | 0.57918 | −0.609 | 0.54247 |
| w | g35 | Dm | 6.486 | <0.00001 | −0.916 | 0.35999 | 7.309 | <0.00001 | 1.023 | 0.30668 |
| w | g41 | Sf | 2.012 | 0.04570 | −1.094 | 0.27540 | 2.339 | 0.02042 | −1.006 | 0.31600 |
| w | g42 | Dm | 8.920 | <0.00001 | −3.594 | 0.00036 | 9.059 | <0.00001 | −2.295 | 0.02209 |
| w | g43 | Pm | 1.134 | 0.25745 | −2.366 | 0.01834 | 1.192 | 0.23372 | −2.321 | 0.02068 |
| w | g44 | Sp | 0.202 | 0.84038 | 0.399 | 0.69014 | 1.164 | 0.24511 | 0.155 | 0.87703 |
| w | g45 | Pm | −0.450 | 0.65270 | 0.544 | 0.58645 | −0.417 | 0.67648 | 0.151 | 0.88026 |
| d | g11 | Mw | −0.053 | 0.95764 | −0.551 | 0.58265 | 0.229 | 0.81970 | −0.298 | 0.76669 |
| d | g12 | Sf | −2.869 | 0.00507 | −1.134 | 0.25976 | −3.034 | 0.00321 | −1.592 | 0.11464 |
| d | g14 | Lb | 3.124 | 0.00236 | 0.675 | 0.50103 | 3.648 | 0.00043 | 0.411 | 0.68173 |
| d | g15 | Sf | 1.739 | 0.08536 | −3.701 | 0.00036 | 1.860 | 0.06603 | −2.858 | 0.00525 |
| d | g22 | Pm | 6.041 | <0.00001 | 2.078 | 0.04053 | 6.586 | <0.00001 | 2.324 | 0.02239 |
| d | g23 | Dm | −0.465 | 0.64331 | −2.027 | 0.04557 | −1.762 | 0.08144 | −2.075 | 0.04080 |
| d | g24 | Mw | 2.801 | 0.00624 | −2.613 | 0.01051 | 3.715 | 0.00035 | 1.170 | 0.24501 |

| | | | | | | | | | | |
|---|---|---|---|---|---|---|---|---|---|---|
| d | g25 | Sf | 1.894 | 0.06150 | –1.939 | 0.05562 | 1.824 | 0.07139 | –1.733 | 0.08654 |
| d | g31 | Mw | 2.834 | 0.00568 | 0.427 | 0.67064 | 3.196 | 0.00193 | 1.003 | 0.31858 |
| d | g32 | Sp | 0.959 | 0.34034 | –2.274 | 0.02533 | 0.701 | 0.48514 | –1.757 | 0.08231 |
| d | g33 | Lb | –3.344 | 0.00120 | 2.927 | 0.00434 | –3.167 | 0.00210 | 2.769 | 0.00683 |
| d | g34 | Dm | –0.769 | 0.44409 | 0.938 | 0.35094 | –0.618 | 0.53787 | 0.111 | 0.91214 |
| d | g35 | Dm | 2.903 | 0.00464 | –2.516 | 0.01365 | 2.611 | 0.01058 | –2.163 | 0.03320 |

See footnote to Table S1 for colour coding.

Appendix 5: Table 5A. Tables of *t*–values and their probability levels for two time lags (0 and 1 days) for GLS–regression SMP*TEMP interaction of daily stem girth increment, and their coefficients and standard errors, versus soil moisture potential (SMP) and ambient temperature (TEMP). In these regression fits there were two independent terms, SMP and TEMP, plus the interaction term. Here SMP had no lag and TEMP had either no lag (0, 0) or was lagged by 1 day also (0, 1). Coefficients of SMP and TEMP for this particular regression model are not reported in the paper.

| season | gth | spec | est_00 | se_00 | t(est)_00 | P(t)_00 | est_01 | se_01 | t(est)_01 | P(t)_01 |
|---|---|---|---|---|---|---|---|---|---|---|
| w | g11 | Mw | −0.1426 | 0.0550 | −2.5912 | 0.00982 | 0.0352 | 0.0583 | 0.6040 | 0.54611 |
| w | g12 | Sf | 0.0475 | 0.0819 | 0.5807 | 0.56166 | −0.0060 | 0.0861 | −0.0696 | 0.94455 |
| w | g14 | Lb | 0.0679 | 0.8327 | 0.5896 | 0.55600 | 0.2917 | 0.9048 | 1.1254 | 0.26200 |
| w | g15 | Sf | −0.0559 | 0.0589 | −0.9505 | 0.34226 | 0.0245 | 0.0603 | 0.4064 | 0.68459 |
| w | g22 | Pm | −0.4455 | 0.1877 | −2.3734 | 0.01805 | −0.2212 | 0.2029 | −1.0898 | 0.27640 |
| w | g23 | Dm | −0.0215 | 0.4146 | −0.1000 | 0.92040 | −0.2266 | 0.4463 | −0.5366 | 0.59220 |
| w | g24 | Mw | −0.0283 | 0.0311 | −0.9094 | 0.36354 | 0.0215 | 0.0329 | 0.6512 | 0.51522 |
| w | g25 | Sf | −0.1791 | 0.0667 | −2.6839 | 0.00750 | 0.0231 | 0.0722 | 0.3201 | 0.74903 |
| w | g31 | Mw | −0.0533 | 0.0228 | −2.3435 | 0.01946 | 0.0528 | 0.0257 | 2.0559 | 0.04026 |
| w | g32 | Sp | −0.0327 | 0.0311 | −1.0499 | 0.29425 | 0.0227 | 0.0341 | 0.6644 | 0.50674 |
| w | g33 | Lb | −0.0405 | 0.0200 | −2.0230 | 0.04356 | 0.0167 | 0.0226 | 0.7375 | 0.46112 |
| w | g34 | Dm | −0.0387 | 0.0274 | −1.4091 | 0.15938 | 0.0296 | 0.0303 | 0.9763 | 0.32936 |
| w | g35 | Dm | −0.0608 | 0.0519 | −1.1719 | 0.24176 | 0.0354 | 0.0573 | 0.6186 | 0.53644 |
| w | g41 | Sf | −0.1200 | 0.1053 | −0.0819 | 0.93480 | −0.0501 | 0.1050 | 0.1927 | 0.84740 |
| w | g42 | Dm | −0.0167 | 0.0197 | −0.8472 | 0.39725 | 0.0009 | 0.0202 | 0.0463 | 0.96310 |
| w | g43 | Pm | −0.0420 | 0.0825 | −0.5090 | 0.61099 | 0.0588 | 0.0897 | 0.6550 | 0.51273 |
| w | g44 | Sp | −0.0217 | 0.0347 | −0.6263 | 0.53135 | 0.0662 | 0.0538 | 1.2306 | 0.21899 |
| w | g45 | Pm | −0.0221 | 0.0361 | −0.6116 | 0.54107 | 0.0016 | 0.0301 | 0.0540 | 0.95697 |
| d | g11 | Mw | 0.0175 | 0.0141 | 1.237 | 0.21931 | 0.0171 | 0.0141 | 1.215 | 0.22732 |
| d | g12 | Sf | −0.0107 | 0.0076 | −1.405 | 0.16320 | −0.0093 | 0.0081 | −1.158 | 0.24972 |
| d | g14 | Lb | −0.0112 | 0.0173 | −0.648 | 0.51862 | −0.0183 | 0.0150 | −1.219 | 0.22608 |
| d | g15 | Sf | 0.1079 | 0.0174 | 6.198 | <0.00001 | 0.1007 | 0.0167 | 6.024 | <0.00001 |
| d | g22 | Pm | 0.0293 | 0.0126 | 2.319 | 0.02269 | 0.0542 | 0.0127 | 4.263 | 0.00005 |
| d | g23 | Dm | 0.0371 | 0.0088 | 4.210 | 0.00006 | 0.0158 | 0.0108 | 1.458 | 0.14828 |

| | | | | | | | | | | |
|---|---|---|---|---|---|---|---|---|---|---|
| d | g24 | Mw | 0.0078 | 0.0106 | 0.740 | 0.46143 | 0.0056 | 0.0101 | 0.549 | 0.58450 |
| d | g25 | Sf | 0.0585 | 0.0173 | 3.388 | 0.00105 | 0.0569 | 0.0171 | 3.335 | 0.00125 |
| d | g31 | Mw | 0.0162 | 0.0251 | 0.645 | 0.52034 | 0.0019 | 0.0268 | 0.070 | 0.94426 |
| d | g32 | Sp | 0.0390 | 0.0075 | 5.175 | <0.00001 | 0.0381 | 0.0073 | 5.204 | <0.00001 |
| d | g33 | Lb | −0.0044 | 0.0114 | −0.384 | 0.70216 | −0.0058 | 0.0116 | −0.502 | 0.61720 |
| d | g34 | Dm | −0.0254 | 0.0068 | −3.757 | 0.00031 | −0.0249 | 0.0068 | −3.677 | 0.00040 |
| d | g35 | Dm | 0.0199 | 0.0105 | 1.905 | 0.06002 | 0.0232 | 0.0106 | 2.180 | 0.03188 |

See footnote to Table S1 for colour coding.

Appendix 5: Table 5B. Tables of t-values and their probability levels for two time lags (0 and 1 days) for GLS−regression SMP*TEMP interaction coefficients of daily stem girth increment, and their coefficients and standard errors, versus soil moisture potential (SMP) and ambient temperature (TEMP). In these regression fits there were two independent terms, SMP and TEMP, plus the interaction term. Here SMP had a lag of 1 day and TEMP had either no lag (1, 0) or was lagged by 1 day also (1, 1). Coefficients of SMP and TEMP for this particular regression model are not reported in the paper.

| season | gth | spec | est_10 | se_10 | t(est)_10 | P(t)_10 | est_11 | se_11 | t(est)_11 | P(t)_11 |
|---|---|---|---|---|---|---|---|---|---|---|
| w | g11 | Mw | −0.1001 | 0.0455 | −2.2023 | 0.02806 | 0.1089 | 0.0426 | 2.5544 | 0.01091 |
| w | g12 | Sf | 0.0356 | 0.0510 | 0.6983 | 0.48528 | −0.0373 | 0.0615 | −0.6069 | 0.54417 |
| w | g14 | Lb | −0.2908 | 0.8124 | −0.3634 | 0.71680 | 0.2488 | 0.8466 | 1.0011 | 0.31820 |
| w | g15 | Sf | −0.0043 | 0.0198 | −0.2150 | 0.82987 | 0.0358 | 0.0318 | 1.1239 | 0.26155 |
| w | g22 | Pm | −0.3796 | 0.1787 | −2.1244 | 0.03419 | −0.3102 | 0.1864 | −1.6639 | 0.09683 |
| w | g23 | Dm | −0.1353 | 0.4142 | −0.7092 | 0.47920 | −0.1500 | 0.4217 | −0.2582 | 0.79660 |
| w | g24 | Mw | −0.0294 | 0.0310 | −0.9479 | 0.34360 | 0.0106 | 0.0300 | 0.3517 | 0.72520 |
| w | g25 | Sf | −0.2031 | 0.0728 | −2.7914 | 0.00543 | 0.1647 | 0.0672 | 2.4490 | 0.01464 |
| w | g31 | Mw | −0.0453 | 0.0245 | −1.8510 | 0.06472 | 0.0341 | 0.0197 | 1.7290 | 0.08437 |
| w | g32 | Sp | −0.0202 | 0.0275 | −0.7352 | 0.46255 | 0.0457 | 0.0117 | 3.9192 | 0.00010 |
| w | g33 | Lb | −0.0154 | 0.0054 | −2.8290 | 0.00484 | 0.0168 | 0.0065 | 2.5644 | 0.01060 |
| w | g34 | Dm | −0.0288 | 0.0216 | −1.3324 | 0.18329 | 0.0349 | 0.0229 | 1.5218 | 0.12863 |
| w | g35 | Dm | −0.0784 | 0.0720 | −1.0893 | 0.27652 | 0.0674 | 0.0709 | 0.9501 | 0.34248 |
| w | g41 | Sf | −0.0939 | 0.0972 | −0.1452 | 0.88480 | −0.0404 | 0.0930 | 0.2820 | 0.77820 |
| w | g42 | Dm | −0.0262 | 0.0186 | −1.4051 | 0.16055 | 0.0152 | 0.0191 | 0.7976 | 0.42547 |
| w | g43 | Pm | 0.0015 | 0.0252 | 0.0586 | 0.95331 | −0.0179 | 0.0236 | −0.7578 | 0.44892 |
| w | g44 | Sp | 0.0005 | 0.0085 | 0.0564 | 0.95503 | −0.0040 | 0.0097 | −0.4079 | 0.68347 |
| w | g45 | Pm | −0.0023 | 0.0098 | −0.2307 | 0.81763 | −0.0029 | 0.0112 | −0.2602 | 0.79483 |
| d | g11 | Mw | 0.0292 | 0.0856 | 0.341 | 0.73414 | −0.0044 | 0.0345 | −0.127 | 0.89938 |
| d | g12 | Sf | −0.0202 | 0.0463 | −0.435 | 0.66448 | 0.0355 | 0.0192 | 1.848 | 0.06779 |
| d | g14 | Lb | −0.0587 | 0.1160 | −0.506 | 0.61422 | −0.1545 | 0.0428 | −3.610 | 0.00049 |
| d | g15 | Sf | −0.4838 | 0.1077 | −4.490 | 0.00002 | −0.1305 | 0.0427 | −3.053 | 0.00295 |
| d | g22 | Pm | −0.0227 | 0.0799 | −0.285 | 0.77645 | −0.2042 | 0.0330 | −6.197 | <0.00001 |
| d | g23 | Dm | −0.1528 | 0.0665 | −2.299 | 0.02383 | 0.0187 | 0.0262 | 0.714 | 0.47679 |

| | | | | | | | | | | |
|---|---|---|---|---|---|---|---|---|---|---|
| d | g24 | Mw | −0.1614 | 0.0744 | −2.169 | 0.03271 | −0.1123 | 0.0298 | −3.774 | 0.00029 |
| d | g25 | Sf | −0.2257 | 0.1096 | −2.059 | 0.04243 | −0.0960 | 0.0420 | −2.287 | 0.02457 |
| d | g31 | Mw | −0.1181 | 0.1628 | −0.725 | 0.47019 | −0.2132 | 0.0631 | −3.378 | 0.00108 |
| d | g32 | Sp | −0.1820 | 0.0489 | −3.720 | 0.00035 | −0.0422 | 0.0182 | −2.316 | 0.02287 |
| d | g33 | Lb | −0.1329 | 0.0695 | −1.913 | 0.05897 | 0.0616 | 0.0260 | 2.364 | 0.02026 |
| d | g34 | Dm | 0.0586 | 0.0453 | 1.293 | 0.19947 | 0.0203 | 0.0164 | 1.238 | 0.21910 |
| d | g35 | Dm | 0.0318 | 0.0705 | 0.451 | 0.65329 | −0.0881 | 0.0267 | −3.299 | 0.00140 |

See footnote to Table S1 for color coding.

Appendix 6: Figure S1. Mean ± SE coefficients (slopes) for SMP of *gthi* regressed on soil moisture potential, SMP, at 0- and 1-day lags, with logger temperature, TEMP, at 0- and 1-day lags, in the wet (green bars) and dry (orange bars) periods for the 18 trees taken for the time-series analyses. Predictions of dry-period estimates for station 4 (see text for explanation, and Appendix 1: Table S5) are shown in blue. Bands are listed in Table 4 (main text) with their individual tree codes.

Appendix 6: Figure S2. Mean ± SE coefficients (slopes) for TEMP of *gthi* regressed on soil moisture potential, SMP, at 0- and 1-day lags, with logger temperature, TEMP, at 0- and 1-day lags, in the wet (green bars) and dry (orange bars) periods for the 18 trees taken for the time-series analyses. Predictions of dry-period estimates for station 4 (see text for explanation, and Appendix 1: Table S5) are shown in blue. Bands are listed in Table 4 (main text) with their individual tree codes.

App.6 Fig.S1 − page 1

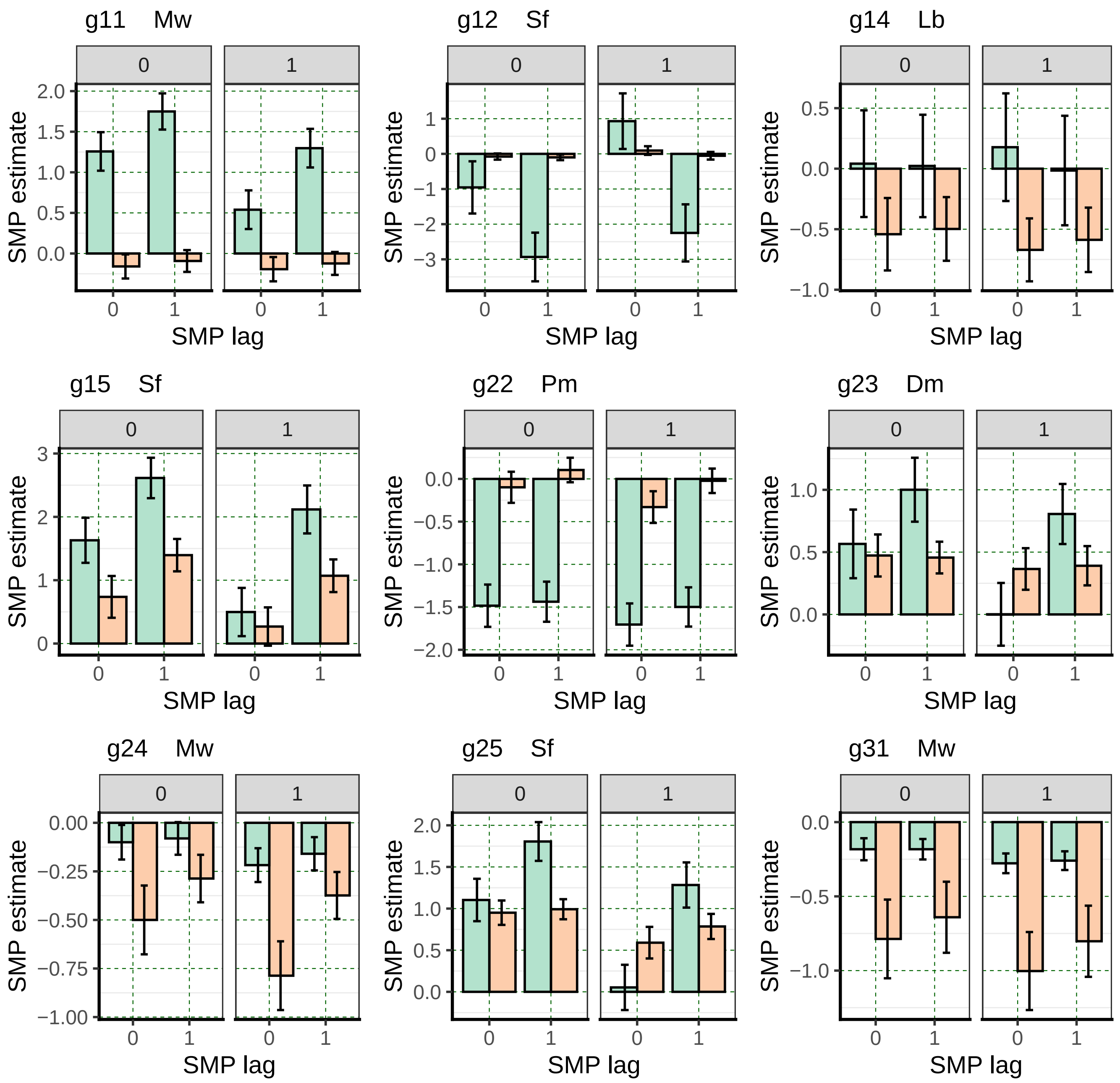

App.6 Fig.S1 − page 2

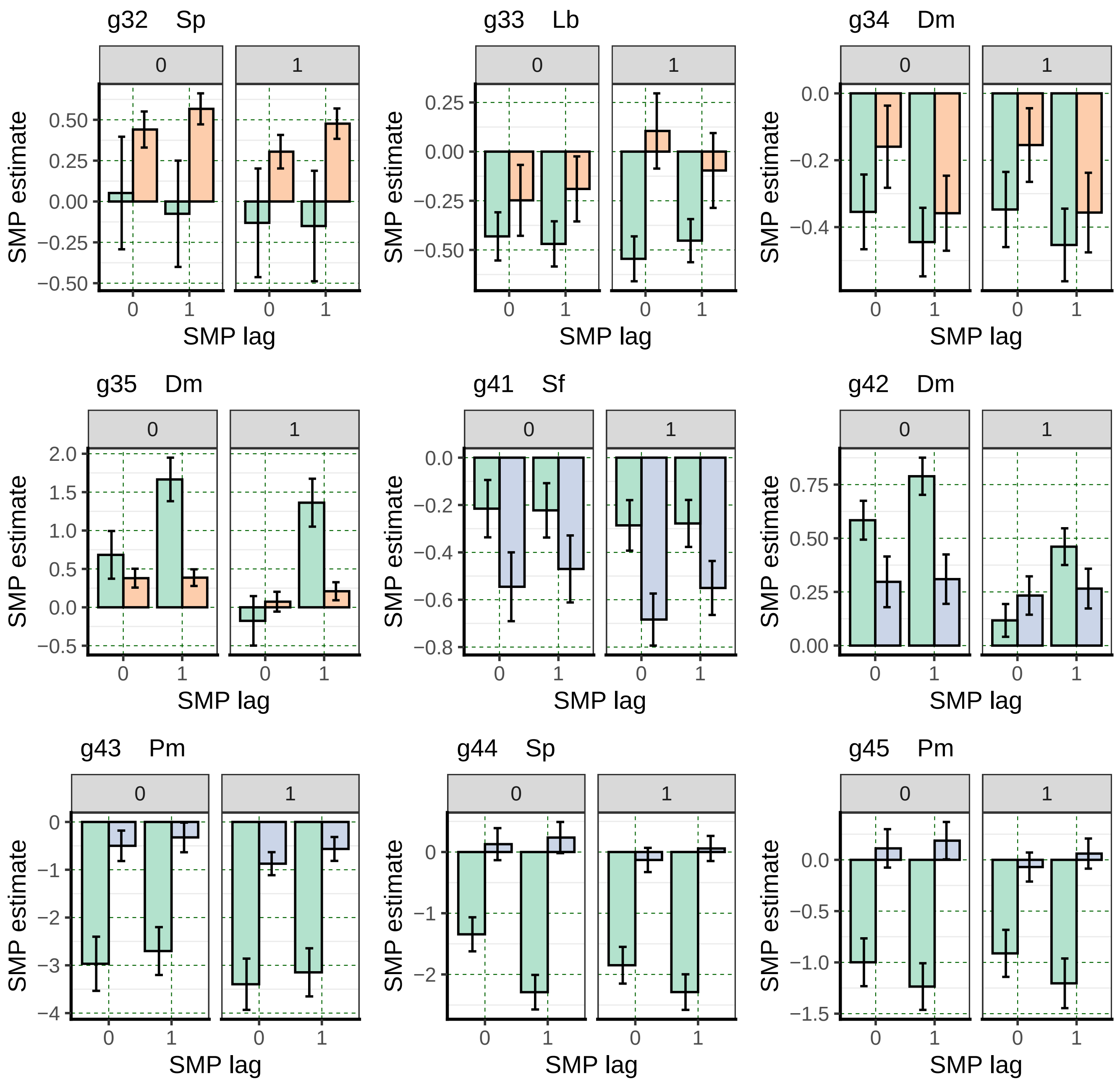

App.6 Fig.S2 − page 1

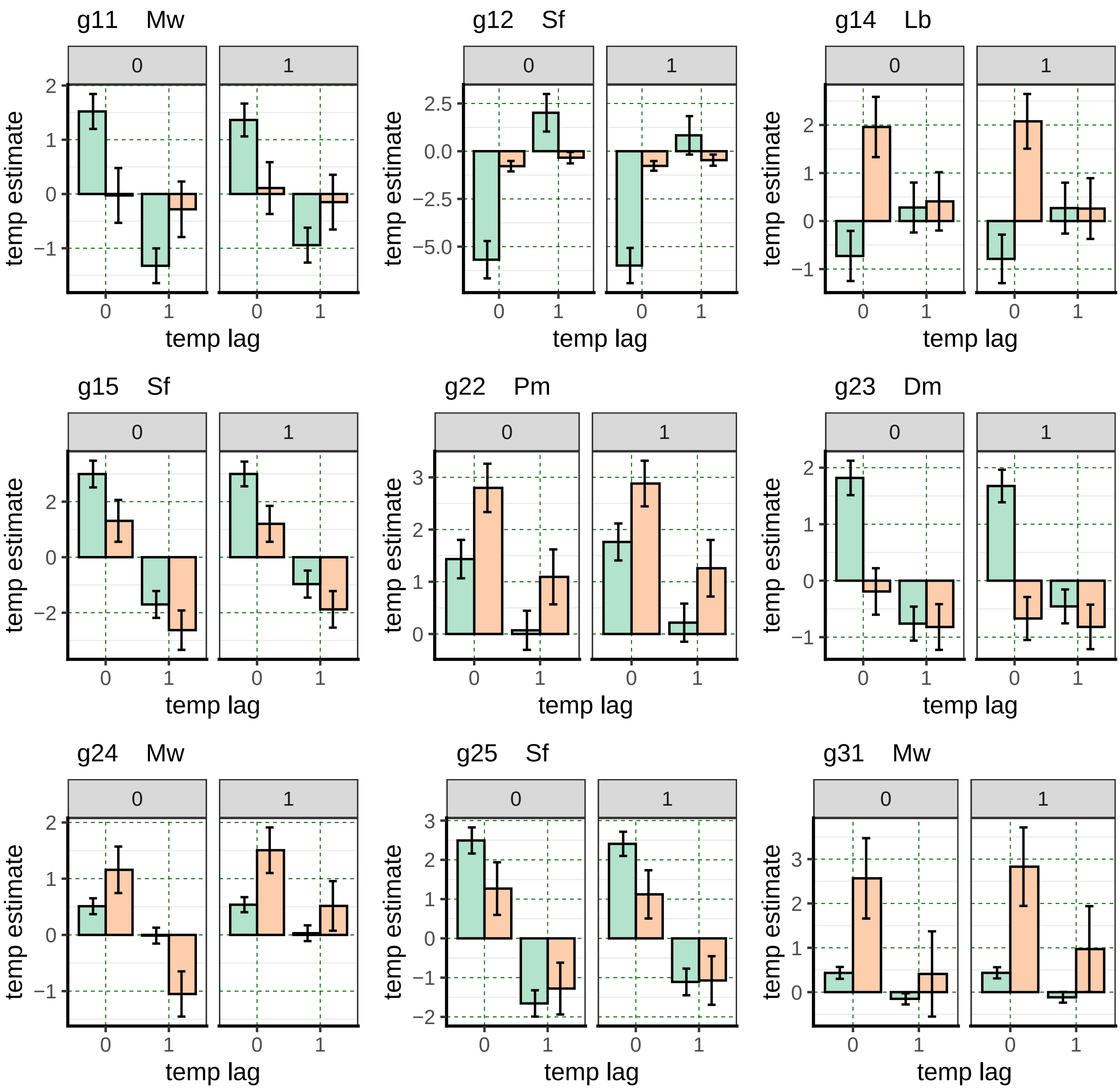

App.6 Fig.S2 − page 2

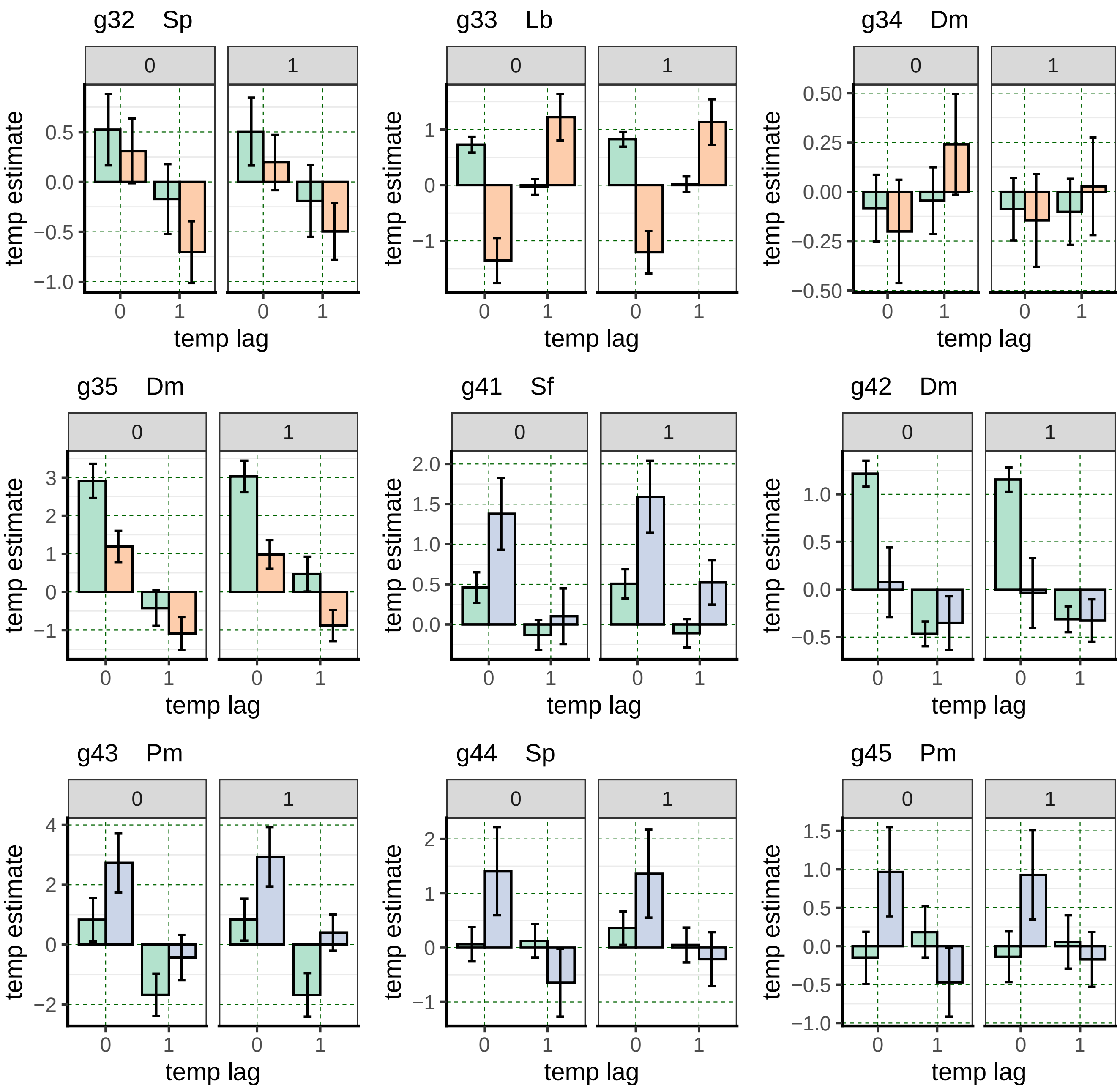
